# Transcriptomic Analysis of Early Stages of Intestinal Regeneration in *Holothuria glaberrima*

**DOI:** 10.1101/2020.09.23.310599

**Authors:** David J. Quispe-Parra, Joshua G. Medina-Feliciano, Sebastián Cruz-González, Humberto Ortiz-Zuazaga, José E. García-Arrarás

## Abstract

Echinoderms comprise a group of animals with impressive regenerative capabilities. They can replace complex internal organs following injury or autotomy. In holothurians or sea cucumbers, cellular processes of intestinal regeneration have been extensively studied. The molecular machinery behind this faculty, however, remains to be understood. Here we assembled and annotated a de novo transcriptome using RNA-seq data consisting of regenerating and non-regenerating intestinal tissues from the sea cucumber *Holothuria glaberrima*. Comparisons of differential expression were made using the mesentery as a reference against 24 hour and 3 days regenerating intestine, revealing a large number of differentially expressed transcripts. Gene ontology and pathway enrichment analysis showed evidence of increasing transcriptional activity. Further analysis of transcripts associated with transcription factors revealed diverse expression patterns with mechanisms involving developmental and cancer-related activity that could be related to the regenerative process. Our study demonstrates the broad and diversified gene expression profile during the early stages of the process using the mesentery as the focal point of intestinal regeneration. It also establishes the genes that are the most important candidates in the cellular processes that underlie regenerative responses.

## Introduction

Regeneration is fundamental for renewal of cells and tissues in multicellular organisms, nevertheless the process of regenerating complex components, such as organs, is only possible in a few animal groups^1^. Understanding the limitations of this capability has become an important goal in improving health conditions related to tissue and organ functional maintenance^2^. However, discerning this process requires a clear understanding of the molecular mechanisms that drive the cellular functions responsible for regeneration.

Organisms with astonishing regenerative capacities are scattered among basal metazoan lineages and in various lophotrochozoan and deuterostome phyla. The latter encompasses a wide group of organisms that includes vertebrates along with tunicates, echinoderms, hemichordates, and cephalochordates. This phylogenetic placement makes highly regenerative groups such as tunicates and echinoderms, attractive models for studying regeneration. Their potentially shared genetic mechanisms highlight their importance for biomedical studies^3,4^.

Echinoderms comprise an important group of organisms with extraordinary regeneration capabilities. However, despite being among the most highly regenerative deuterostomes and their close relationship to vertebrates, the number of studies on their regenerative capacity is rather limited^5^. In recent years their use as regeneration model organisms has seen an upsurge with studies on spine regeneration in sea urchins^6^, arm regeneration in crinoids, brittle stars and sea stars^7^,^8,9^ and nervous system and viscera regeneration in sea cucumbers^10,11,12,13,14^. Sea cucumbers have been particularly attractive for intestinal regeneration studies because of their innate capacity of expelling their digestive tract in a process known as evisceration, which is then followed by its regeneration. Research relating regeneration of the whole digestive system is scarce among regenerating competent organisms, where in addition to sea cucumbers, other studies include planarians^15,16^.

Efforts in understanding the regeneration of the holothurian digestive tract have provided important insights on the cellular processes involved in organogenesis. Experiments have identified the fundamental role of the cells in the mesentery during intestinal regeneration^17^. In normal, non-eviscerated animals, the mesentery extends along the interior body wall providing support to the intestine. Figure 1 shows a simplified drawing of the tissues found in the mesentery and intestine. This includes (1) the outer mesothelium made of peritoneocytes (or coelomic epithelium), myocytes and some neurons, (2) the inner connective tissue layer where few cells are found within an abundant extracellular matrix, and (3) the luminal epithelium made up primarily of enterocytes and some enteroendocrine cells. Following evisceration, the torn edges at the distal part of the mesentery acquire a thick oval morphology forming the initial intestinal rudiment from the esophagus to the cloaca. This growth is formed by evisceration-activated signals that trigger dedifferentiation of peritoneocytes and muscle cells in the mesothelium of the mesentery. Myocyte dedifferentiation is associated with elimination of the contractile apparatus through condensation of myofilaments into membrane bound spindle-like structures (SLSs)^18^. The presence of these structures has been used to reveal a spatio-temporal gradient where, initially, dedifferentiation is localized near the free margin of the mesentery, while the rest of the mesentery remains mostly in a differentiated state. This pattern progresses until eventually different regions with increasing levels of de-differentiation can be observed in the mesentery. It has been hypothesized that dedifferentiated cells give rise to precursor cells that eventually divide and re-differentiate into the cellular components that form the initial rudiment of the regenerating organ^19^.

**Figure 1.**
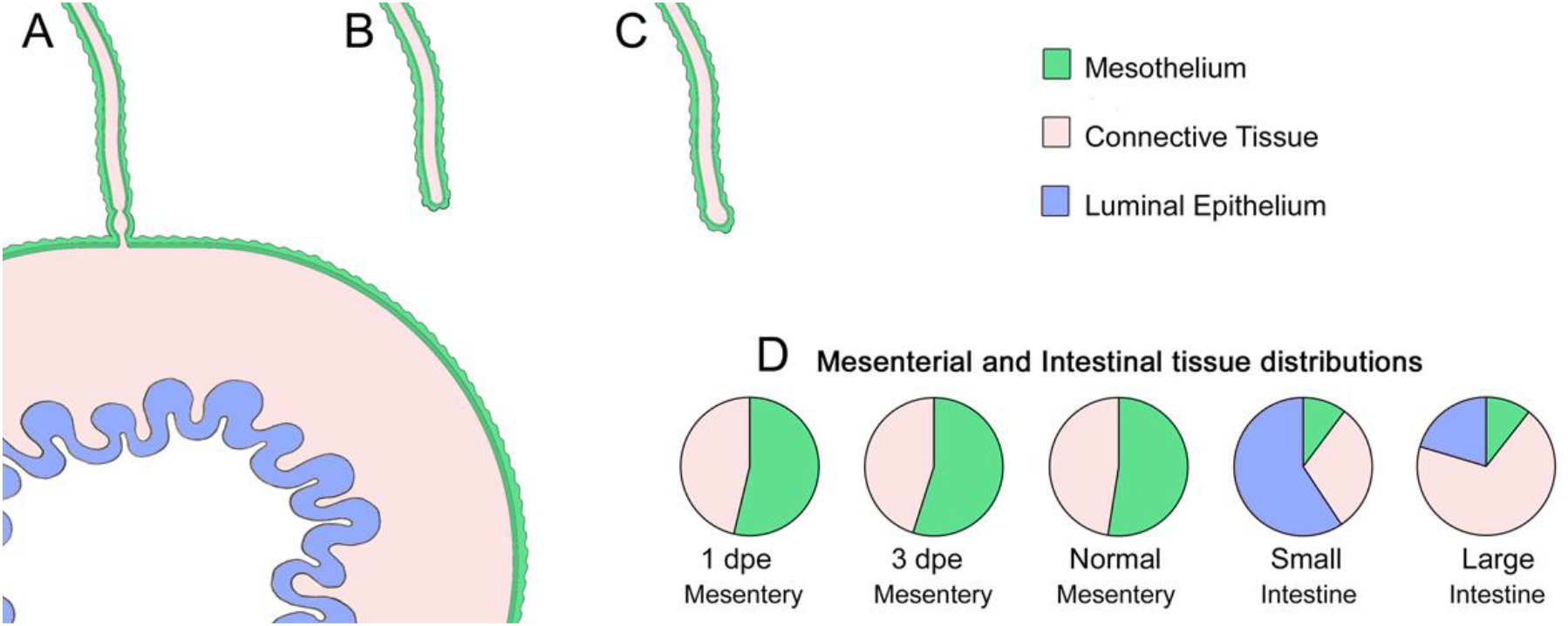
Diagram illustrating the intestine composition of normal and regenerating stages in *H. glaberrima*. (A) Normal intestine and attached mesentery. The other end of the mesentery (not shown) is attached to the body wall. (B) Mesentery one day after evisceration of the intestine. (C) Mesentery three days following evisceration Distribution of tissues in normal intestine and mesentery and in regenerating mesentery/intestine at 1 and 3 days post-evisceration (dpe).

In subsequent stages, mesothelial cells within the regenerating structure undergo an epithelial to mesenchymal transition (EMT) and ingress into the underlying connective tissue^14^. The increasing number of cells in the distal part of the mesentery together with the possible deposition of new extracellular matrix (ECM), contribute to the rudiment growth. Therefore, about a week after evisceration, the intestinal rudiment can be clearly observed as a solid rod at the free margin of the mesentery. In histological sections the rudiment appears as an elongated thickening of the mesentery with many mesenchymal cells in the inner connective tissue surrounded by a dedifferentiated coelomic epithelium. Within two weeks, the lumen forms and the reconstitution of mucosal epithelium is achieved by the intrusion of tubular projections of the mucosal epithelium that migrate from anterior and posterior ends. It is important to highlight the role of the mesentery during all the intestinal regeneration process, since new cells originate from this structure as well as the extracellular signals that modulate the whole process^17^.

Although substantial advances have been achieved in dissecting the cellular mechanisms of intestinal regeneration, a similar analysis of the molecular basis is lagging. Initial methods to identify differentially expressed genes (DEGs) during intestinal regeneration consisted of differential screening (DS) and differential display (DD) to study specific genes that could be upregulated during regeneration^20,21,22^. Further studies with a broader approach targeted expressed sequence tags (ESTs) from cDNA libraries of regenerating intestines and the preparation of microarrays to reveal DEGs. In the last decade, these technologies have been used to uncover regeneration-associated genes in the transcriptomes of two other sea cucumbers, *Apostichopus japonicus*^23^ and *Eupentacta fraudatrix*^24^. This has led to the identification of several DEGs linked with intestinal regeneration.

All previous intestinal comparative gene studies, however, share a major problem, described by Boyko as “the artificiality of comparing gene expression levels in regenerating and intact tissues”^24^. Many studies end up using the normal intestine as a baseline in their comparisons to determine the differences in gene expression profiles. This comparison is apt for late regenerative stages, but during the early events of regeneration, the regenerating rudiment shares more cell and tissue types in common with the normal mesentery than with the normal intestine. This point is highlighted in Figure 1, where the percentage composition of the different tissue layers is shown for the early stages of regeneration. Therefore, when using the normal intestine as a reference, genes expressed in the intestine luminal epithelium would appear as being under-expressed during the first week of regeneration since at that stage no lumen has formed. In contrast, genes that appeared as highly expressed in the early regenerate might be due to their preferential expression in the mesothelial layer and not necessarily due to being activated by the regeneration process.

We have now applied RNA-seq technology to the analysis of intestinal regeneration in our model system, the sea cucumber *Holothuria glaberrima*. More importantly, to narrow down the genes involved in the early steps of the regenerative process we made two critical modifications in our experimental strategy. First, we focused on tissues at day 1 and day 3 of regeneration, stages that take place prior to, or concomitant with, the first observed cellular events. Second, regenerating tissues were compared to normal, uneviscerated mesentery. By using this approach, we obtained a real view concerning temporal differential gene expression during early stages of intestinal regeneration. Moreover, we were able to obtain a considerable amount of information regarding novel transcripts that appear to be activated during regeneration.

## Results

Our results highlight what was found by comparing the mesentery at three different stages of regeneration. The normal, uninjured mesentery is mainly composed of mesothelium (peritoneocytes, muscles and neurons) and connective tissue. The regenerating mesentery at 1 and 3 days after evisceration is made of the same tissues, but have undergone some changes in the differentiated state of its cells, and, as regeneration advances more undifferentiated cells are found in the area where the new intestine will form.

Paired end reads obtained from the sequencing process had a satisfactory quality level after trimming. A phred score of 30 was obtained for all samples while N content was practically null in all samples as well (see Supplementary Fig. S1 online). De novo assembly was generated using reads from this study and previous raw sequencing reads from the central nervous system of *H. glaberrima*^25^. The resulting transcriptome with 490 172 contigs was assessed with BUSCO and showed 99.1% of complete core genes detected with 0.4% of fragmented genes and only 0.5% of missing core genes (see Supplementary Table S1 online). Furthermore, the N50 sequence length was 1692 nt with the longest and shortest sequence being 34610 nt and 200 nt respectively (see Supplementary Table S2 online).

### Differential expression analysis

Mapped reads against the reference transcriptome encompassed >90% of the total reads used for the analysis (see supplementary Table S3 online). A total of 8460 differentially expressed transcripts (DETs) were found significant (adjusted P-value ≤ 0.05) at day 1 of regeneration (Fig. 2A). Upregulated transcripts (Log2 Fold Change ≥ 2) comprised 3585, while downregulated (Log2 Fold Change ≤ -2) were 4875. On day 3 of regeneration, the total number of transcripts with differential expression was 8216, of these 3663 were upregulated and 4553 downregulated (Fig. 2B). When both days 1 and 3 regenerating intestines were compared to controls, a total of 3884 transcripts were differentially co-expressed on both days; 1723 were upregulated and 2161 downregulated (Fig. 2C).

**Figure 2.**
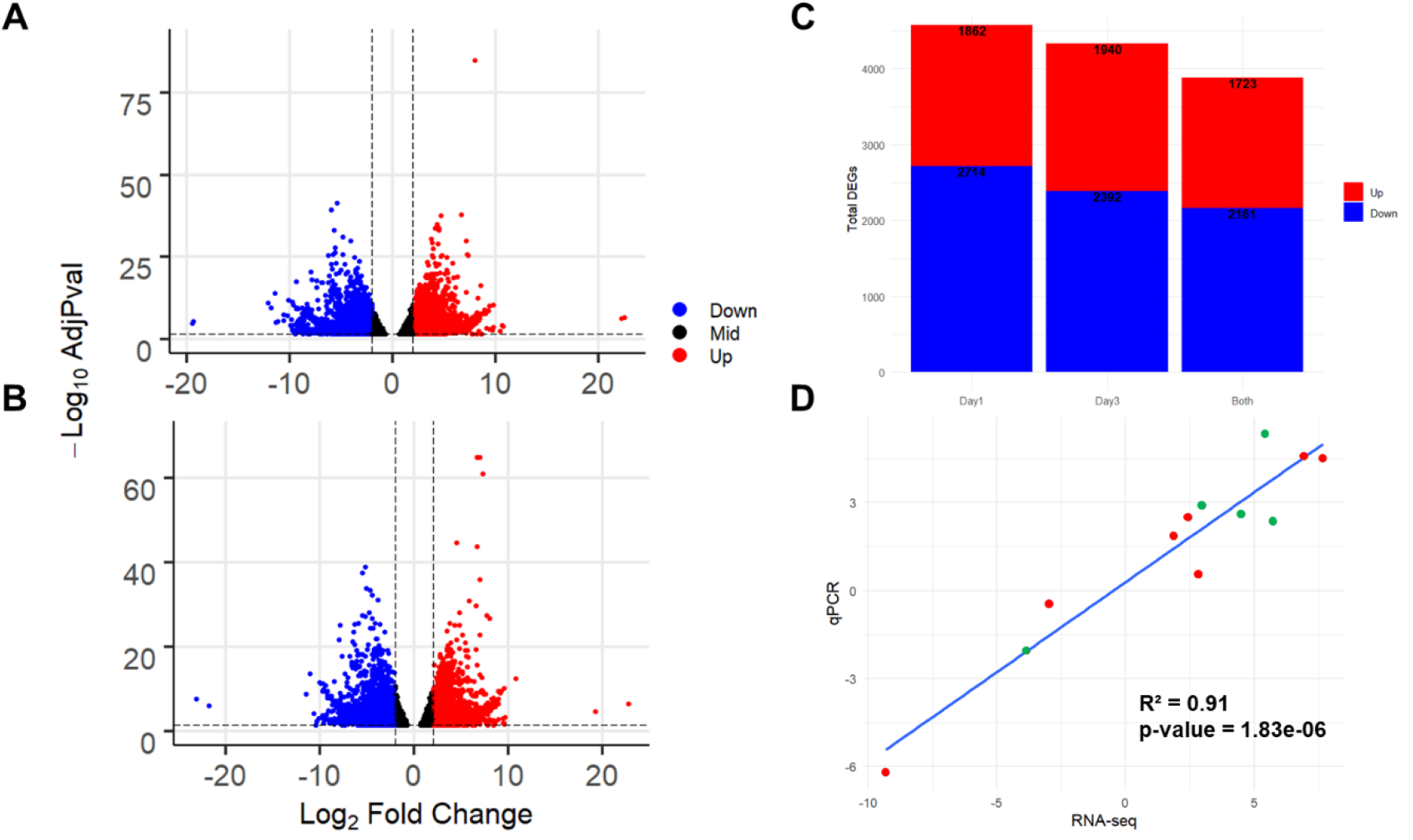
Gene expression patterns of RNA-seq and correlation with RT-qPCR. (A) Volcano plot of differentially expressed transcripts (DETs) at day 1 of regeneration. (B) Volcano plot of DETs at day 3 of regeneration. (C) Bar plots show upregulated and downregulated DETs in one or both days of regeneration. (D) Plot showing correlation line, squared Pearson correlation coefficient and p-value between 12 expression values (Log2FC) of seven genes in regeneration stages compared to the normal stage. RNA-seq values are located in the x axis and RT-qPCR values are located on the y axis. Green colored points are from day 1 values, while red colored are from day 3.

Comparisons were made with previous studies where the same genes had been analyzed using semiquantitative RT-PCR, RT-qPCR or *in situ* hybridization (see supplementary Table S6 online). These transcripts correspond to annotated genes Myc, Serum amyloid A, Mtf, WNT9, BMP, β-catenin and survivin, and all of them shared the fact that their expression in the early intestinal rudiment (1-3 dpe) had been compared to that of the normal mesentery^20,26,27,28,29^. Our RNA-seq expression analysis for all these genes showed a similar expression pattern to their expression in prior studies. Thus, Myc, Serum amyloid A, Mtf, WNT9, BMP, were over-expressed while β-catenin and survivin showed no considerable difference between regenerating intestine and normal mesentery. To further validate our data, we selected 7 additional genes that were differentially expressed in our RNAseq and probed their expression using RT-qPCR. The analyzed upregulated genes were VBP, Wnt6, Sox4, TAP26 and Mortalin. On the other hand, downregulated genes included FoxA and Tenascin. The expression values were compared between the RNA-seq and RT-qPCR expression data. In all cases the data from both day 1 and 3 were used, with the exemption of Tenascin and Mortalin with only the day 3 values. The comparison showed a high correlation between both results with a square of the Pearson’s correlation coefficient of 0.91 (Fig. 2D).

A wide range of annotated transcripts with differential expression were designated as DEGs. Here we provide a list of the top 10 upregulated and downregulated DEGs for days 1 and 3 as examples of some of the genes in our database (Tables 1-4). The ID column is the top hit transcript identifier for the reference assembly that matched with the sequence comprising the GeneID given. Adjusted p-values were determined by a multiple test correction with the Benjamini and Hochberg method using p-values obtained from the Wald test. Only the transcripts with adjusted p-value < 0.01 were chosen as a conservative measure. At least half of upregulated DEGs were specific for each day, while others encoded proteins with similar descriptions between stages as in the case of actin, elongation factors, and cathepsin. Although annotated, some transcripts encode proteins not characterized in the databases as in the case of upregulated LOC592324 (accession: XM_030978239) and downregulated LOC575162 (accession: XM_030973217) at day 1. Downregulated DEGs shared the same characteristic with around half of their protein descriptions being specific for each stage and sharing proteins associated with transporters, aminopeptidases, lyases, homeobox proteins and aquaporin channels. Additional comparison between day 1 and 3 of regeneration using day 1 as the reference gave interesting results concerning downregulated nervous system genes (see Supplementary Table S8 online). Previous studies from our lab have shown that the nervous system within the mesentery is disrupted early in regeneration and slightly retracts or degenerates in the area adjacent to the evisceration rupture^30^. A decrease in the expression of nervous system-associated genes concurs with a negative effect on the mesentery nervous component.

**Table 1.** Top 10 upregulated DEGs at day 1 of regeneration.

| ID | E-value | GeneID | Description | Log2FC | Adjusted P-value |
| --- | --- | --- | --- | --- | --- |
| Transcript_7357 | 5.50E-15 | 580617 | cathepsin Z | 9.44↑ | 1.37e-10* |
| Transcript_331800 | 7.60E-51 | 592324 | uncharacterized protein LOC592324 | 8.03↑ | 1.33e-07* |
| Transcript_77032 | 8.00E-14 | 580195 | cubilin | 7.78↑ | 9.56e-05* |
| Transcript_370380 | 4.40E-19 | 574783 | serine/threonine-protein kinase NLK | 7.43↑ | 3.73e-05* |
| **Transcript_257047** | 2.00E-53 | 581500 | actin | 7.37↑ | 1.78e-04* |
| **Transcript_166464** | 3.60E-140 | 592801 | translation elongation factor 2 | 7.20↑ | 3.16e-04* |
| **Transcript_261599** | 9.00E-49 | 548620 | elongation factor 1 alpha | 7.15↑ | 4.40e-05* |
| **Transcript_121057** | 9.40E-62 | 593221 | V-type proton ATPase 16 kDa proteolipid subunit | 7.14↑ | 9.04e-04* |
| **Transcript_196171** | 8.30E-70 | 585216 | peptidyl-prolyl cis-trans isomerase | 6.88↑ | 5.02e-05* |
| **Transcript_276623** | 1.6E-44 | 752213 | Histone H4 | 6.71↑ | 1.97e-04* |

↑: Upregulated (Log2FC > 2); \*: Significant (Pvalue < 0.01); Log2FC: Log<sub>2</sub> Fold Change

**Table 2.** Top 10 downregulated DEGs at day 1 of regeneration.

| **ID** | **E-value** | **GeneID** | **Description** | **Log2FC** | **Adjusted P-value** |
| --- | --- | --- | --- | --- | --- |
| **Transcript_236563** | 3.10E-94 | 105439658 | protein FAM166B | -19.48↓ | 1.92e-05* |
| **Transcript_386663** | 8.80E-89 | 592851 | pancreatic lipase-related protein 2 | -11.15↓ | 4.67e-06* |
| **Transcript_182900** | 6.00E-233 | 580845 | sodium-coupled monocarboxylate transporter 1 | -10.09↓ | 2.33e-12* |
| **Transcript_383623** | 1.40E-245 | 583726 | histidine ammonia-lyase | -9.94↓ | 1.30e-04* |
| **Transcript_258721** | 3.80E-80 | 755759 | glutamyl aminopeptidase | -9.73↓ | 1.03e-11* |
| **Transcript_395355** | 4.00E-22 | 105438405 | echinoidin-like | -9.46↓ | 3.23e-02* |
| **Transcript_236625** | 1.90E-46 | 584191 | homeobox protein CDX-2 | -8.64↓ | 2.33e-03* |
| **Transcript_427783** | 2.00E-171 | 575162 | uncharacterized protein LOC575162 | -8.20↓ | 7.09e-10* |
| **Transcript_264429** | 0 | 583122 | sushi domain-containing protein 2 | -7.92↓ | 4.74e-21* |
| **Transcript_246361** | 7.3E-77 | 762564 | aquaporin-8 | -7.86↓ | 2.44e-07* |

↓: Downregulated (Log2FC < -2); \*: Significant (Pvalue < 0.01); Log2FC: Log<sub>2</sub> Fold Change

**Table 3.** Top 10 upregulated DEGs at day 3 of regeneration.

| ID | E-value | GeneID | Description | Log2FC | Adjusted P-value |
| --- | --- | --- | --- | --- | --- |
| Transcript_26623 | 8.00E-58 | 592324 | uncharacterized protein LOC592324 | 9.12↑ | 1.01e-09* |
| Transcript_368226 | 2.70E-79 | 584839 | ubiquitin | 7.82↑ | 7.54e-06* |
| Transcript_375637 | 1.20E-164 | 585147 | protein Wnt-6 | 7.68↑ | 3.64e-28* |
| Transcript_432544 | 1.50E-251 | 581500 | actin related protein 1 | 7.47↑ | 3.31e-05* |
| Transcript_166108 | 6.00E-98 | 575735 | major vault protein | 7.35↑ | 1.26e-04* |
| Transcript_187774 | 2.60E-142 | 581500 | actin CyI, cytoplasmic | 7.22↑ | 1.79e-05* |
| Transcript_270219 | 1.40E-56 | 579019 | ubiquitin-40S ribosomal protein S27a | 7.04↑ | 3.05e-05* |
| Transcript_290528 | 2.30E-57 | 580363 | transcription factor VBP | 6.95↑ | 9.12e-10* |
| Transcript_200945 | 4.20E-124 | 575203 | cathepsin L1 | 6.88↑ | 8.98e-04* |
| Transcript_206208 | 5E-148 | 592801 | translation elongation factor 2 | 6.85↑ | 5.92e-04* |

↑: Upregulated (Log2FC > 2); \*: Significant (Pvalue < 0.01); Log2FC: Log<sub>2</sub> Fold Change

**Table 4.** Top 10 downregulated DEGs at day 3 of regeneration.

| ID | E-value | GeneID | Description | Log2FC | Adjusted P-value |
| --- | --- | --- | --- | --- | --- |
| Transcript_200469 | 1.20E-102 | 589852 | aquaporin-8 | -11.08↓ | 2.52e-14* |
| Transcript_182900 | 6.00E-233 | 580845 | sodium-coupled monocarboxylate transporter 1 | -10.05↓ | 3.14e-12* |
| Transcript_236625 | 1.90E-46 | 584191 | homeobox protein CDX-2 | -9.57↓ | 6.50e-04* |
| Transcript_429455 | 2.50E-15 | 753235 | tenascin-N | -9.31↓ | 1.87e-03* |
| Transcript_43695 | 1.70E-204 | 579553 | b (0, +)-type amino acid transporter 1 | -8.17↓ | 2.53e-07* |
| Transcript_258721 | 3.80E-80 | 755759 | glutamyl aminopeptidase | -7.97↓ | 2.87e-22* |
| **Transcript_385039** | 6.40E-11 | 754702 | aminopeptidase N | -7.83↓ | 9.95e-26* |
| **Transcript_383623** | 1.40E-245 | 583726 | histidine ammonia-lyase | -7.82↓ | 1.19e-03* |
| **Transcript_235911** | 1.50E-113 | 100890912 | phospholipase B1, membrane-associated | -7.82↓ | 4.01e-06* |
| **Transcript_207127** | 1.1E-182 | 754102 | tubulin alpha-1A chain | -7.72↓ | 1.15e-06* |

↓: Downregulated (Log2FC < -2); \*: Significant (Pvalue < 0.01); Log2FC: Log<sub>2</sub> Fold Change

### Functional classification of DEGs

GO terms obtained from DEGs with a p-value less than 0.05 and considerable relation between the number of genes associated with the term (Gene Ratio) are shown in Figure 3. Most terms obtained from upregulated genes at both day 1 and 3, correlate with processes involved in transcription of DNA information into RNA, as well as the translation of mRNA into proteins. Enriched terms of biological process (BP) include various protein transport mechanisms; cellular component (CC) terms are characterized for processes involving translation activity; molecular function (MF) terms comprise protein transport, translation, and RNA binding. Downregulated terms showed a wider variety of associations. In general, terms associated with signaling systems, cell adhesion, metabolic processes and the immune system were found. Similar terms were shown for CC, being the membrane components an enriched factor for downregulated genes. MF terms associated with downregulated genes in both days represented mostly cell signaling processes (see supplementary Tables S10-S13 online).

**Figure 3.**
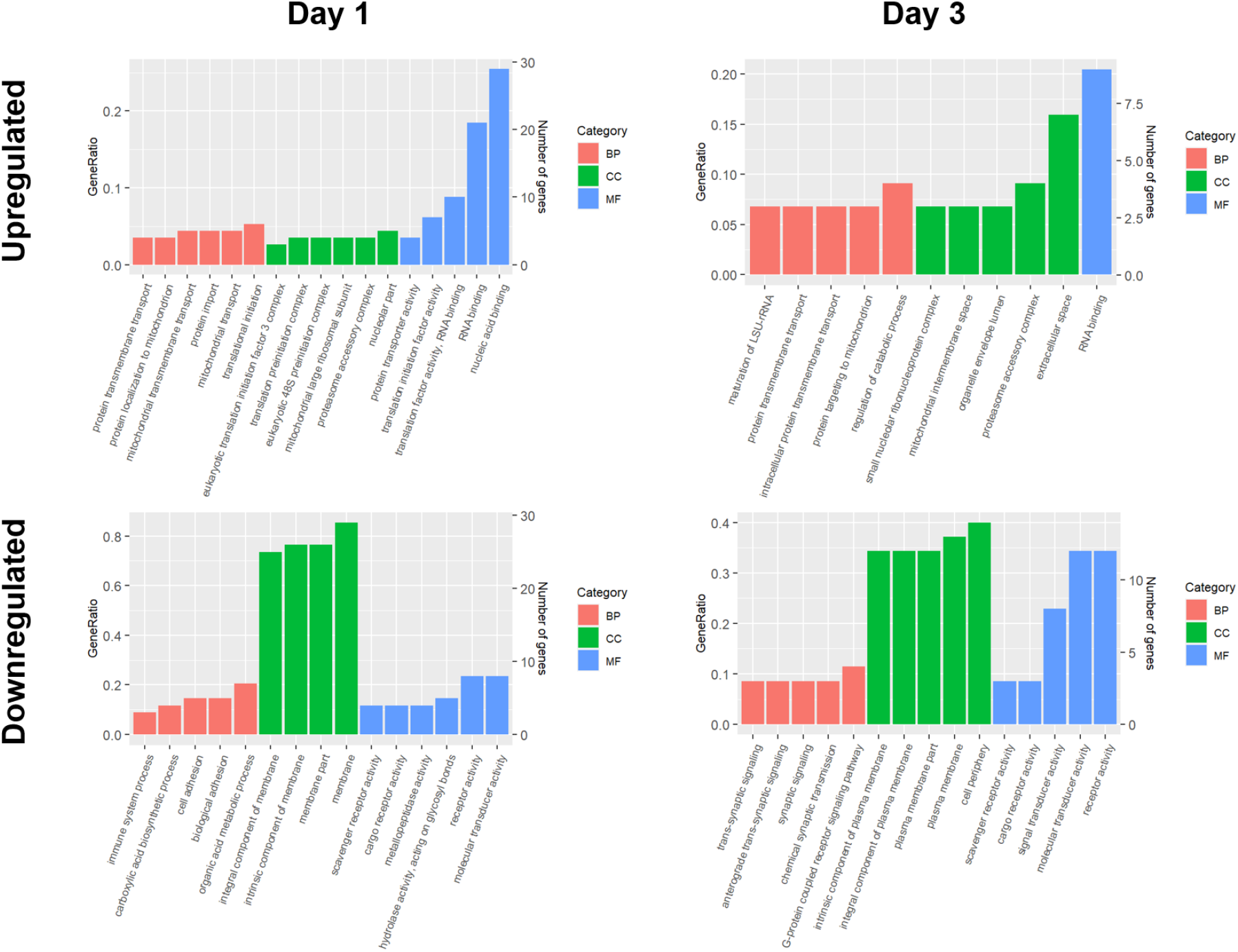
Gene ontology enrichment analysis of differentially expressed genes (DEGs) at day 1 and day 3 of regeneration using DAVID database. Ontology domains comprise biological process (BP), cellular component (CC) and molecular function (MF). The GeneRation is a measure of the gene count divided by the number of genes in the total group of genes assayed that belong to the specific gene category.

Pathways associated with intestinal regeneration were classified according to upregulated and downregulated genes for each day. Only resulting pathways with p-value < 0.05 were depicted in Figure 4. Enriched pathways for upregulated genes in day 1 and 3 were mostly associated with protein processing and transcriptional activity, being the pathway involved in proteasome activity the most enriched. Downregulated genes correlated mainly with various types of metabolic activity. Furthermore, enzymatic activity associated with peroxisomes were found to be enriched for both days. Intriguingly, phagosome activity was shown in pathways of upregulated genes for day 1, as well as the downregulated genes of the same day (see supplementary Tables S14-S17 online).

**Figure 4.**
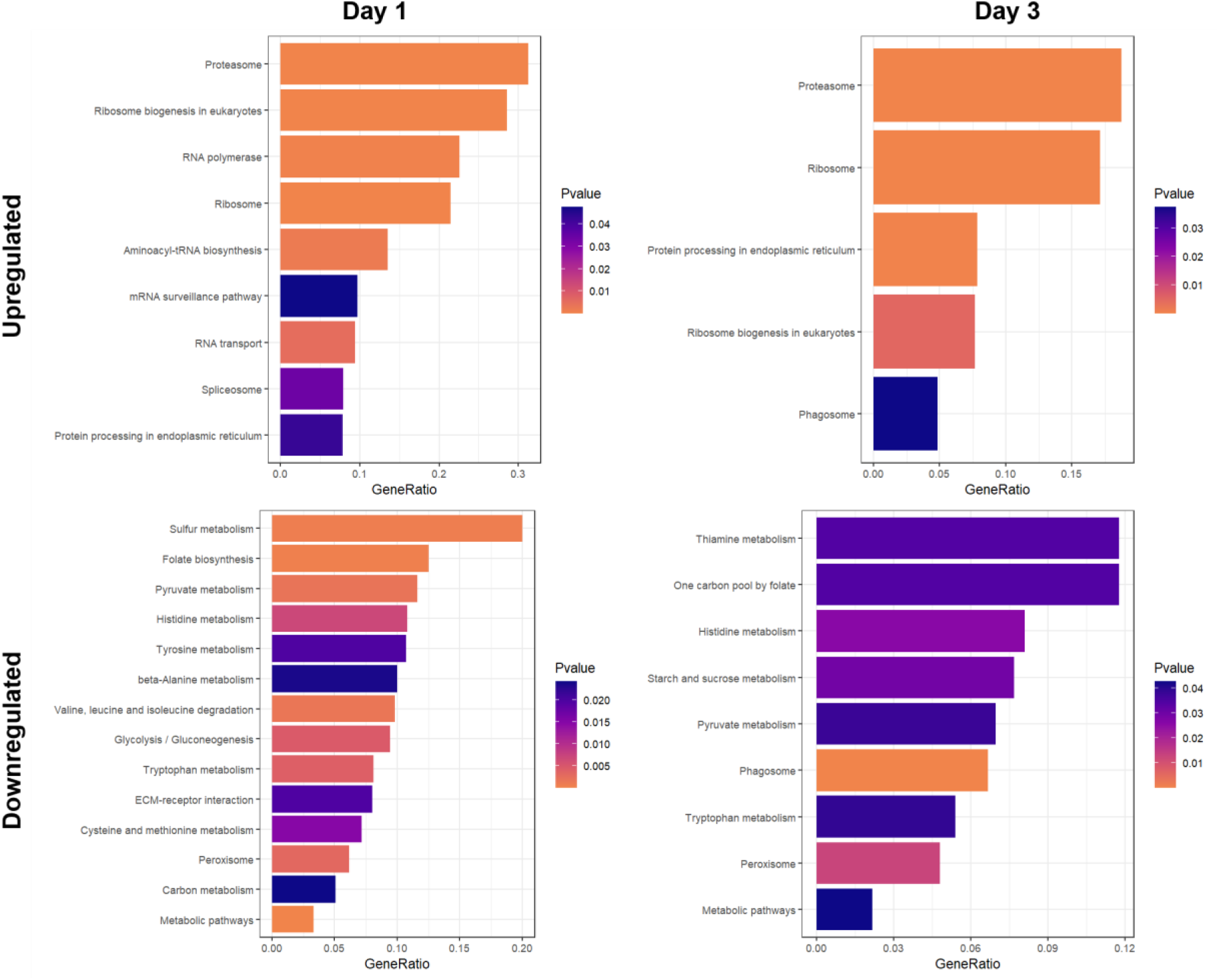
Pathway enrichment analysis of differentially expressed genes (DEGs) at day 1 and day 3 of regeneration using KEGG database. The GeneRation is a measure of the number of genes in the differentially expressed set divided by the total number of genes for that specific pathway found in KEGG.

### Differentially Expressed Transcription Factors (DETFs)

One of the first cellular events associated with regeneration is the dedifferentiation of the mesentery cells^31^. Such dedifferentiation is usually associated with cellular reprogramming and the activation or inactivation of specific transcription factors (TFs). Thus, we identified the annotated TFs with expression changes (see Supplementary Table S9 online). The heatmap in Figure 5 shows five main expression patterns. Group 1 (grey) corresponds to factors with moderate-to-low expression in normal tissues that increase during regeneration with a peak at day 1. Group 2 (light blue) corresponds to factors with high expression in normal tissues that decrease slightly in day 1 and 3. Group 3 (green) corresponds to factors with regular expression in normal mesentery that increase during the first three days of regeneration. Group 4 (cyan) corresponds to factors that are moderately expressed in normal tissue and decrease their expression during regeneration days 1 and 3. Finally, Group 5 (blue) includes factors that are weakly expressed in normal mesentery but increase their expression during days 1 and 3 of regeneration.

**Figure 5.**
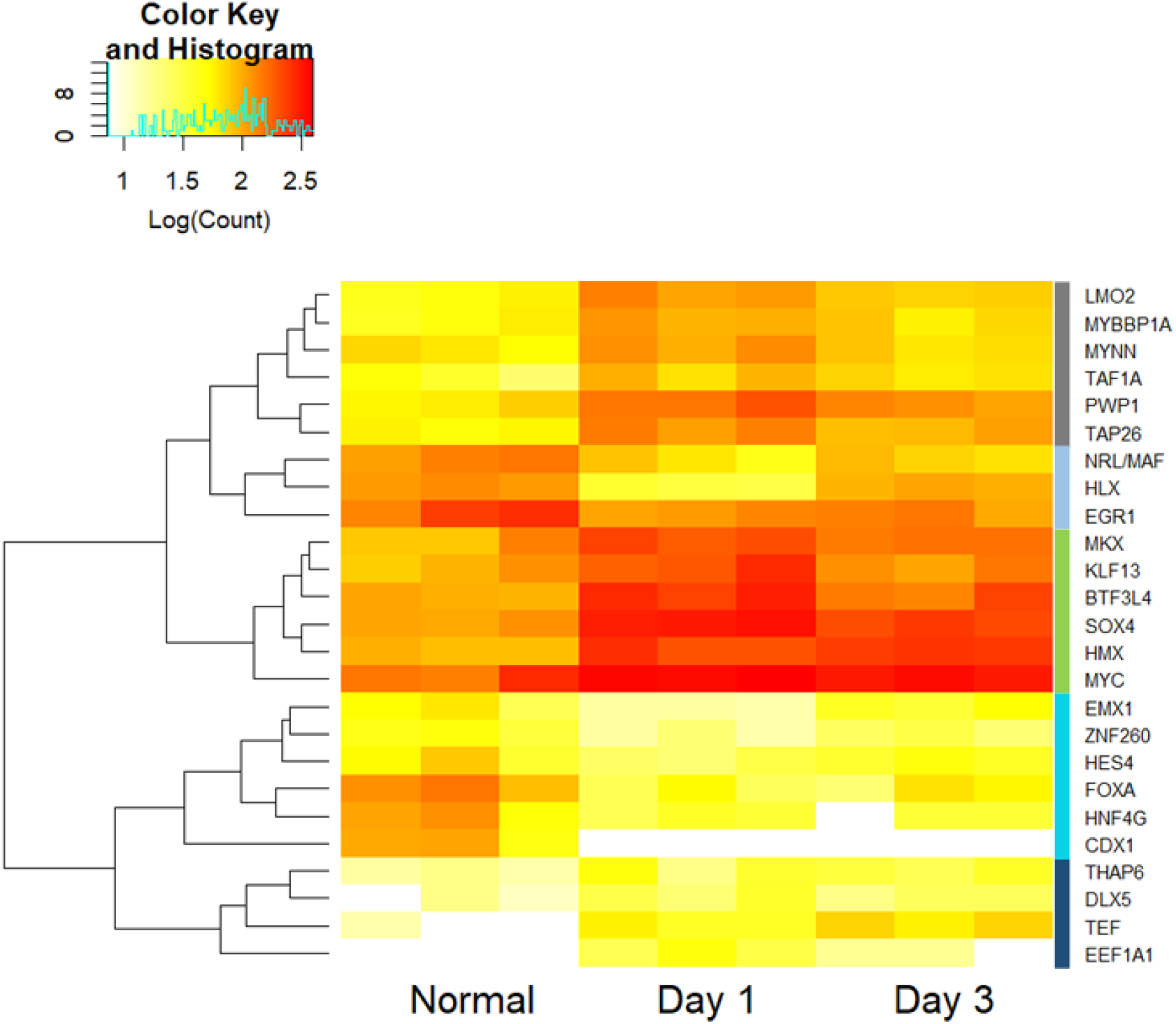
Heatmap of differentially expressed transcription factors (DETFs) in non-regenerating stage mesentery (Normal), and regenerating (days 1 3) intestines. DETFs showing a similar expression profiles are grouped in the dendrogram and labeled by colors in the right part of the heatmap. Logarithm count measures for expression values encompass non-negative integer values from the expression analysis.

To examine association with particular cellular processes or regenerative events, the TFs were clustered with differentially expressed transcripts and subjected to a functional analysis with GO. Groups 2 and 4 were mostly associated with cell adhesion and cell receptor activity in the membrane, while groups 1 and 3 were involved with biosynthesis and translation. Finally, TFs in group 5 were correlated with proteolytic and structural processes in the cell (see supplementary Table S19 online).

## Discussion

In the present study, using RNA-seq, we have constructed a large-scale differential gene expression profile of the regenerating intestine in *H. glaberrima*. We validated our results by comparing the expression of genes between our database and expression trends using different methods in previous studies. Results showed an identical expression pattern in the early, 1-& 3-days post evisceration (dpe) stages. Among the genes showing parallel upregulated expression between RNA-seq and previous methods, we found genes encoding proteins involved in cell differentiation and proliferation as in the case of Myc and Mtf^26,27^. Similarly, β-catenin showed no considerable change of expression in early stages from previous studies using RT-qPCR, and no changes were observed in our RNA-Seq results^26^. Other methods to detect gene expression were also found to give consistent results with our data. For instance, northern blot analysis of Serum amyloid A expression showed upregulation during day 3 of regeneration^20^. We found an increase in the expression of its mRNA at the same stage with RNA-seq. Furthermore, *in situ* hybridization studies, although not quantitative, do show differences in mRNA expression that coincide with our data. In this case, WNT9, BMP and mortalin showed considerable increased labeling in the early regenerate, while survivin appeared to have scarce and dispersed labeling in the regenerating mesenteric region^28,29^. Finally, we applied the RT-qPCR method to validate the expression of 7 genes with 12 expression values from day 1 and 3 of regeneration. The analysis showed a high correlation using both methods, thus reassuring the reliability of the analysis obtained in our experimental approach to study early stages of intestinal regeneration in *H. glaberrima*.

The high number of DETs for 1 & 3 dpe (>8000), contrasts with a previous study of intestinal regeneration in *A. japonicus* where less than 4000 DETs were found^23^. Notwithstanding, these differences can be attributed to the significance criteria established for each study. In *A. japonicus*, an adjusted p-value of < 0.001 was established as threshold for significance, while our study used a value of < 0.05. In fact, using the same significance criteria (see Supplementary Fig. S2 online), the number of DETs is similar (3587) to those reported in the *A. japonicus* study. Our results are also comparable with a study done in *E. fraudatrix* applying equivalent significance benchmarks (adjusted p-value < 0.05) documenting 17,227 DETs in regenerating intestine relative to intact gut^24^.

Our interest, however, lies beyond simply quantifying the number of DETs but targets the expression trends during early stages of intestinal regeneration to determine genes involved in cellular events such as dedifferentiation, proliferation and migration. It is in this analysis where the major discrepancies are observed between this study and those of others.

Comparison of top DEGs at day 3 of regeneration between *A. japonicus*^23^ and our data revealed no parallelism. All genes in up-regulated and down-regulated lists were different. Additional disparity was found in *E. fraudatrix* study. Although there are some differences in the evisceration and regeneration process of this species that might explain the differential gene expression, its regeneration process also involves dedifferentiation of cells within the mesenteric tissue, thus a certain degree of similarity would have been expected. Nonetheless, DETF identified in *E. fraudatrix*^24^ were not found in our study, or if present, their differential expression showed no correlation. Examples of this expression discrepancy: EGR1, TCF24 and SNAI2, upregulated in *E. fraudatrix*, downregulated in *H. glaberrima*.

There are even differences with previous studies from our own group using the same species^32^. Diverging expressions were found mostly with ECM-related genes such as collagen alpha-1, laminin alpha 1, stromelysin-3 (MMP II) and developmental related Hox 9 (see supplementary Table S7 online). In all cases, these genes were upregulated in our previous microarray study, whereas they were downregulated in our RNA-seq. As mentioned earlier we posit the differences between our results and previous studies stem from what each study used as a point of comparison to determine the gene expression profile. In all previous studies, the adult non-eviscerated intestine was used as the reference point to determine differential expression. However, as explained earlier, the ratio of the area of connective tissue to that of the mesothelium in the early intestinal rudiment is much smaller than in the adult intestine (see Fig. 1). Moreover, the adult intestine has a prominent mucosal layer that does not appear in the intestinal rudiment until the second week of regeneration. Therefore, the relative changes in cell/tissue distribution between the early regenerate and the fully formed organ are a clear cause for a skew in the resulting gene expression profile. Thus, the expression differences in previous studies might reflect differences in cell or tissue gene expression rather than differences in regeneration stages. On the other hand, our study takes these facts into account and focuses on a comparison with the normal mesentery which is a more trustworthy point of comparison because of similar distribution of cells and tissues.

Both functional and signal pathway analyses point out to certain activities that are central to the regeneration process. Overall upregulated transcripts were mostly involved with transcriptional processes, translation, and protein transport. This was expected as the regenerative process requires expression of abundant components for reconstitution of lost tissues. Post-translational protein processing and proteasome related pathways were also represented in the results. Both mechanisms appear to be evidently important for cell renewal especially the former which is needed for enzymatic modifications following synthesis. Proteasome-related transcripts have been previously observed to be upregulated during intestinal regeneration^33^ and activity of the proteasome complex has been classified as indispensable for stem cell maintenance^34^. Proteasome activity is also important in cell signaling as the degradation of particular proteins can cause large shifts in cell processes^35^. Downregulated GO terms include cell adhesion, which comprise proteins with diverse expression patterns such as the cadherins^36^. A possible reduction in cell adhesion is consistent with mesenterial cell migration toward the forming blastema and with EMT events. In this respect it is interesting to highlight the observed correlation of certain TFs expression with genes associated with cell adhesion. Immune system at day 1 appears surprising given the fact that this activity is believed to be activated during regeneration and previous studies have shown representation of inflammatory response terms at day 3^37^. However, there are no studies analyzing expression patterns as early as day 1, which could indicate a late immune response in the beginning of the process. There is also evidence of positive regeneration response by suppression of the immune response in tadpole tail regeneration^38^, which highlights the puzzling role of this complex network in regeneration. The term peroxisome pathway in this downregulated group, suggests a decreased activity towards removal of reactive oxygen species (ROS). The importance of ROS has been documented in various regeneration model organisms for its positive effect during regeneration^39,40,41^. Finally, and worth highlighting, the phagosome related pathway was represented in upregulated and downregulated transcripts for day 3. This tendency results from similar DEGs of actin and tubulin involved in phagosome motility and structure^42,43,44^. However, transcripts grouped in both expression trends were different, e.g. tubulin alpha-1A in downregulated and tubulin beta-4B in upregulated. Additionally, there were a greater number of transcripts promoting the phagocytic process in upregulated transcripts such as calnexin and calreticulin, two pro-phagocytic proteins with chaperone activity in endoplasmic reticulum^45^.

Among the DEGs between normal mesentery and the early stages of intestinal regeneration, we have focused on TFs for obvious reasons: These are the genes that play central roles in most cellular processes by activating or inhibiting the transcription of many downstream genes involved in the process.

The DETFs can be classified depending on the genes that are being regulated and the effect they produce when expressed. Thus, it is not surprising that many of the DETFs regulate the transcription process itself. For example, TAF1A, MYBBP1A and PWP1 are involved in the assembly of RNA polymerase I preinitiation complex, processing of ribosomal RNA and ribosome assembly, respectively^46,47,48^. Similarly, EEF1A1 has been recognized as a translation elongation factor^49^. Interestingly all these genes appear to increase their expression at day 1 and are still upregulated, but at a lower level at day 3. This suggests that the injury triggers an early response to establish the transcription machinery necessary for the subsequent regenerative processes.

Far more interesting are those TF associated with cellular identity. In this respect the differential expression observed with MYC calls for particular attention. MYC is one of the Yamanaka factors used to induce pluripotency^50^. Moreover, previous studies from our lab have shown that it is overexpressed during both intestinal and radial nerve regeneration, two processes where cells undergo dedifferentiation^51,52^. RNAi performed in the radial nerve shows that downregulation of MYC decreases the dedifferentiation of the radial glia^26^. Thus, our study fully supports the view that MYC is important in the early regenerative process suggesting that it is important for the dedifferentiation of the mesenterial cells.

One of the most interesting findings is that most DETFs share one common characteristic: they have been associated with developmental processes involving mesodermal derivatives. Many of them have been shown to activate or inhibit processes leading to muscle, cartilage, or bone formation^53,54,55^. In many cases they are involved in keeping cells in an immature state or inhibiting their differentiation^56,57^. Furthermore, some are directly associated with the formation of digestive system organs, such as the liver, pancreas, cloaca, stomach and even with enterocyte identity^54,58,59,60^. Among these factors we found various transcripts associated with developmental-related homeobox family of genes. Our results show activation of MKX, HMX, TAP26 and DLX5 at day 1, while inhibition of HLX, EMX1 and CDX1 is observed for the same day. This contrasting expression trend is particular of homeobox genes, which are characterized by their dynamic expression pattern as shown by members of the Hox family during sea urchin development^61^. Although homeobox TFs appear to be involved in countless mechanisms, functions associated with MKX, HLX and CDX1 are worth mentioning. MKX has been shown to have a role as a regulator of collagen production^53^, a structural protein previously shown to have important implications during intestine regeneration^54^. As for HLX and CDX1, their implication in enteric nervous system and intestine development^58,59^ makes them interesting study targets. This activity could also be associated with downregulation of neurogenic associated transcripts observed when comparing day 3 vs 1 (see Supplementary Table S8 online). Previous studies from our group have shown that an important plexus made up of nerves and fibers is present in the intestinal mesentery^30^. Following evisceration, the plexus connection to the intestinal tissues is lost, since the latter have been eliminated. Thus, many of the axons connecting the mesentery to the intestine would have been severed. Morphological changes induced by the evisceration have been documented including the disorganization of the fiber tracts and their retraction at the regenerating end^30^. Therefore, it can be hypothesized that significant changes in nervous system gene expression are taking place during early regeneration stages.

Other upregulated development-related factors include MYBBP1A, TEF, and MYNN. The first two have important roles in embryonic development^62,63^, whereas MYNN has a localized expression in muscle cells^55^. On the other hand, downregulated transcripts identified as NRL/MAF (neural retina leucine zipper, Maf type), EGR1 and FOXA, have all been associated with differentiation mechanisms. The former has been shown to have an essential role in rod photoreceptor development^64^; EGR1 has a vast range of activities including differentiation and proliferation, most importantly its association with other factors that regulate cell growth^65^; FOXA has a role in competence establishment of the foregut endoderm and induction of liver specific genes for hepatogenesis^60^. Finally, it is worth mentioning that two transcripts annotated as KLF13 and ZNF260 were found, respectively upregulated and downregulated at day 1. Both belong to the Krüppel family of transcriptional regulators involved in modulation of numerous developmental features^56,57^. Downregulation of most of these transcripts appears reasonable since their differentiation role is contrary to the dedifferentiation processes taking place at the onset of regeneration.

Finally, we identified a group of DETFs that have been associated with oncogenesis. SOX4 and LMO2 show upregulation after day 1 while HNF4G is downregulated in the same period. These cancer-related TFs were recognized among the top considerable transcripts with differential expression. Similarities between regeneration and cancer-related events have been studied for their involvement in dedifferentiating mechanisms during cellular reprogramming^66^. This compelling process of returning to an undifferentiated state during regeneration, confers high proliferative and migratory capacities to cells, without the loss of cell growth control distinctive of cancer^67^. In the case of SOX4 and LMO2, their role has been established as regulators with aberrant expression leading to an oncogenic state^68,69^. As for HNF4G, its upregulation has been correlated as a prognostic marker for cancer development by promoting cell proliferation^70^, which is interesting considering the opposite expression trend in our results. Other transcripts related to THAP6 and BTF3L4 that show upregulation at day 1, have been associated with an apoptotic regulatory role^71,72^. This contrasting expression portrays regulatory differences of cancer related mechanisms being activated for dedifferentiation in early stages.

In summary, diverse TF activity indicates the abundant network of mechanisms taking place during intestinal regeneration. Various TF were found to be directly or indirectly involved with regenerative processes. Furthermore, functional analysis regarding transcripts coexpressed with TFs showed processes associated with a structuration of the cell, as well as cell communication mechanisms during the early regenerative stage. Although their primary role is rather aimed at developmental purposes in most cases, there are previously studied TF such as MYC whose DE can be proposed as landmarks of regeneration.

## Conclusion

This is the first study with a mesentery directed approach for improving specificity in the characterization of differential gene expression in regenerating tissues. Furthermore, it is also the first one in sea cucumbers for analysis of differential expression during intestinal regeneration at a stage as early as 24 hours after the organism has expelled its internal organs. An expected high number of DETs were found using high-throughput sequencing methods and numerous functional processes associated with regeneration. What is more appealing, TF activity correlated with regenerative machinery was diverse in expression trends that did not necessarily belong to the same group of processes. Further studies can be considered to unravel the regulatory networks associated with TF activity and connect them with distinctive cellular processes.

## Materials and Methods

### Animals and treatment

Sea cucumbers were collected from the northeastern rocky coast of Puerto Rico^73^. Evisceration was carried out by an intracoelomic injection of 0.35M KCl. Subsequently, the organisms were left in salt-water aquaria for 1 and 3 dpe to undergo regeneration. Uneviscerated animals kept under the same conditions and for the same lengths of time as the eviscerated animals were used as controls. Prior to dissection, animals were anesthetized by immersion in ice-cold water for 45 minutes. A dorsal incision was made to expose the internal cavity of the organism and allow dissection of the mesentery separated from the body wall and intestine. In the case of non-eviscerated organisms, the intestine was separated from the mesentery that remained attached to the body wall and removed using small surgical scissors. The mesentery was obtained throughout its length from the anterior end of the animal (next to the esophagus) to the posterior end (next to the cloaca). Dissected tissues were placed in RNAlater (Sigma, USA) solution and stored at 4ºC until the RNA extraction was performed.

### RNA extraction

RNA extraction was done with a combination of the method established by Chomczynski (1993) using Tri-reagent® (N.93289, Sigma, USA) and the RNAeasy mini kit (Qiagen, Germany). Three extractions were made for each stage (normal, day 1 and day 3) and each extraction consisted of pooled tissue samples from three organisms at the same stage. The concentration and quality of the RNA was determined by using the 2100 Bioanalyzer (Agilent Technologies, USA). Samples with a concentration greater than 200 ng/μL and an RNA Integrity Number (RIN) value of ≥ 8 were used for sequencing.

### Sequencing of extracted RNA

RNA isolated from the mesentery of regenerating and non-regenerating sea cucumbers was sequenced at the Sequencing and Genotyping Facility of the University of Puerto Rico. Sequencing libraries were prepared using the TruSeq Stranded mRNA Library Prep Kit (Illumina, USA). Paired-end sequencing (2 × 150) was performed with the Illumina NextSeq® 500 system.

### Transcriptome assembly

Raw reads were uploaded to the High-Performance Computing Facility of the University of Puerto Rico. Samples for the assembly comprised 3 from normal mesentery, 3 from day 1 regenerating intestine, 5 from day 3 regenerating intestine, and 6 samples from the radial organ complex from a previous study by Mashanov et al^25^. Preprocessing steps of quality, trimming and filtering were done using FastQCv0.11.5^74^ and Trimmomaticv0.36^75^ applying the following parameters ILLUMINACLIP:2:30:10 LEADING:3 TRAILING:3 SLIDING-WINDOW:4:15 MINLEN:36. A kmer trimming and digital normalization was applied previously to the assembly process using the khmer package^76^ and following the khmer protocol documentation ^77^. A de novo transcriptome assembly was carried out using Trinityv2.9.0^78^ with the following parameters --no_bowtie --seqType fq --max_memory 105G --left left.fq --right right.fq --CPU 14.

### Differential gene expression

The differential expression analysis was carried out with Salmon^79^ applying the default parameters and using the quality filtered RNA-seq data and the assembled transcriptome generated with Trinity as reference. Three samples of normal mesentery, 3 of day 1 regenerating intestine and 3 of day 3 regenerating intestine were used for the differential expression analysis. Equivalence classes generated by Salmon were used to hierarchically cluster contigs by sequence similarity and expression using Corsetv1.06^80^ with default settings and the parameter -D 9 for all samples (9). Lastly, DESeq2 tool package^81^ was used to extract differentially expressed clusters of contigs from the read count files generated with Salmon. Files with less than 30 read counts were filtered from every biological replicate of each sample. Workflow used in the differential expression analysis can be found at https://github.com/davidjquispe/DGE_Analysis.

### Functional annotation

The annotation pipeline Dammit^82^ was used for annotating the transcriptome using the default parameters. Protein databases Pfam-A^83^, Rfam^84^, OrthoDB^85^ and uniref90^86^ were used for the annotation process with a cutoff value of 1e-05. The protein sequences from the genome of the sea urchin *Strongylocentrotus purpuratus* (Accession: GCA_000002235.4) were used as reference for the annotation process. Annotation completeness was evaluated with BUSCO^87^.

### Gene ontology and enrichment analysis

In order to have a broader understanding of the molecular processes that take place at different stages of gut regeneration we mapped Entrez Gene IDs of our transcripts using DAVID gene functional classification^88^ tool to obtain a gene ontology (GO) analysis. Only DEGs with significant p-value (<0.05) and absolute expression level higher than 2 (Log2FC) were associated with GO terms. Results of ontology terms were summarized with Revigo^89^. Pathway information was obtained by mapping KEGG ids from *S. purpuratus* datasets to *H. glaberrima* Entrez Gene IDs using kegga function from the Bioconductor package limmav3.11^90^. Gene ratio was manually calculated for each individual pathway using the number of DEGs for a term divided by the total number for that term. All transcripts used in the GO and pathway enrichment analysis were filtered based on their p value (<0.05).

### Transcription factor analysis

Sequences annotated as transcription factors (TFs) were selected among the 150 most and least expressed genes from the expression analysis by individual extraction based on their annotation name. A heatmap was generated with the gplots^91^ package using the TF that had an absolute Log2FC value higher than 1. Transcription factors were clustered along with annotated genes from the differential expression analysis using dist function to calculate the distance from each gene and hclust function for clustering from statsv3.6.0^92^ (see supplementary Table S18 online). Clusters of genes were subjected to a functional analysis using the DAVID database^88^.

### Validation of Gene Expression

Gene expression values (Log2FC) obtained from RNA-seq data were verified using the QuantStudio 3 Real-Time PCR system (Applied Biosystems, USA) and SYBR Green I as the DNA intercalating agent. A total of 7 genes were used for the analysis (see Supplementary Table S4 online) among which there was the mortalin gene analyzed in a previous study using the same RT-qPCR methodology^28^. Reaction volumes comprised 12 μL SYBR Green I, 0.6 μL of each primer and 1 μL of DNA, the rest of the volume was adjusted to 20 μL with nuclease-free water. The Ct values (see Supplementary Table S5 online) were analyzed with the 2-ΔΔCt (Livak)^93^ method using NADH as the housekeeping gene and the normal stage as the calibrator. A total of 12 expression values from day 1 and 3 regenerating intestine were compared between RNA-seq and RT-qPCR using equivalent measuring units of Log2FC.

## Data availability

The raw RNA-seq data can be found at the SRA NCBI Database under the BioProject accession PRJNA660762. Assembly has been deposited at DDBJ/EMBL/GenBank under the accession GIVL00000000. The version described in this paper is the first version, GIVL01000000. The files showing read counts and differential expression values were deposited in the GEO database under series record GSE160340. Custom scripts and data for differential gene expression are available at https://github.com/davidjquispe/DGE_Analysis.

## Acknowledgments

This project was funded by NIH (Grant R15NS01686 and R21AG057974) and the University of Puerto Rico. The authors acknowledge the help from the Sequencing and Genomics Facility of the UPR Río Piedras & MSRC/UPR (NIH/NIGMS-Award Number P20GM103475) that are supported by an Institutional Development Award (IDeA) from the National Institute of General Medical Sciences of the NIH under grant number P20GM103475.

## Author contributions statement

D.Q.P. performed preliminary work for data generation. J.M.F. built reference transcriptome and annotation with contribution from S.C.G. and input from H.O.Z. in computational proceedings. D.Q.P. and J.G.A carried out the analysis of results and drafting of the paper. All authors reviewed the manuscript.

