## Supplemental Figures for "Transcriptomic Analysis of Early Stages of Intestinal Regeneration in *Holothuria glaberrima*"

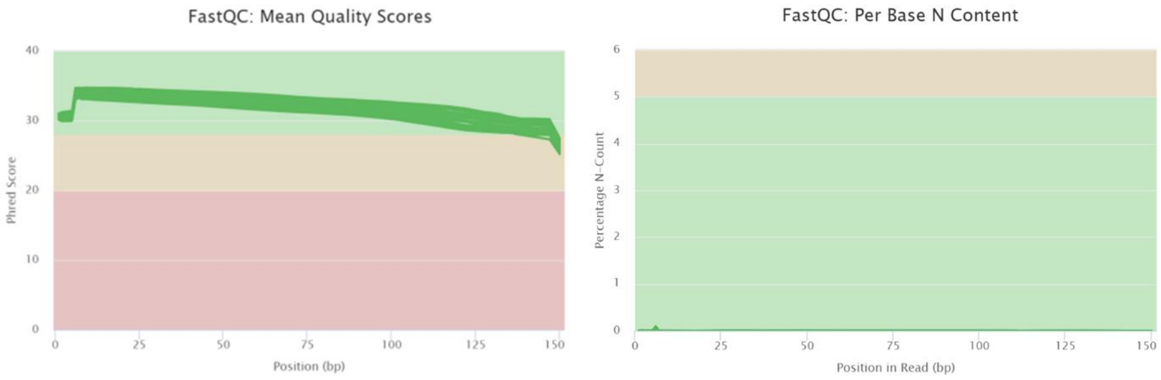


**Figure S1.** Quality assessment of reads after processing with Trimmomatic. All reads obtained a Phred score of 30 (left) meaning a 99.9% of accuracy and a null per base N content (right).

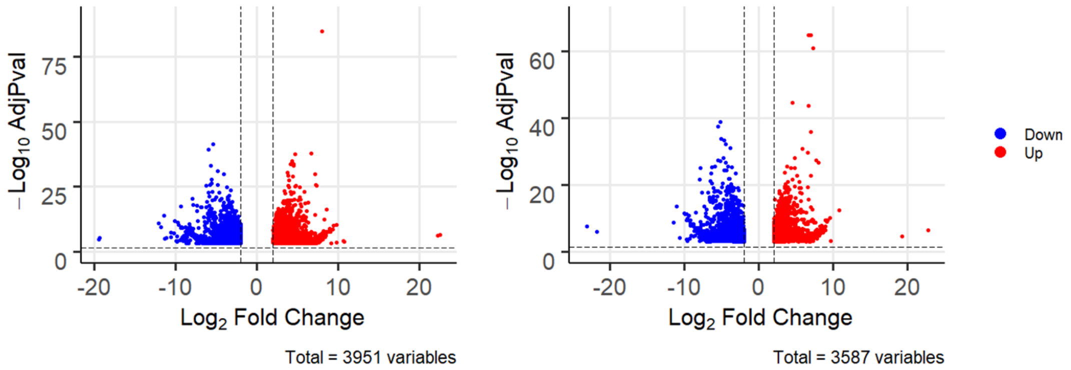


**Figure S2.** Differential gene expression analysis with adjusted p-value < 0.001.
