## Supplemental Tables for "Transcriptomic Analysis of Early Stages of Intestinal Regeneration in *Holothuria glaberrima*"

Table S1. Results of transcriptome assessment with BUSCO

| Parameter | BUSCO result |
|---|---|
| **Core genes queried** | 978 |
| **Complete core genes detected** | 99.1% |
| **Complete single copy core genes** | 27.2% |
| **Complete duplicated core genes** | 71.9% |
| **Fragmented core genes detected** | 0.4% |
| **Missing core genes** | 0.5% |

Table S2. Transcriptome length statistics and composition assessments with gVolante

| Parameter | Result |
|---|---|
| **Number of sequences** | 491 436 |
| **Total length (nt)** | 408 930 895 |
| **Longest sequence (nt)** | 34 610 |
| **Shortest sequence (nt)** | 200 |
| **Mean sequence length (nt)** | 832 |
| **N50 sequence length (nt)** | 1 691 |

Table S3. RNA-seq data read statistic values

| Accession (SRA) | Sample | Quantity of reads | | Mapped reads |
|---|---|---|---|---|
| | | Before Filtering | After Filtering | |
| SRR12564573 | NormalA | 89 424 588.00 | 88 112 720.00 | 94.18% |
| SRR12564572 | NormalB | 74 105 862.00 | 72 983 434.00 | 93.79% |
| SRR12564570 | NormalC | 60 626 094.00 | 59 767 896.00 | 91.55% |

| | | | | |
|---|---|---|---|---|
| SRR12564564 | Day1A | 32 499 910.00 | 31 721 976.00 | 90.67% |
| SRR12564563 | Day1B | 35 164 514.00 | 34 237 960.00 | 91.37% |
| SRR12564571 | Day1C | 43 519 280.00 | 42 179 220.00 | 91.53% |
| SRR12564567 | Day3A | 64 094 536.00 | 62 995 792.00 | 90.26% |
| SRR12564566 | Day3B | 71 654 462.00 | 70 158 078.00 | 91.07% |
| SRR12564565 | Day3C | 71 259 560.00 | 70 418 332.00 | 90.75% |
| SRR12564569 | Day3D* | 107 688 184.00 | 106 083 898.00 | - |
| SRR12564568 | Day3E* | 108 372 334.00 | 106 767 522.00 | - |

*Samples used for the assembly but not for differential expression analysis

Table S4. Genes and primers used for RT-qPCR

| Gene | Primers |
|---|---|
| **VBP** | F: AGGTCAAAATGTCCGGCCTCTGT<br>R: TCACGCTGGCGGTGTCAAAACT |
| **FoxA** | F: ACGGCGAAGGTAGCATCCGTTT<br>R: ATGGCAATCCAGCAAGCGCCAA |
| **Wnt6** | F: CCTGCAATAATCCGGTGAGT<br>R: CGCGAATGTAAATGTCATGG |
| **Sox4** | F: CGCTCTCCATTTCAGAGGAC<br>R: GTCTTTCTGCCTCCTCAACG |
| **Tap26** | F: TGCCTGAGCTTTCTGTTCCT<br>R: CCCTCAAAGAATGGTGGAAA |
| **NADH** | F: CAATGGTTGTTGCTGGAGTCTTT R:<br>CGCAGAAGTAGCCGCGAATAT |
| **Tenascin-N** | F: CGTTGAAATACCGTCCATCC<br>R: AACGAAGGATACGCTGGAGA |

Table S5. Mean Ct values obtained from RT-qPCR

| Gene | Stage | Ct Mean | Ct SD |
|---|---|---|---|
| **VBP** | Normal | 28.636 | 1.268 |
| | Day 1 | 28.263 | 0.640 |
| | Day 3 | 27.958 | 0.738 |
| **FoxA** | Normal | 23.611 | 0.371 |
| | Day 1 | 27.643 | 3.124 |
| | Day 3 | 27.824 | 3.201 |
| **Wnt6** | Normal | 28.607 | 1.399 |

| | | | |
|---|---|---|---|
| | Day 1 | 25.253 | 1.965 |
| | Day 3 | 27.842 | 3.703 |
| **Sox4** | Normal | 26.038 | 0.784 |
| | Day 1 | 25.424 | 1.143 |
| | Day 3 | 29.242 | 1.014 |
| **Tap26** | Normal | 27.634 | 1.319 |
| | Day 1 | 26.727 | 0.483 |
| | Day 3 | 29.512 | 1.311 |
| **NADH** | Normal | 18.843 | 0.271 |
| | Day 1 | 20.829 | 0.355 |
| | Day 3 | 22.583 | 0.341 |
| **Tenascin** | Normal | 21.025 | 3.217 |
| | Day 3 | 30.992 | 1.519 |

Table S6. Comparison of expressed transcripts in RNA-seq with previous studies

| **Gene** | **RNA-seq** | | **RT-PCR/qPCR** | **Northern Blot/In situ** |
|---|---|---|---|---|
| | Day 1 (Log2FC) | Day 3 (Log2FC) | Day 3 | Days 2-3 |
| **Myc** | 2.72 ↑ | 2.05 ↑ | ↑ | NA |
| **β-catenin** | 0.88 | $0.46^{NS}$ | $-^{NS}$ | NA |
| **Melanotransferrin** | 3.55 ↑ | 3.7 ↑ | ↑ | NA |
| **Serum amyloid A protein** | 2.39 ↑ | 3.2 ↑ | NA | ↑ |
| **WNT9** | 2.69 ↑ | 2.88 ↑ | NA | ↑ |
| **BMP** | 3.53 ↑ | 3.31 ↑ | NA | ↑ |
| **survivin** | $-0.90^{NS}$ | $-0.31^{NS}$ | $-^{NS}$ | - |

↑: Upregulated; ↓: Downregulated; -: No significant change; NS: Not significant (Pvalue > 0.05)

Table S7. Comparison of expressed transcripts in RNA-seq with microarray study

| **Gene** | RNA-seq | Microarray |
|---|---|---|

| | Day 1 (Log2FC) | Day 3 (Log2FC) | Day 3 |
|---|---|---|---|
| **Tensc-R** | 3.26↑ | 2.03 [NS] | ↑ |
| **Actin-1** | 6.12↑ | 7.47↑ | ↑ |
| **Actin-2** | 1.71↑ | 1.56↑ | ↑ |
| **Collagen alfa-1** | -1.54↓ | -0.67 [NS] | ↑ |
| **Laminin alpha1** | -3.67↓ | -2.79↓ | ↑ |
| **Stromelysin-3 (MMP-11)** | -4.16↓ | -2.65↓ | ↑ |
| **Hox9** | 1.03↑ | -0.83 [NS] | ↑ |
| **Krueppel like** | -1.09↓ | 0.12 [NS] | ↓ |
| **Gelsolin** | -0.79 [NS] | -2.69↓ | ↓ |

↑: Upregulated; ↓: Downregulated; -: No significant change; NS: Not significant (Pvalue > 0.05)

Table S8. Differentially expressed transcripts between day 3 vs 1 comparison

| ID | Description | Log2FC | Adjusted p-value |
|---|---|---|---|
| **Transcript_223195** | 15-hydroxyprostaglandin dehydrogenase [NAD(+)] | 2.13 | 3.58E-02 |
| **Transcript_298217** | 4-aminobutyrate aminotransferase, mitochondrial | 2.98 | 2.81E-05 |
| **Transcript_334800** | 40S ribosomal protein S15 | 6.73 | 1.89E-04 |
| **Transcript_254071** | actin, cytoskeletal 3B\|actin, cytoskeletal 3 | 3.00 | 1.19E-02 |
| **Transcript_432551** | actin, muscle | 3.65 | 5.45E-03 |
| **Transcript_383802** | adenylyltransferase and sulfurtransferase MOCS3-like | 3.21 | 3.38E-02 |
| **Transcript_308851** | alcohol dehydrogenase class-3 | 2.33 | 1.59E-02 |
| **Transcript_425811** | calreticulin | 2.45 | 9.78E-08 |
| **Transcript_298293** | collagen alpha-1(XII) chain\|tenascin-X | 2.08 | 7.99E-05 |
| **Transcript_196327** | creatine kinase, flagellar | 5.16 | 4.69E-02 |
| **Transcript_380459** | cryptochrome-2 | 2.03 | 3.75E-03 |
| **Transcript_226806** | cytochrome P450 3A24 | 2.25 | 1.83E-02 |
| **Transcript_262552** | cytochrome P450 3A9 | 4.03 | 3.31E-02 |
| **Transcript_175314** | cytochrome P450 4V2 | 2.00 | 4.85E-02 |
| **Transcript_172290** | cytosol aminopeptidase | 2.21 | 1.97E-05 |

| | | | |
|---|---|---|---|
| **Transcript_184414** | dehydrogenase/reductase SDR family member 7 | 2.25 | 1.39E-03 |
| **Transcript_175770** | deleted in malignant brain tumors 1 protein | 2.65 | 2.63E-02 |
| **Transcript_461577** | deoxyribodipyrimidine photo-lyase | 2.68 | 2.66E-03 |
| **Transcript_432963** | DNA replication licensing factor mcm5 | 2.57 | 4.81E-07 |
| **Transcript_359454** | dynein light chain LC6, flagellar outer arm | 2.35 | 6.25E-04 |
| **Transcript_347359** | ectonucleotide pyrophosphatase/phosphodiesterase family member 7-like | 3.16 | 4.27E-03 |
| **Transcript_273238** | epidermal growth factor-like protein 7 | 2.49 | 1.23E-05 |
| **Transcript_276903** | ERI1 exoribonuclease 2 | 2.86 | 3.80E-03 |
| **Transcript_365063** | extracellular transglutaminase | 2.62 | 5.99E-09 |
| **Transcript_350713** | fibrillin-1 | 2.08 | 9.82E-03 |
| **Transcript_356914** | ficolin-2-like | 3.74 | 1.04E-02 |
| **Transcript_179574** | ficolin-2-like | 3.05 | 3.76E-02 |
| **Transcript_221519** | formin-J | 2.02 | 1.31E-04 |
| **Transcript_358088** | fucose mutarotase | 2.46 | 1.86E-08 |
| **Transcript_351559** | G patch domain-containing protein 3 | 4.02 | 5.91E-03 |
| **Transcript_185057** | GDH/6PGL endoplasmic bifunctional protein\|hexose-6-phosphate dehydrogenase (glucose 1-dehydrogenase) | 2.07 | 2.55E-06 |
| **Transcript_402303** | GTP 3',8-cyclase, mitochondrial\|cyclic pyranopterin monophosphate synthase, mitochondrial\|molybdenum cofactor biosynthesis protein 1 | 2.32 | 8.19E-03 |
| **Transcript_438891** | H2.0-like homeobox protein\|Homeobox domain-containing protein | 3.45 | 1.75E-11 |
| **Transcript_251717** | hairy/enhancer-of-split related with YRPW motif protein 1 | 3.17 | 2.39E-02 |
| **Transcript_456276** | heparan sulfate glucosamine 3-O-sulfotransferase 1 | 3.31 | 1.15E-02 |
| **Transcript_438353** | homeobox protein EMX1 | 3.86 | 8.19E-03 |
| **Transcript_290553** | isoamyl acetate-hydrolyzing esterase 1 homolog | 2.80 | 5.95E-05 |
| **Transcript_409717** | isochorismatase domain-containing protein 2\|isochorismatase domain-containing protein 2, mitochondrial | 2.95 | 4.66E-03 |
| **Transcript_258900** | KRP170 | 2.69 | 1.74E-02 |
| **Transcript_189831** | L-gulonolactone oxidase | 2.45 | 6.22E-03 |
| **Transcript_297492** | laminin subunit alpha-2\|laminin subunit alpha-1 | 2.21 | 1.74E-05 |
| **Transcript_305145** | LOW QUALITY PROTEIN: DNA replication licensing factor mcm2 | 2.94 | 6.14E-07 |
| **Transcript_381204** | LOW QUALITY PROTEIN: transcription factor Sox-10 | 3.54 | 6.83E-09 |
| **Transcript_285150** | LOW QUALITY PROTEIN: zygotic DNA replication licensing factor mcm6-B | 2.29 | 2.08E-07 |
| **Transcript_294196** | low-density lipoprotein receptor-related protein 1 | 2.14 | 2.64E-04 |
| **Transcript_328691** | MAM and LDL-receptor class A domain-containing protein 1 | 2.14 | 7.82E-04 |
| **Transcript_257273** | methylsterol monooxygenase 1 | 3.02 | 3.79E-03 |

| | | | |
|---|---|---|---|
| **Transcript_324570** | microfibril-associated glycoprotein 4\|tenascin-N | 2.48 | 3.32E-02 |
| **Transcript_263515** | polycystic kidney disease protein 1-like 2 | 2.32 | 1.61E-02 |
| **Transcript_269460** | probable D-lactate dehydrogenase, mitochondrial\|lactate dehydrogenase D | 3.55 | 2.01E-02 |
| **Transcript_236563** | protein FAM166B-like | 17.42 | 4.96E-04 |
| **Transcript_437843** | protein lin-52 homolog | 2.10 | 2.16E-03 |
| **Transcript_375637** | protein Wnt-6 | 2.25 | 2.75E-03 |
| **Transcript_197666** | putative aminopeptidase W07G4.4 | 2.06 | 2.59E-04 |
| **Transcript_311247** | putative hydroxypyruvate isomerase | 2.52 | 5.99E-04 |
| **Transcript_443748** | pyridine nucleotide-disulfide oxidoreductase domain-containing protein 2\|pyridine nucleotide-disulphide oxidoreductase domain 2 | 2.41 | 1.73E-02 |
| **Transcript_297199** | ryncolin-1-like | 5.18 | 3.37E-02 |
| **Transcript_406209** | sulfite oxidase | 3.85 | 2.99E-07 |
| **Transcript_54760** | testis-specific serine/threonine-protein kinase 4-like | 5.30 | 3.42E-02 |
| **Transcript_251414** | transcription factor Sp5 | 2.40 | 4.19E-03 |
| **Transcript_384474** | transmembrane protein KIAA1109 | 2.08 | 1.48E-05 |
| **Transcript_266841** | tuftelin | 2.47 | 1.33E-02 |
| **Transcript_368226** | ubiquitin | 6.29 | 1.18E-03 |
| **Transcript_270219** | ubiquitin-40S ribosomal protein S27a | 6.47 | 4.00E-04 |
| **Transcript_437026** | uncharacterized protein K02A2.6-like | 3.43 | 2.72E-05 |
| **Transcript_190019** | uncharacterized protein K02A2.6-like\|Reverse transcriptase domain-containing protein | 2.32 | 1.79E-02 |
| **Transcript_291904** | uncharacterized protein LOC105438010\|Glycoside hydrolase family 31 domain containing protein | 2.14 | 3.79E-04 |
| **Transcript_283678** | uncharacterized protein LOC583353\|neurotrypsin | 4.75 | 9.63E-03 |
| **Transcript_26623** | uncharacterized protein LOC592324 | 7.59 | 2.53E-06 |
| **Transcript_442525** | uncharacterized protein LOC753842\|Protease inhibitor I35 (TIMP) domain containing protein | 2.01 | 2.72E-04 |
| **Transcript_270528** | uncharacterized protein LOC757055\|glyoxylate/hydroxypyruvate reductase A HPR2 | 2.01 | 7.71E-05 |
| **Transcript_263687** | valacyclovir hydrolase-like | 2.34 | 1.86E-03 |
| **Transcript_428635** | zygotic DNA replication licensing factor mcm3 | 2.64 | 3.93E-04 |
| **Transcript_167684** | | | |
| **Transcript_294311** | - | -3.44 | 2.27E-02 |
| **Transcript_251847** | - | -2.32 | 7.18E-05 |
| **Transcript_254514** | - | -2.01 | 2.89E-02 |
| **Transcript_395263** | 14-3-3 family protein artA\|14-3-3 protein 3 | -5.90 | 2.83E-03 |
| **Transcript_285870** | 26S proteasome non-ATPase regulatory subunit 5 | -3.03 | 1.24E-09 |

| | | | |
|---|---|---|---|
| **Transcript_42693** | 26S proteasome regulatory subunit 6A-B\|26S protease regulatory subunit 6A-B\|26S protease regulatory subunit 6A-like | -2.16 | 6.22E-06 |
| **Transcript_48454** | 40S ribosomal protein S13 | -3.47 | 4.95E-02 |
| **Transcript_345624** | 40S ribosomal protein S16 | -3.43 | 4.17E-02 |
| **Transcript_399286** | 40S ribosomal protein S9 | -4.03 | 1.07E-03 |
| **Transcript_382713** | 60S ribosomal protein L10a | -2.42 | 9.80E-03 |
| **Transcript_6160** | 60S ribosomal protein L19 | -3.63 | 1.40E-03 |
| **Transcript_1415** | 60S ribosomal protein L27a | -3.01 | 1.48E-02 |
| **Transcript_110850** | 60S ribosomal protein L3 | -4.01 | 1.67E-05 |
| **Transcript_274172** | 60S ribosomal protein L3 | -3.54 | 3.44E-03 |
| **Transcript_242646** | 60S ribosomal protein L31 | -3.19 | 2.25E-02 |
| **Transcript_258108** | 60S ribosomal protein L8 | -4.33 | 5.37E-05 |
| **Transcript_438569** | acetylcholine receptor subunit beta\|gamma-aminobutyric acid receptor subunit gamma-2 | -2.21 | 8.54E-03 |
| **Transcript_310621** | acid-sensing ion channel 1A | -4.52 | 9.37E-03 |
| **Transcript_388716** | actin CyI, cytoplasmic | -4.82 | 1.87E-02 |
| **Transcript_305867** | actin CyI, cytoplasmic | -2.41 | 1.99E-02 |
| **Transcript_198859** | actin-5C | -2.66 | 1.48E-02 |
| **Transcript_407345** | actin, muscle | -2.21 | 2.99E-07 |
| **Transcript_225582** | adenylate cyclase type 3 | -2.06 | 2.98E-03 |
| **Transcript_360808** | ADP-ribosylation factor | -2.02 | 9.73E-04 |
| **Transcript_197796** | alpha-1 collagen | -3.43 | 3.20E-11 |
| **Transcript_198232** | alpha-1 collagen | -2.61 | 2.61E-08 |
| **Transcript_396043** | alpha-1 collagen | -2.18 | 6.97E-05 |
| **Transcript_345429** | alpha-amylase 4N | -2.34 | 2.95E-03 |
| **Transcript_410512** | alpha-crystallin B chain | -2.94 | 1.49E-03 |
| **Transcript_430725** | alpha-crystallin B chain | -2.03 | 1.93E-05 |
| **Transcript_195199** | angiopoietin-4-like | -2.43 | 9.40E-03 |
| **Transcript_211686** | arylsulfatase | -4.17 | 1.71E-03 |
| **Transcript_234345** | barH-like 2 homeobox protein | -3.23 | 7.50E-05 |
| **Transcript_374682** | bromodomain-containing protein 4-like\|dentin sialophosphoprotein | -2.12 | 6.69E-03 |
| **Transcript_203368** | calcitonin gene-related peptide type 1 receptor | -2.32 | 5.00E-05 |
| **Transcript_82280** | calcium-activated chloride channel regulator 1\|epithelial chloride channel protein | -2.04 | 4.83E-06 |
| **Transcript_454815** | calmodulin | -3.16 | 1.16E-02 |
| **Transcript_346536** | calmodulin | -2.13 | 5.15E-06 |
| **Transcript_60050** | cAMP-dependent protein kinase catalytic subunit 1\|catalytic subunit of cAMP-dependent histone kinase | -2.62 | 7.18E-05 |
| **Transcript_7357** | cardioacceleratory peptide receptor-like | -2.28 | 2.73E-02 |
| **Transcript_387252** | cathepsin Z | -8.84 | 1.47E-08 |
| **Transcript_443599** | cholecystokinin receptor type A | -2.23 | 1.66E-02 |
| **Transcript_198131** | cyclin-dependent kinase 20-like | -2.29 | 2.97E-05 |

| | | | |
|---|---|---|---|
| **Transcript_338163** | cysteine and glycine-rich protein 1\|cysteine and glycine-rich protein 2 | -2.87 | 2.40E-16 |
| **Transcript_53888** | cytochrome P450 27C1\|25-hydroxyvitamin D-1 alpha hydroxylase, mitochondrial\|probable cytochrome P450 49a1 | -2.45 | 2.73E-03 |
| **Transcript_179647** | D(1) dopamine receptor | -2.15 | 1.15E-02 |
| **Transcript_213703** | deleted in malignant brain tumors 1 protein | -3.53 | 2.76E-03 |
| **Transcript_321418** | deleted in malignant brain tumors 1 protein | -2.73 | 2.76E-03 |
| **Transcript_74981** | disintegrin and metalloproteinase domain-containing protein 23-like\|disintegrin and metalloproteinase domain-containing protein 12 | -2.32 | 8.72E-03 |
| **Transcript_455894** | DNA-directed RNA polymerases I, II, and III subunit RPABC3 | -2.59 | 9.85E-03 |
| **Transcript_44175** | E-selectin-like | -2.09 | 6.66E-03 |
| **Transcript_412294** | E3 ubiquitin-protein ligase TRIM56-like | -4.42 | 3.14E-02 |
| **Transcript_366986** | fibrinogen C domain-containing protein 1 | -2.20 | 3.17E-02 |
| **Transcript_298990** | ficolin-1-like | -4.27 | 5.65E-06 |
| **Transcript_366365** | ficolin-1-like | -2.21 | 1.60E-02 |
| **Transcript_259611** | ficolin-2 | -2.54 | 1.29E-02 |
| **Transcript_435128** | formin-J | -2.24 | 2.03E-05 |
| **Transcript_86447** | G-protein coupled receptor 54 | -2.30 | 3.74E-02 |
| **Transcript_380362** | glyceraldehyde-3-phosphate dehydrogenase | -3.84 | 1.53E-02 |
| **Transcript_205220** | glycine receptor subunit alpha-4 | -2.27 | 6.61E-12 |
| **Transcript_288548** | glycine-rich cell wall structural protein 1\|glycine-rich protein DOT1 | -2.14 | 3.14E-02 |
| **Transcript_305980** | golgin subfamily A member 6-like protein 22 | -2.25 | 1.60E-02 |
| **Transcript_264943** | GTP-binding protein Rit1-like\|GTP-binding protein Rit1 pseudogene | -2.07 | 4.07E-03 |
| **Transcript_281300** | hairy/enhancer-of-split related with YRPW motif protein\|uncharacterized protein LOC593175 | -2.17 | 2.97E-02 |
| **Transcript_281299** | heat shock cognate 71 kDa protein\|heat shock 70kDa protein 8 | -2.96 | 1.97E-04 |
| **Transcript_300969** | heat shock cognate 71 kDa protein\|heat shock 70kDa protein 8 | -2.37 | 8.25E-03 |
| **Transcript_200327** | histamine H2 receptor-like | -4.30 | 7.87E-04 |
| **Transcript_338151** | histidine triad nucleotide-binding protein 3 | -2.68 | 4.10E-03 |
| **Transcript_161417** | histone H3, embryonic | -3.59 | 1.41E-02 |
| **Transcript_276623** | histone H4 | -4.81 | 4.60E-02 |
| **Transcript_212508** | histone H4 | -3.12 | 3.75E-02 |
| **Transcript_339518** | homeobox protein Hox-A7 | -2.50 | 3.18E-02 |
| **Transcript_394760** | homeodomain protein | -4.44 | 2.56E-03 |
| **Transcript_436417** | IgGFc-binding protein\|Fc fragment of IgG binding protein\|zonadhesin | -2.19 | 6.67E-03 |
| **Transcript_374429** | ileal sodium/bile acid cotransporter-like | -4.01 | 8.60E-03 |
| **Transcript_279040** | kelch-like protein 2 | -2.18 | 3.64E-05 |
| **Transcript_219771** | Krueppel-like factor 13 | -2.29 | 2.33E-03 |

| | | | |
|---|---|---|---|
| **Transcript_376962** | lactase-phlorizin hydrolase | -2.76 | 3.76E-02 |
| **Transcript_274293** | lambda-crystallin homolog | -2.16 | 2.13E-05 |
| **Transcript_429112** | large neutral amino acids transporter small subunit 2 | -2.27 | 1.22E-05 |
| **Transcript_204036** | serine/threonine-protein kinase mos | -2.13 | 2.72E-04 |
| **Transcript_229121** | serine/threonine-protein kinase mos | -2.02 | 2.54E-03 |
| **Transcript_257994** | LOW QUALITY PROTEIN: MMP37-like protein, mitochondrial\|MMP37-like protein, mitochondrial | -2.04 | 4.07E-02 |
| **Transcript_391508** | LOW QUALITY PROTEIN: tubulin alpha-1 chain | -2.73 | 2.20E-08 |
| **Transcript_362387** | LOW QUALITY PROTEIN: tubulin alpha-1 chain | -2.36 | 6.65E-04 |
| **Transcript_420771** | LOW QUALITY PROTEIN: tubulin alpha-1C chain | -2.16 | 2.86E-02 |
| **Transcript_364074** | LOW QUALITY PROTEIN: uncharacterized protein LOC588722 | -3.12 | 4.00E-04 |
| **Transcript_339876** | LOW QUALITY PROTEIN: zinc finger protein 708-like | -3.37 | 1.48E-02 |
| **Transcript_273801** | metabotropic glutamate receptor 8 | -2.22 | 4.80E-05 |
| **Transcript_59717** | methylmalonic aciduria and homocystinuria type C protein homolog | -2.07 | 3.76E-03 |
| **Transcript_343882** | microfibril-associated glycoprotein 4 | -4.38 | 4.85E-03 |
| **Transcript_216921** | microfibril-associated glycoprotein 4 | -2.53 | 1.48E-04 |
| **Transcript_410262** | microfibril-associated glycoprotein 4-like | -4.02 | 2.18E-02 |
| **Transcript_179645** | microfibril-associated glycoprotein 4-like | -2.94 | 3.25E-03 |
| **Transcript_445086** | microfibril-associated glycoprotein 4-like | -2.41 | 1.99E-03 |
| **Transcript_429455** | microfibril-associated glycoprotein 4-like | -2.38 | 4.46E-03 |
| **Transcript_220874** | microfibril-associated glycoprotein 4\|tenascin-N | -9.00 | 5.41E-03 |
| **Transcript_362512** | microfibril-associated glycoprotein 4\|tenascin-N | -4.39 | 4.85E-03 |
| **Transcript_335205** | microfibril-associated glycoprotein 4\|tenascin-N | -2.89 | 8.65E-03 |
| **Transcript_367362** | microfibril-associated glycoprotein 4\|tenascin-N | -2.64 | 5.42E-03 |
| **Transcript_343707** | microfibril-associated glycoprotein 4\|tenascin-N | -2.25 | 3.28E-02 |
| **Transcript_365407** | microfibril-associated glycoprotein 4\|tenascin-N | -2.22 | 8.36E-06 |
| **Transcript_343323** | microfibril-associated glycoprotein 4\|tenascin-N | -2.11 | 9.52E-03 |
| **Transcript_264072** | monocarboxylate transporter 12\|retinol dehydrogenase 8\|Short-chain dehydrogenase/reductase SDR domain containing protein | -2.68 | 5.62E-03 |
| **Transcript_429832** | muscle-specific protein 20 | -2.01 | 3.32E-04 |
| **Transcript_414296** | NADPH oxidase 5 | -2.55 | 1.02E-03 |
| **Transcript_341092** | neurogenic differentiation factor 4 | -2.19 | 2.56E-02 |
| **Transcript_398720** | neuronal acetylcholine receptor subunit alpha-3 | -2.89 | 3.68E-13 |
| **Transcript_183115** | neurotrophin 5\|prepro-neurotrophin | -2.35 | 3.71E-07 |

| | | | |
|---|---|---|---|
| **Transcript_254653** | nucleolar GTP-binding protein 2 | -2.17 | 7.10E-05 |
| **Transcript_420284** | orexin receptor type 2 | -2.50 | 2.06E-04 |
| **Transcript_321671** | oxytocin receptor\|gonadotropin-releasing hormone receptor | -3.48 | 5.51E-04 |
| **Transcript_196171** | paired box protein Pax-2a | -2.30 | 3.24E-04 |
| **Transcript_335269** | peptidyl-prolyl cis-trans isomerase | -4.41 | 2.41E-03 |
| **Transcript_418613** | phospholipid scramblase 2\|phospholipid scramblase family member 5 | -2.09 | 6.80E-03 |
| **Transcript_391646** | PIN2/TERF1-interacting telomerase inhibitor 1 | -2.41 | 2.09E-04 |
| **Transcript_348719** | popeye domain-containing protein 3 | -2.49 | 8.96E-13 |
| **Transcript_208618** | potassium channel subfamily K member 9 | -2.68 | 4.58E-04 |
| **Transcript_297999** | probable cationic amino acid transporter\|putative cationic amino acid transporter\|solute carrier family 7 (orphan transporter), member 14 | -2.43 | 2.93E-03 |
| **Transcript_255002** | probable threonine protease PRSS50 | -2.87 | 9.99E-09 |
| **Transcript_217497** | protein giant | -2.12 | 1.54E-04 |
| **Transcript_303695** | protein PLANT CADMIUM RESISTANCE 3-like | -2.66 | 6.01E-04 |
| **Transcript_409861** | protein SSUH2 homolog | -2.28 | 1.53E-02 |
| **Transcript_380758** | protein SSUH2 homolog | -2.22 | 4.36E-05 |
| **Transcript_315415** | protein Tob1 | -2.75 | 1.53E-02 |
| **Transcript_405702** | putative uncharacterized protein CXorf58 | -2.53 | 9.26E-04 |
| **Transcript_483697** | rab3 GTPase | -2.08 | 1.81E-03 |
| **Transcript_426847** | ras-related protein ORAB-1 | -4.04 | 8.23E-03 |
| **Transcript_182361** | retinol dehydrogenase 8 | -2.15 | 1.46E-02 |
| **Transcript_269277** | rho GTPase-activating protein 25 | -2.73 | 7.41E-03 |
| **Transcript_367031** | ribosomal protein S14 | -3.32 | 1.75E-02 |
| **Transcript_350598** | ribosome biogenesis protein bop1-B | -2.08 | 4.29E-03 |
| **Transcript_373223** | RNA 3'-terminal phosphate cyclase-like protein | -2.05 | 3.54E-03 |
| **Transcript_253908** | RNA polymerase II subunit A C-terminal domain phosphatase SSU72 | -2.44 | 3.37E-05 |
| **Transcript_300509** | ryncolin-1-like | -3.08 | 4.21E-03 |
| **Transcript_370378** | serine/threonine-protein kinase mos | -2.03 | 3.42E-02 |
| **Transcript_381580** | serine/threonine-protein kinase NLK\|serine/threonine-protein kinase NLK2 | -2.27 | 3.18E-02 |
| **Transcript_285582** | short-chain collagen C4 | -2.84 | 5.23E-04 |
| **Transcript_285813** | splicing factor 3A subunit 2 | -2.31 | 1.66E-04 |
| **Transcript_309701** | sugar phosphate exchanger 3 | -2.12 | 3.13E-02 |
| **Transcript_179855** | sushi, von Willebrand factor type A, EGF and pentraxin domain-containing protein 1 | -2.49 | 3.74E-03 |
| **Transcript_236683** | synaptotagmin-15 | -2.28 | 2.41E-03 |
| **Transcript_275400** | synaptotagmin-17-like | -2.77 | 1.50E-02 |
| **Transcript_229301** | synaptotagmin-7 | -2.73 | 2.00E-06 |
| **Transcript_324193** | tubulin alpha chain | -3.36 | 3.81E-20 |
| **Transcript_332311** | tubulin alpha chain | -2.85 | 6.61E-12 |

| | | | |
|---|---|---|---|
| **Transcript_285898** | tubulin alpha chain | -2.15 | 3.71E-04 |
| **Transcript_49445** | tubulin alpha-1 chain | -3.02 | 2.93E-09 |
| **Transcript_327532** | tubulin alpha-1 chain | -2.96 | 1.46E-12 |
| **Transcript_285897** | tubulin alpha-1 chain | -2.41 | 1.12E-03 |
| **Transcript_229300** | tubulin alpha-1 chain | -2.04 | 1.20E-04 |
| **Transcript_207127** | tubulin alpha-1 chain\|tubulin alpha-1A chain | -2.23 | 2.67E-03 |
| **Transcript_306226** | tubulin alpha-1A chain | -5.73 | 2.00E-03 |
| **Transcript_354200** | tubulin alpha-1A chain | -4.12 | 8.00E-04 |
| **Transcript_89028** | tubulin alpha-1A chain | -3.17 | 1.17E-02 |
| **Transcript_367771** | tubulin alpha-1A chain | -3.12 | 4.06E-05 |
| **Transcript_362085** | tubulin alpha-1A chain | -2.62 | 1.04E-04 |
| **Transcript_262388** | tubulin alpha-1A chain | -2.56 | 7.67E-05 |
| **Transcript_377194** | tubulin alpha-1A chain | -2.54 | 1.27E-06 |
| **Transcript_222321** | tubulin alpha-1A chain | -2.05 | 5.62E-04 |
| **Transcript_412907** | tubulin alpha-1A chain | -2.04 | 7.70E-04 |
| **Transcript_323033** | tubulin alpha-2/alpha-4 chain | -2.83 | 1.99E-02 |
| **Transcript_367619** | tubulin beta chain | -3.62 | 3.14E-04 |
| **Transcript_277435** | tubulin beta chain | -2.20 | 2.18E-02 |
| **Transcript_230874** | tyrosine-protein phosphatase non-receptor type 9 | -2.10 | 4.46E-02 |
| **Transcript_196821** | ubiquitin | -2.88 | 2.92E-02 |
| **Transcript_433726** | UDP-glucuronosyltransferase 2C1\|UDP-glucuronosyltransferase 1-2\|UDP-glucuronosyltransferase 2A2 | -2.73 | 2.97E-02 |
| **Transcript_402138** | uncharacterized protein LOC100888048 | -3.85 | 4.20E-04 |
| **Transcript_355756** | uncharacterized protein LOC100888517 | -2.19 | 9.35E-05 |
| **Transcript_369230** | uncharacterized protein LOC100891189 | -2.61 | 8.60E-04 |
| **Transcript_40273** | uncharacterized protein LOC105446839 | -3.22 | 6.08E-03 |
| **Transcript_349228** | uncharacterized protein LOC576450 | -2.43 | 1.07E-07 |
| **Transcript_331800** | uncharacterized protein LOC591826 | -2.18 | 1.06E-02 |
| **Transcript_247803** | uncharacterized protein LOC592324 | -2.48 | 6.86E-03 |
| **Transcript_456195** | uncharacterized protein LOC753087\|185/333 D1 alpha\|185/333 D5 epsilon\|185/333 E6 alpha | -2.11 | 1.73E-04 |
| **Transcript_399035** | uncharacterized protein LOC756005\|protein of unknown function DUF2181 containing protein | -3.38 | 7.19E-08 |
| **Transcript_199904** | uncharacterized protein LOC756131 | -2.05 | 1.38E-04 |
| **Transcript_448557** | vasoactive intestinal polypeptide receptor 1\|parathyroid hormone 2 receptor | -2.04 | 4.20E-03 |
| **Transcript_339985** | von Willebrand factor A domain-containing protein 5B1 | -2.40 | 3.64E-04 |
| **Transcript_223195** | zinc finger SWIM domain-containing protein 8\|zinc finger SWIM domain-containing protein KIAA0913-like | -2.01 | 3.14E-02 |

*Highlighted transcripts in yellow are nervous system associated genes

Table S9. Differentially expressed transcription factors

| ID | Annotated name | Abbreviation | Log2FC | Adjusted p-value |
|---|---|---|---|---|
| **Transcript_319952** | Transcription factor SOX-4 | SOX4 | 4.50 | 9.30E-34 |
| **Transcript_263596** | Transcription factor BTF3 homolog 4 | BTF3L4 | 4.16 | 2.16E-11 |
| **Transcript_175325** | Thyroid transcription factor 1-associated protein 26 homolog | TAP26 | 3.00 | 2.98E-11 |
| **Transcript_290528** | Transcription factor VBP | TEF | 5.74 | 1.06E-06 |
| **Transcript_423204** | Forkhead transcription factor A | FOXA | -3.83 | 3.80E-05 |
| **Transcript_287026** | Transcription factor 25 | TCF25 | 1.77 | 1.29E-04 |
| **Transcript_443183** | Myc protein | MYC | 2.72 | 2.63E-04 |
| **Transcript_387814** | Transcription factor IIIA | GTF3A | 1.86 | 5.36E-04 |
| **Transcript_239737** | Pre-B-cell leukemia transcription factor 1 | PBX3 | 1.36 | 1.46E-03 |
| **Transcript_265397** | Transcription factor soxd1 | SOXD1 | 1.43 | 1.90E-03 |
| **Transcript_373546** | Nuclear transcription factor Y subunit gamma | NFYC | 1.35 | 2.20E-03 |
| **Transcript_244221** | Transcription factor AP-1 | AP-1 | -1.81 | 3.44E-03 |
| **Transcript_269278** | Transcription factor HES-4 | HES4 | -2.55 | 6.28E-03 |
| **Transcript_286791** | General transcription factor IIH subunit 1 | GTF2H1 | 1.27 | 1.13E-02 |
| **Transcript_220104** | Homeobox protein Hox-A10 | HOXA10 | 1.55 | 1.20E-02 |
| **Transcript_321671** | Paired box protein Pax-2a | PAX2A | 1.50 | 1.69E-02 |
| **Transcript_441452** | LIM homeobox transcription factor 1-beta | LMX1B | -1.61 | 1.83E-02 |
| **Transcript_304792** | ETS-related transcription factor Elf-3 | ELF3 | 1.33 | 2.47E-02 |
| **Transcript_211042** | Myelin transcription factor 1-like protein | MYT1L1 | 1.04 | 2.47E-02 |
| **Transcript_274240** | Beta-catenin | CTNNB | 1.03 | 2.96E-02 |
| **Transcript_224155** | Zinc-finger transcription factor Snail | SNAIL | 1.19 | 9.00E-02 |
| **Transcript_291169** | GATA transcription factor e | GATAE | -1.97 | 1.20E-01 |
| **Transcript_441904** | Transcription factor E2F5 | E2F5 | 1.01 | 1.21E-01 |
| **Transcript_39673** | Transcription factor AP-2-alpha | TFAP2A | 2.43 | 1.31E-01 |
| **Transcript_307462** | Homeobox transcription factor Nk1 | NK1 | -1.93 | 2.44E-01 |
| **Transcript_361564** | Forkhead transcription factor J1 | FOXJ1 | -1.08 | 3.06E-01 |
| **Transcript_107633** | LIM domain transcription factor LMO4-B | LMO4 | 1.23 | 3.40E-01 |
| **Transcript_76067** | Winged helix transcription factor Forkhead-1 | FKH1 | 1.05 | 3.58E-01 |
| **Transcript_168671** | Elongation factor 1 alpha | EEF1A1 | 7.32 | 8.38E-05 |
| **Transcript_345735** | Homeobox protein DLX-5\|homeobox protein Hox-B4 | DLX5 | 3.07 | 4.06E-02 |
| **Transcript_429867** | Homeobox protein Hmx | HMX | 3.59 | 8.86E-17 |
| **Transcript_279040** | Krueppel-like factor 13 | KLF13 | 3.21 | 1.17E-06 |
| **Transcript_433953** | Periodic tryptophan protein 1 homolog | PWP1 | 3.35 | 4.80E-12 |
| **Transcript_432532** | Rhombotin-2 | LMO2 | 2.95 | 1.48E-10 |
| **Transcript_236625** | Homeobox protein CDX-2 | CDX1 | -8.64 | 2.33E-03 |
| **Transcript_271521** | Fos-related antigen 1 | FOSL2 | -4.85 | 5.48E-03 |

| | | | | |
|---|---|---|---|---|
| **Transcript_438891** | Homeobox domain-containing protein | HLX | -4.10 | 1.38E-17 |
| **Transcript_173387** | Hepatocyte nuclear factor 4-gamma | HNF4G | -3.38 | 1.70E-03 |
| **Transcript_438353** | Homeobox protein EMX1 | EMX1 | -4.28 | 1.09E-03 |
| **Transcript_459460** | THAP domain-containing protein 5 | THAP4 | 2.86 | 1.08E-02 |
| **Transcript_433507** | Homeobox protein Mohawk-like | MKX | 2.61 | 7.15E-06 |
| **Transcript_284772** | TATA box-binding protein-associated factor RNA polymerase I subunit A | TAF1A | 2.54 | 2.14E-04 |
| **Transcript_189718** | Myb-binding protein 1A-like protein | MYBBP1A | 2.38 | 1.71E-05 |
| **Transcript_459696** | Myoneurin | MYNN | 2.11 | 4.88E-05 |
| **Transcript_268430** | Gastrula zinc finger protein xlcgf57.1 isoform X5 | ZNF | -3.06 | 8.46E-03 |
| **Transcript_247937** | Early growth response protein 1-B | EGR1 | -2.75 | 1.42E-04 |
| **Transcript_330100** | Basic leucine zipper domain, Maf-type | BSL78 | -2.45 | 5.85E-05 |

Table S10. Gene ontology analysis from day 1 downregulated genes using DAVID

| Category | Term | Count | PValue | List.Total | Pop.Hits | Pop.Total | Fold.Enrichment | Bonferroni | Benjamini | FDR | Description |
|---|---|---|---|---|---|---|---|---|---|---|---|
| BP | GO:0007155 | 5 | 9.67E-04 | 39 | 14 | 1094 | 10.02 | 0.31 | 0.31 | 1.33 | cell adhesion |
| BP | GO:0022610 | 5 | 1.29E-03 | 39 | 15 | 1094 | 9.35 | 0.38 | 0.22 | 1.76 | biological adhesion |
| BP | GO:0006082 | 7 | 1.15E-02 | 39 | 57 | 1094 | 3.44 | 0.99 | 0.58 | 14.81 | organic acid metabolic process |
| BP | GO:0044699 | 28 | 1.97E-02 | 39 | 586 | 1094 | 1.34 | 1.00 | 0.66 | 24.03 | single-organism process |
| BP | GO:0046394 | 4 | 3.34E-02 | 39 | 21 | 1094 | 5.34 | 1.00 | 0.80 | 37.47 | carboxylic acid biosynthetic process |
| BP | GO:0044763 | 26 | 3.49E-02 | 39 | 548 | 1094 | 1.33 | 1.00 | 0.77 | 38.79 | single-organism cellular process |
| BP | GO:0002376 | 3 | 4.44E-02 | 39 | 10 | 1094 | 8.42 | 1.00 | 0.79 | 46.60 | immune system process |
| BP | GO:0032501 | 5 | 4.98E-02 | 39 | 41 | 1094 | 3.42 | 1.00 | 0.80 | 50.65 | multicellular organismal process |
| BP | GO:0044283 | 4 | 6.36E-02 | 39 | 27 | 1094 | 4.16 | 1.00 | 0.85 | 59.72 | small molecule biosynthetic process |
| CC | GO:0031224 | 26 | 6.01E-03 | 47 | 445 | 1253 | 1.56 | 0.42 | 0.42 | 6.35 | intrinsic component of membrane |
| CC | GO:0016020 | 29 | 7.78E-03 | 47 | 531 | 1253 | 1.46 | 0.50 | 0.30 | 8.15 | membrane |
| CC | GO:0016021 | 25 | 1.32E-02 | 47 | 444 | 1253 | 1.50 | 0.70 | 0.33 | 13.43 | integral component of membrane |
| CC | GO:0044425 | 26 | 2.30E-02 | 47 | 489 | 1253 | 1.42 | 0.88 | 0.41 | 22.35 | membrane part |
| MF | GO:0016798 | 5 | 1.02E-03 | 54 | 12 | 1277 | 9.85 | 0.16 | 0.16 | 1.24 | hydrolase activity, acting on glycosyl bonds |
| MF | GO:0005044 | 4 | 3.26E-03 | 54 | 8 | 1277 | 11.82 | 0.43 | 0.24 | 3.91 | scavenger receptor activity |
| MF | GO:0038024 | 4 | 3.26E-03 | 54 | 8 | 1277 | 11.82 | 0.43 | 0.24 | 3.91 | cargo receptor activity |
| MF | GO:0004872 | 8 | 1.19E-02 | 54 | 62 | 1277 | 3.05 | 0.87 | 0.49 | 13.56 | receptor activity |
| MF | GO:0060089 | 8 | 1.19E-02 | 54 | 62 | 1277 | 3.05 | 0.87 | 0.49 | 13.56 | molecular transducer activity |
| MF | GO:0008237 | 4 | 1.78E-02 | 54 | 14 | 1277 | 6.76 | 0.95 | 0.53 | 19.65 | metallopeptidase activity |
| MF | GO:0003824 | 34 | 2.42E-02 | 54 | 614 | 1277 | 1.31 | 0.98 | 0.57 | 25.83 | catalytic activity |
| MF | GO:0016491 | 10 | 2.50E-02 | 54 | 105 | 1277 | 2.25 | 0.99 | 0.51 | 26.58 | oxidoreductase activity |
| MF | GO:0004553 | 3 | 5.05E-02 | 54 | 9 | 1277 | 7.88 | 1.00 | 0.72 | 46.84 | hydrolase activity, hydrolyzing O-glycosyl compounds |
| MF | GO:0005509 | 5 | 8.00E-02 | 54 | 40 | 1277 | 2.96 | 1.00 | 0.83 | 63.85 | calcium ion binding |
| MF | GO:0016805 | 2 | 8.13E-02 | 54 | 2 | 1277 | 23.65 | 1.00 | 0.80 | 64.47 | dipeptidase activity |
| MF | GO:0003796 | 2 | 8.13E-02 | 54 | 2 | 1277 | 23.65 | 1.00 | 0.80 | 64.47 | lysozyme activity |
| MF | GO:0004497 | 3 | 9.84E-02 | 54 | 13 | 1277 | 5.46 | 1.00 | 0.83 | 71.75 | monooxygenase activity |

Table S11. Gene ontology analysis from day 1 upregulated genes using DAVID

| Category | Term | Count | PValue | List.Total | Pop.Hits | Pop.Total | Fold.Enrichment | Bonferroni | Benjamini | FDR | Description |
|---|---|---|---|---|---|---|---|---|---|---|---|
| BP | GO:0022613 | 23 | 1.28E-08 | 99 | 68 | 1094 | 3.74 | 1.02E-05 | 1.02E-05 | 1.97E-05 | ribonucleoprotein complex biogenesis |
| BP | GO:0016072 | 12 | 1.52E-05 | 99 | 29 | 1094 | 4.57 | 1.21E-02 | 4.05E-03 | 2.34E-02 | rRNA metabolic process |
| BP | GO:0034660 | 15 | 2.45E-05 | 99 | 47 | 1094 | 3.53 | 1.94E-02 | 4.87E-03 | 3.76E-02 | ncRNA metabolic process |
| BP | GO:0071840 | 38 | 3.45E-04 | 99 | 250 | 1094 | 1.68 | 2.41E-01 | 4.49E-02 | 5.28E-01 | cellular component organization or biogenesis |
| BP | GO:0010467 | 41 | 5.40E-04 | 99 | 284 | 1094 | 1.60 | 3.50E-01 | 5.97E-02 | 8.25E-01 | gene expression |
| BP | GO:0044085 | 25 | 6.48E-04 | 99 | 140 | 1094 | 1.97 | 4.04E-01 | 6.27E-02 | 9.90E-01 | cellular component biogenesis |
| BP | GO:0006413 | 6 | 9.15E-04 | 99 | 10 | 1094 | 6.63 | 5.18E-01 | 7.80E-02 | 1.40E+00 | translational initiation |
| BP | GO:0034641 | 53 | 1.09E-03 | 99 | 416 | 1094 | 1.41 | 5.83E-01 | 8.37E-02 | 1.67E+00 | cellular nitrogen compound metabolic process |
| BP | GO:0006412 | 17 | 1.49E-03 | 99 | 82 | 1094 | 2.29 | 6.96E-01 | 1.02E-01 | 2.26E+00 | translation |
| BP | GO:0006807 | 54 | 2.72E-03 | 99 | 441 | 1094 | 1.35 | 8.86E-01 | 1.44E-01 | 4.09E+00 | nitrogen compound metabolic process |
| BP | GO:1990542 | 5 | 3.20E-03 | 99 | 8 | 1094 | 6.91 | 9.23E-01 | 1.57E-01 | 4.81E+00 | mitochondrial transmembrane transport |
| BP | GO:0006396 | 16 | 3.23E-03 | 99 | 80 | 1094 | 2.21 | 9.24E-01 | 1.49E-01 | 4.85E+00 | RNA processing |
| BP | GO:1901566 | 24 | 4.64E-03 | 99 | 151 | 1094 | 1.76 | 9.76E-01 | 1.86E-01 | 6.89E+00 | organonitrogen compound biosynthetic process |
| BP | GO:0044260 | 56 | 4.67E-03 | 99 | 472 | 1094 | 1.31 | 9.76E-01 | 1.78E-01 | 6.93E+00 | cellular macromolecule metabolic process |
| BP | GO:1902582 | 6 | 5.43E-03 | 99 | 14 | 1094 | 4.74 | 9.87E-01 | 1.95E-01 | 8.01E+00 | single-organism intracellular transport |
| BP | GO:0043603 | 18 | 6.25E-03 | 99 | 102 | 1094 | 1.95 | 9.93E-01 | 2.04E-01 | 9.18E+00 | cellular amide metabolic process |
| BP | GO:0017038 | 5 | 8.34E-03 | 99 | 10 | 1094 | 5.53 | 9.99E-01 | 2.43E-01 | 1.21E+01 | protein import |
| BP | GO:0006839 | 5 | 8.34E-03 | 99 | 10 | 1094 | 5.53 | 9.99E-01 | 2.43E-01 | 1.21E+01 | mitochondrial transport |
| BP | GO:0043170 | 56 | 8.50E-03 | 99 | 483 | 1094 | 1.28 | 9.99E-01 | 2.38E-01 | 1.23E+01 | macromolecule metabolic process |
| BP | GO:0007005 | 8 | 1.12E-02 | 99 | 29 | 1094 | 3.05 | 1.00E+00 | 2.93E-01 | 1.59E+01 | mitochondrion organization |
| BP | GO:0071826 | 8 | 1.12E-02 | 99 | 29 | 1094 | 3.05 | 1.00E+00 | 2.93E-01 | 1.59E+01 | ribonucleoprotein complex subunit organization |
| BP | GO:0071806 | 4 | 1.87E-02 | 99 | 7 | 1094 | 6.31 | 1.00E+00 | 4.16E-01 | 2.52E+01 | protein transmembrane transport |
| BP | GO:0051246 | 10 | 1.96E-02 | 99 | 47 | 1094 | 2.35 | 1.00E+00 | 4.20E-01 | 2.62E+01 | regulation of protein metabolic process |
| BP | GO:0016070 | 28 | 2.24E-02 | 99 | 210 | 1094 | 1.47 | 1.00E+00 | 4.53E-01 | 2.94E+01 | RNA metabolic process |
| BP | GO:0070585 | 4 | 2.80E-02 | 99 | 8 | 1094 | 5.53 | 1.00E+00 | 5.19E-01 | 3.53E+01 | protein localization to mitochondrion |
| BP | GO:0044267 | 32 | 3.19E-02 | 99 | 256 | 1094 | 1.38 | 1.00E+00 | 5.55E-01 | 3.92E+01 | cellular protein metabolic process |
| BP | GO:0044271 | 32 | 3.36E-02 | 99 | 257 | 1094 | 1.38 | 1.00E+00 | 5.63E-01 | 4.09E+01 | cellular nitrogen compound biosynthetic process |
| BP | GO:0032268 | 9 | 3.65E-02 | 99 | 44 | 1094 | 2.26 | 1.00E+00 | 5.82E-01 | 4.35E+01 | regulation of cellular protein metabolic process |
| BP | GO:0090304 | 31 | 4.15E-02 | 99 | 251 | 1094 | 1.36 | 1.00E+00 | 6.20E-01 | 4.78E+01 | nucleic acid metabolic process |
| BP | GO:0044237 | 66 | 4.44E-02 | 99 | 632 | 1094 | 1.15 | 1.00E+00 | 6.25E-01 | 5.02E+01 | cellular metabolic process |
| BP | GO:0006457 | 6 | 4.77E-02 | 99 | 23 | 1094 | 2.88 | 1.00E+00 | 6.41E-01 | 5.27E+01 | protein folding |
| BP | GO:1901564 | 25 | 4.86E-02 | 99 | 194 | 1094 | 1.42 | 1.00E+00 | 6.39E-01 | 5.34E+01 | organonitrogen compound metabolic process |
| BP | GO:0046483 | 38 | 4.97E-02 | 99 | 327 | 1094 | 1.28 | 1.00E+00 | 6.38E-01 | 5.43E+01 | heterocycle metabolic process |
| BP | GO:0019538 | 32 | 5.01E-02 | 99 | 265 | 1094 | 1.33 | 1.00E+00 | 6.32E-01 | 5.45E+01 | protein metabolic process |
| BP | GO:0006139 | 37 | 5.07E-02 | 99 | 317 | 1094 | 1.29 | 1.00E+00 | 6.28E-01 | 5.50E+01 | nucleobase-containing compound metabolic process |
| BP | GO:0010608 | 4 | 5.26E-02 | 99 | 10 | 1094 | 4.42 | 1.00E+00 | 6.33E-01 | 5.63E+01 | posttranscriptional regulation of gene expression |
| BP | GO:0071704 | 66 | 5.62E-02 | 99 | 638 | 1094 | 1.14 | 1.00E+00 | 6.50E-01 | 5.89E+01 | organic substance metabolic process |
| BP | GO:0006725 | 38 | 5.66E-02 | 99 | 330 | 1094 | 1.27 | 1.00E+00 | 6.44E-01 | 5.91E+01 | cellular aromatic compound metabolic process |
| BP | GO:0044238 | 63 | 6.62E-02 | 99 | 608 | 1094 | 1.15 | 1.00E+00 | 6.88E-01 | 6.51E+01 | primary metabolic process |
| BP | GO:0000469 | 3 | 6.64E-02 | 99 | 5 | 1094 | 6.63 | 1.00E+00 | 6.81E-01 | 6.52E+01 | cleavage involved in rRNA processing |
| BP | GO:0051345 | 3 | 6.64E-02 | 99 | 5 | 1094 | 6.63 | 1.00E+00 | 6.81E-01 | 6.52E+01 | positive regulation of hydrolase activity |

| | | | | | | | | | | | |
|---|---|---|---|---|---|---|---|---|---|---|---|
| BP | GO:0009894 | 4 | 6.77E-02 | 99 | 11 | 1094 | 4.02 | 1.00E+00 | 6.81E-01 | 6.59E+01 | regulation of catabolic process |
| BP | GO:0034248 | 4 | 6.77E-02 | 99 | 11 | 1094 | 4.02 | 1.00E+00 | 6.81E-01 | 6.59E+01 | regulation of cellular amide metabolic process |
| BP | GO:1901360 | 38 | 6.97E-02 | 99 | 335 | 1094 | 1.25 | 1.00E+00 | 6.77E-01 | 6.70E+01 | organic cyclic compound metabolic process |
| BP | GO:0090501 | 3 | 9.39E-02 | 99 | 6 | 1094 | 5.53 | 1.00E+00 | 7.80E-01 | 7.80E+01 | RNA phosphodiester bond hydrolysis |
| BP | GO:0000470 | 3 | 9.39E-02 | 99 | 6 | 1094 | 5.53 | 1.00E+00 | 7.80E-01 | 7.80E+01 | maturation of LSU-rRNA |
| BP | GO:0032543 | 3 | 9.39E-02 | 99 | 6 | 1094 | 5.53 | 1.00E+00 | 7.80E-01 | 7.80E+01 | mitochondrial translation |
| CC | GO:0005730 | 13 | 1.67E-07 | 108 | 25 | 1253 | 6.03 | 3.48E-05 | 3.48E-05 | 2.11E-04 | nucleolus |
| CC | GO:0031974 | 25 | 8.06E-06 | 108 | 113 | 1253 | 2.57 | 1.68E-03 | 8.38E-04 | 1.02E-02 | membrane-enclosed lumen |
| CC | GO:0030529 | 25 | 1.80E-05 | 108 | 118 | 1253 | 2.46 | 3.74E-03 | 1.25E-03 | 2.27E-02 | intracellular ribonucleoprotein complex |
| CC | GO:1990904 | 25 | 1.80E-05 | 108 | 118 | 1253 | 2.46 | 3.74E-03 | 1.25E-03 | 2.27E-02 | ribonucleoprotein complex |
| CC | GO:0030684 | 8 | 9.92E-05 | 108 | 15 | 1253 | 6.19 | 2.04E-02 | 5.14E-03 | 1.25E-01 | preribosome |
| CC | GO:0032991 | 47 | 8.50E-04 | 108 | 363 | 1253 | 1.50 | 1.62E-01 | 2.50E-02 | 1.07E+00 | macromolecular complex |
| CC | GO:0044428 | 22 | 1.15E-03 | 108 | 125 | 1253 | 2.04 | 2.13E-01 | 2.94E-02 | 1.44E+00 | nuclear part |
| CC | GO:0043228 | 28 | 1.70E-03 | 108 | 183 | 1253 | 1.78 | 2.99E-01 | 3.86E-02 | 2.13E+00 | non-membrane-bounded organelle |
| CC | GO:0043232 | 28 | 1.70E-03 | 108 | 183 | 1253 | 1.78 | 2.99E-01 | 3.86E-02 | 2.13E+00 | intracellular non-membrane-bounded organelle |
| CC | GO:0044452 | 5 | 2.69E-03 | 108 | 8 | 1253 | 7.25 | 4.29E-01 | 5.45E-02 | 3.34E+00 | nucleolar part |
| CC | GO:0070993 | 4 | 5.34E-03 | 108 | 5 | 1253 | 9.28 | 6.71E-01 | 9.62E-02 | 6.53E+00 | translation preinitiation complex |
| CC | GO:0033290 | 4 | 5.34E-03 | 108 | 5 | 1253 | 9.28 | 6.71E-01 | 9.62E-02 | 6.53E+00 | eukaryotic 48S preinitiation complex |
| CC | GO:0005622 | 86 | 1.03E-02 | 108 | 869 | 1253 | 1.15 | 8.83E-01 | 1.64E-01 | 1.22E+01 | intracellular |
| CC | GO:0005762 | 4 | 1.64E-02 | 108 | 7 | 1253 | 6.63 | 9.68E-01 | 2.33E-01 | 1.89E+01 | mitochondrial large ribosomal subunit |
| CC | GO:0022624 | 4 | 1.64E-02 | 108 | 7 | 1253 | 6.63 | 9.68E-01 | 2.33E-01 | 1.89E+01 | proteasome accessory complex |
| CC | GO:0098798 | 6 | 2.27E-02 | 108 | 20 | 1253 | 3.48 | 9.92E-01 | 2.89E-01 | 2.51E+01 | mitochondrial protein complex |
| CC | GO:0044424 | 80 | 2.45E-02 | 108 | 812 | 1253 | 1.14 | 9.94E-01 | 2.91E-01 | 2.69E+01 | intracellular part |
| CC | GO:0043234 | 32 | 2.72E-02 | 108 | 264 | 1253 | 1.41 | 9.97E-01 | 3.01E-01 | 2.93E+01 | protein complex |
| CC | GO:0005634 | 36 | 2.96E-02 | 108 | 308 | 1253 | 1.36 | 9.98E-01 | 3.07E-01 | 3.15E+01 | nucleus |
| CC | GO:0005623 | 91 | 3.12E-02 | 108 | 959 | 1253 | 1.10 | 9.99E-01 | 3.07E-01 | 3.30E+01 | cell |
| CC | GO:0005852 | 3 | 3.87E-02 | 108 | 4 | 1253 | 8.70 | 1.00E+00 | 3.36E-01 | 3.92E+01 | eukaryotic translation initiation factor 3 complex |
| CC | GO:0005615 | 6 | 5.52E-02 | 108 | 25 | 1253 | 2.78 | 1.00E+00 | 4.15E-01 | 5.12E+01 | extracellular space |
| CC | GO:0005732 | 3 | 6.09E-02 | 108 | 5 | 1253 | 6.96 | 1.00E+00 | 4.33E-01 | 5.47E+01 | small nucleolar ribonucleoprotein complex |
| CC | GO:0005739 | 15 | 6.72E-02 | 108 | 108 | 1253 | 1.61 | 1.00E+00 | 4.53E-01 | 5.84E+01 | mitochondrion |
| CC | GO:0044422 | 40 | 8.38E-02 | 108 | 377 | 1253 | 1.23 | 1.00E+00 | 5.03E-01 | 6.68E+01 | organelle part |
| CC | GO:0032040 | 3 | 8.63E-02 | 108 | 6 | 1253 | 5.80 | 1.00E+00 | 5.01E-01 | 6.80E+01 | small-subunit processome |
| CC | GO:0005758 | 3 | 8.63E-02 | 108 | 6 | 1253 | 5.80 | 1.00E+00 | 5.01E-01 | 6.80E+01 | mitochondrial intermembrane space |
| CC | GO:0031970 | 3 | 8.63E-02 | 108 | 6 | 1253 | 5.80 | 1.00E+00 | 5.01E-01 | 6.80E+01 | organelle envelope lumen |
| MF | GO:0003723 | 21 | 1.85E-06 | 98 | 86 | 1277 | 3.18 | 4.64E-04 | 4.64E-04 | 2.40E-03 | RNA binding |
| MF | GO:0008135 | 10 | 8.01E-06 | 98 | 21 | 1277 | 6.21 | 2.01E-03 | 1.00E-03 | 1.04E-02 | translation factor activity, RNA binding |
| MF | GO:0003743 | 7 | 4.71E-04 | 98 | 15 | 1277 | 6.08 | 1.11E-01 | 3.86E-02 | 6.10E-01 | translation initiation factor activity |
| MF | GO:0008565 | 4 | 1.80E-02 | 98 | 8 | 1277 | 6.52 | 9.89E-01 | 6.80E-01 | 2.10E+01 | protein transporter activity |
| MF | GO:0003676 | 29 | 4.59E-02 | 98 | 274 | 1277 | 1.38 | 1.00E+00 | 9.05E-01 | 4.57E+01 | nucleic acid binding |
| MF | GO:1901363 | 48 | 5.68E-02 | 98 | 515 | 1277 | 1.21 | 1.00E+00 | 9.13E-01 | 5.32E+01 | heterocyclic compound binding |
| MF | GO:0097159 | 48 | 6.47E-02 | 98 | 519 | 1277 | 1.21 | 1.00E+00 | 9.09E-01 | 5.81E+01 | organic cyclic compound binding |
| MF | GO:0034062 | 3 | 7.00E-02 | 98 | 6 | 1277 | 6.52 | 1.00E+00 | 8.98E-01 | 6.11E+01 | RNA polymerase activity |
| MF | GO:0003899 | 3 | 7.00E-02 | 98 | 6 | 1277 | 6.52 | 1.00E+00 | 8.98E-01 | 6.11E+01 | DNA-directed RNA polymerase activity |
| MF | GO:0016779 | 4 | 8.38E-02 | 98 | 14 | 1277 | 3.72 | 1.00E+00 | 9.13E-01 | 6.79E+01 | nucleotidyltransferase activity |

Table S12. Gene ontology analysis from day 3 downregulated genes using DAVID

| Category | Term | Count | PValue | List.Total | Pop.Hits | Pop.Total | Fold.Enrichment | Bonferroni | Benjamini | FDR | Description |
|---|---|---|---|---|---|---|---|---|---|---|---|
| BP | GO:0099537 | 3 | 3.22E-02 | 33 | 10 | 1094 | 9.95 | 1.00 | 1.00 | 35.00 | trans-synaptic signaling |
| BP | GO:0098916 | 3 | 3.22E-02 | 33 | 10 | 1094 | 9.95 | 1.00 | 1.00 | 35.00 | anterograde trans-synaptic signaling |
| BP | GO:0099536 | 3 | 3.22E-02 | 33 | 10 | 1094 | 9.95 | 1.00 | 1.00 | 35.00 | synaptic signaling |
| BP | GO:0007268 | 3 | 3.22E-02 | 33 | 10 | 1094 | 9.95 | 1.00 | 1.00 | 35.00 | chemical synaptic transmission |
| BP | GO:0007186 | 4 | 4.13E-02 | 33 | 27 | 1094 | 4.91 | 1.00 | 1.00 | 42.55 | G-protein coupled receptor signaling pathway |
| BP | GO:0007154 | 10 | 5.60E-02 | 33 | 175 | 1094 | 1.89 | 1.00 | 0.99 | 53.11 | cell communication |
| BP | GO:0044699 | 23 | 5.70E-02 | 33 | 586 | 1094 | 1.30 | 1.00 | 0.98 | 53.80 | single-organism process |
| BP | GO:0007155 | 3 | 6.06E-02 | 33 | 14 | 1094 | 7.10 | 1.00 | 0.97 | 56.04 | cell adhesion |
| BP | GO:0022610 | 3 | 6.87E-02 | 33 | 15 | 1094 | 6.63 | 1.00 | 0.96 | 60.76 | biological adhesion |
| CC | GO:0031224 | 33 | 1.77E-06 | 47 | 445 | 1253 | 1.98 | 0.00 | 0.00 | 0.00 | intrinsic component of membrane |
| CC | GO:0016021 | 32 | 7.07E-06 | 47 | 444 | 1253 | 1.92 | 0.00 | 0.00 | 0.01 | integral component of membrane |
| CC | GO:0016020 | 35 | 1.02E-05 | 47 | 531 | 1253 | 1.76 | 0.00 | 0.00 | 0.01 | membrane |
| CC | GO:0031226 | 12 | 1.35E-05 | 47 | 67 | 1253 | 4.77 | 0.00 | 0.00 | 0.02 | intrinsic component of plasma membrane |
| CC | GO:0005887 | 12 | 1.35E-05 | 47 | 67 | 1253 | 4.77 | 0.00 | 0.00 | 0.02 | integral component of plasma membrane |
| CC | GO:0044425 | 33 | 1.91E-05 | 47 | 489 | 1253 | 1.80 | 0.00 | 0.00 | 0.02 | membrane part |
| CC | GO:0071944 | 14 | 3.05E-05 | 47 | 101 | 1253 | 3.70 | 0.00 | 0.00 | 0.04 | cell periphery |
| CC | GO:0044459 | 12 | 5.36E-05 | 47 | 77 | 1253 | 4.15 | 0.01 | 0.00 | 0.06 | plasma membrane part |
| CC | GO:0005886 | 13 | 1.11E-04 | 47 | 98 | 1253 | 3.54 | 0.02 | 0.00 | 0.13 | plasma membrane |
| CC | GO:0005576 | 5 | 4.71E-02 | 47 | 38 | 1253 | 3.51 | 1.00 | 0.58 | 44.16 | extracellular region |
| MF | GO:0060089 | 12 | 2.23E-05 | 54 | 62 | 1277 | 4.58 | 0.00 | 0.00 | 0.03 | molecular transducer activity |
| MF | GO:0004872 | 12 | 2.23E-05 | 54 | 62 | 1277 | 4.58 | 0.00 | 0.00 | 0.03 | receptor activity |
| MF | GO:0004871 | 8 | 1.19E-02 | 54 | 62 | 1277 | 3.05 | 0.89 | 0.31 | 13.74 | signal transducer activity |
| MF | GO:0016491 | 10 | 2.50E-02 | 54 | 105 | 1277 | 2.25 | 0.99 | 0.49 | 26.90 | oxidoreductase activity |
| MF | GO:0005044 | 3 | 4.03E-02 | 54 | 8 | 1277 | 8.87 | 1.00 | 0.61 | 39.90 | scavenger receptor activity |
| MF | GO:0038024 | 3 | 4.03E-02 | 54 | 8 | 1277 | 8.87 | 1.00 | 0.61 | 39.90 | cargo receptor activity |
| MF | GO:1901618 | 2 | 8.13E-02 | 54 | 2 | 1277 | 23.65 | 1.00 | 0.83 | 64.99 | organic hydroxy compound transmembrane transporter activity |
| MF | GO:0052689 | 3 | 8.55E-02 | 54 | 12 | 1277 | 5.91 | 1.00 | 0.81 | 66.91 | carboxylic ester hydrolase activity |
| MF | GO:0004497 | 3 | 9.84E-02 | 54 | 13 | 1277 | 5.46 | 1.00 | 0.82 | 72.26 | monooxygenase activity |

Table S13. Gene ontology analysis from day 3 Upregulated genes using DAVID

| Category | Term | Count | PValue | List.Total | Pop.Hits | Pop.Total | Fold.Enrichment | Bonferroni | Benjamini | FDR | Description |
|---|---|---|---|---|---|---|---|---|---|---|---|
| BP | GO:0051246 | 9 | 7.48E-04 | 50 | 47 | 1094 | 4.19 | 0.35 | 0.35 | 1.10 | regulation of protein metabolic process |
| BP | GO:0032268 | 8 | 2.51E-03 | 50 | 44 | 1094 | 3.98 | 0.77 | 0.52 | 3.63 | regulation of cellular protein metabolic process |
| BP | GO:0006396 | 10 | 6.99E-03 | 50 | 80 | 1094 | 2.74 | 0.98 | 0.64 | 9.80 | RNA processing |
| BP | GO:0034470 | 7 | 9.04E-03 | 50 | 42 | 1094 | 3.65 | 0.99 | 0.65 | 12.50 | ncRNA processing |
| BP | GO:0009894 | 4 | 1.08E-02 | 50 | 11 | 1094 | 7.96 | 1.00 | 0.65 | 14.78 | regulation of catabolic process |
| BP | GO:0034660 | 7 | 1.56E-02 | 50 | 47 | 1094 | 3.26 | 1.00 | 0.68 | 20.61 | ncRNA metabolic process |
| BP | GO:0006457 | 5 | 1.68E-02 | 50 | 23 | 1094 | 4.76 | 1.00 | 0.67 | 22.02 | protein folding |
| BP | GO:0000470 | 3 | 2.63E-02 | 50 | 6 | 1094 | 10.94 | 1.00 | 0.79 | 32.40 | maturation of LSU-rRNA |
| BP | GO:0022613 | 8 | 2.77E-02 | 50 | 68 | 1094 | 2.57 | 1.00 | 0.77 | 33.86 | ribonucleoprotein complex biogenesis |
| BP | GO:0044260 | 29 | 3.08E-02 | 50 | 472 | 1094 | 1.34 | 1.00 | 0.78 | 36.91 | cellular macromolecule metabolic process |
| BP | GO:0071806 | 3 | 3.57E-02 | 50 | 7 | 1094 | 9.38 | 1.00 | 0.80 | 41.45 | protein transmembrane transport |
| BP | GO:0065002 | 3 | 3.57E-02 | 50 | 7 | 1094 | 9.38 | 1.00 | 0.80 | 41.45 | intracellular protein transmembrane transport |
| BP | GO:0042273 | 4 | 3.69E-02 | 50 | 17 | 1094 | 5.15 | 1.00 | 0.79 | 42.46 | ribosomal large subunit biogenesis |
| BP | GO:0043170 | 29 | 4.25E-02 | 50 | 483 | 1094 | 1.31 | 1.00 | 0.79 | 47.24 | macromolecule metabolic process |
| BP | GO:0044267 | 18 | 4.52E-02 | 50 | 256 | 1094 | 1.54 | 1.00 | 0.79 | 49.33 | cellular protein metabolic process |
| BP | GO:0006626 | 3 | 4.63E-02 | 50 | 8 | 1094 | 8.21 | 1.00 | 0.78 | 50.21 | protein targeting to mitochondrion |
| BP | GO:0009451 | 4 | 4.94E-02 | 50 | 19 | 1094 | 4.61 | 1.00 | 0.79 | 52.50 | RNA modification |
| BP | GO:0010467 | 19 | 5.89E-02 | 50 | 284 | 1094 | 1.46 | 1.00 | 0.81 | 59.08 | gene expression |
| BP | GO:0019538 | 18 | 6.08E-02 | 50 | 265 | 1094 | 1.49 | 1.00 | 0.81 | 60.25 | protein metabolic process |
| BP | GO:0048522 | 6 | 6.30E-02 | 50 | 49 | 1094 | 2.68 | 1.00 | 0.81 | 61.61 | positive regulation of cellular process |
| BP | GO:0048518 | 6 | 6.30E-02 | 50 | 49 | 1094 | 2.68 | 1.00 | 0.81 | 61.61 | positive regulation of biological process |
| BP | GO:0006412 | 8 | 6.70E-02 | 50 | 82 | 1094 | 2.13 | 1.00 | 0.80 | 63.95 | translation |
| BP | GO:0051336 | 3 | 7.03E-02 | 50 | 10 | 1094 | 6.56 | 1.00 | 0.80 | 65.77 | regulation of hydrolase activity |
| BP | GO:0042981 | 3 | 7.03E-02 | 50 | 10 | 1094 | 6.56 | 1.00 | 0.80 | 65.77 | regulation of apoptotic process |
| BP | GO:0006839 | 3 | 7.03E-02 | 50 | 10 | 1094 | 6.56 | 1.00 | 0.80 | 65.77 | mitochondrial transport |
| BP | GO:0010608 | 3 | 7.03E-02 | 50 | 10 | 1094 | 6.56 | 1.00 | 0.80 | 65.77 | posttranscriptional regulation of gene expression |
| BP | GO:0017038 | 3 | 7.03E-02 | 50 | 10 | 1094 | 6.56 | 1.00 | 0.80 | 65.77 | protein import |
| BP | GO:0006413 | 3 | 7.03E-02 | 50 | 10 | 1094 | 6.56 | 1.00 | 0.80 | 65.77 | translational initiation |
| BP | GO:0034641 | 25 | 7.27E-02 | 50 | 416 | 1094 | 1.31 | 1.00 | 0.79 | 67.04 | cellular nitrogen compound metabolic process |
| BP | GO:0043603 | 9 | 7.74E-02 | 50 | 102 | 1094 | 1.93 | 1.00 | 0.80 | 69.42 | cellular amide metabolic process |
| BP | GO:0006807 | 26 | 7.95E-02 | 50 | 441 | 1094 | 1.29 | 1.00 | 0.80 | 70.43 | nitrogen compound metabolic process |
| BP | GO:0034248 | 3 | 8.35E-02 | 50 | 11 | 1094 | 5.97 | 1.00 | 0.81 | 72.27 | regulation of cellular amide metabolic process |
| BP | GO:0044085 | 11 | 8.43E-02 | 50 | 140 | 1094 | 1.72 | 1.00 | 0.80 | 72.60 | cellular component biogenesis |
| BP | GO:0030155 | 2 | 8.76E-02 | 50 | 2 | 1094 | 21.88 | 1.00 | 0.80 | 74.04 | regulation of cell adhesion |
| BP | GO:0030334 | 2 | 8.76E-02 | 50 | 2 | 1094 | 21.88 | 1.00 | 0.80 | 74.04 | regulation of cell migration |
| BP | GO:0032231 | 2 | 8.76E-02 | 50 | 2 | 1094 | 21.88 | 1.00 | 0.80 | 74.04 | regulation of actin filament bundle assembly |
| CC | GO:0030529 | 16 | 1.24E-04 | 58 | 118 | 1253 | 2.93 | 0.02 | 0.02 | 0.15 | intracellular ribonucleoprotein complex |
| CC | GO:1990904 | 16 | 1.24E-04 | 58 | 118 | 1253 | 2.93 | 0.02 | 0.02 | 0.15 | ribonucleoprotein complex |
| CC | GO:0005615 | 7 | 6.15E-04 | 58 | 25 | 1253 | 6.05 | 0.09 | 0.05 | 0.74 | extracellular space |

| | | | | | | | | | | | |
|---|---|---|---|---|---|---|---|---|---|---|---|
| CC | GO:0032991 | 29 | 8.02E-04 | 58 | 363 | 1253 | 1.73 | 0.12 | 0.04 | 0.97 | macromolecular complex |
| CC | GO:0022624 | 4 | 2.75E-03 | 58 | 7 | 1253 | 12.34 | 0.36 | 0.09 | 3.27 | proteasome accessory complex |
| CC | GO:0005730 | 6 | 4.31E-03 | 58 | 25 | 1253 | 5.18 | 0.50 | 0.11 | 5.09 | nucleolus |
| CC | GO:0005576 | 7 | 6.04E-03 | 58 | 38 | 1253 | 3.98 | 0.62 | 0.13 | 7.06 | extracellular region |
| CC | GO:0005732 | 3 | 1.86E-02 | 58 | 5 | 1253 | 12.96 | 0.95 | 0.29 | 20.34 | small nucleolar ribonucleoprotein complex |
| CC | GO:0005758 | 3 | 2.71E-02 | 58 | 6 | 1253 | 10.80 | 0.99 | 0.36 | 28.29 | mitochondrial intermembrane space |
| CC | GO:0031970 | 3 | 2.71E-02 | 58 | 6 | 1253 | 10.80 | 0.99 | 0.36 | 28.29 | organelle envelope lumen |
| CC | GO:0030684 | 4 | 2.75E-02 | 58 | 15 | 1253 | 5.76 | 0.99 | 0.34 | 28.65 | preribosome |
| CC | GO:0043228 | 15 | 2.91E-02 | 58 | 183 | 1253 | 1.77 | 0.99 | 0.33 | 30.01 | non-membrane-bounded organelle |
| CC | GO:0043232 | 15 | 2.91E-02 | 58 | 183 | 1253 | 1.77 | 0.99 | 0.33 | 30.01 | intracellular non-membrane-bounded organelle |
| CC | GO:0044424 | 44 | 5.45E-02 | 58 | 812 | 1253 | 1.17 | 1.00 | 0.50 | 49.23 | intracellular part |
| CC | GO:0031974 | 10 | 6.36E-02 | 58 | 113 | 1253 | 1.91 | 1.00 | 0.53 | 54.83 | membrane-enclosed lumen |
| CC | GO:0043234 | 18 | 7.16E-02 | 58 | 264 | 1253 | 1.47 | 1.00 | 0.55 | 59.29 | protein complex |
| CC | GO:0031429 | 2 | 8.89E-02 | 58 | 2 | 1253 | 21.60 | 1.00 | 0.61 | 67.61 | box H/ACA snoRNP complex |
| CC | GO:0042555 | 2 | 8.89E-02 | 58 | 2 | 1253 | 21.60 | 1.00 | 0.61 | 67.61 | MCM complex |
| CC | GO:0005697 | 2 | 8.89E-02 | 58 | 2 | 1253 | 21.60 | 1.00 | 0.61 | 67.61 | telomerase holoenzyme complex |
| MF | GO:0003723 | 9 | 7.28E-03 | 45 | 86 | 1277 | 2.97 | 0.65 | 0.65 | 8.30 | RNA binding |
| MF | GO:0003676 | 15 | 6.90E-02 | 45 | 274 | 1277 | 1.55 | 1.00 | 0.99 | 57.15 | nucleic acid binding |
| MF | GO:0097159 | 24 | 7.56E-02 | 45 | 519 | 1277 | 1.31 | 1.00 | 0.98 | 60.61 | organic cyclic compound binding |
| MF | GO:0061135 | 2 | 9.99E-02 | 45 | 3 | 1277 | 18.92 | 1.00 | 0.98 | 71.30 | endopeptidase regulator activity |
| MF | GO:0061134 | 2 | 9.99E-02 | 45 | 3 | 1277 | 18.92 | 1.00 | 0.98 | 71.30 | peptidase regulator activity |

Table S14. Pathway enrichment analysis from day 1 downregulated genes using KEGG database

| KEGG ID | Pathway | N | Count | Pvalue | GeneRatio |
|---|---|---|---|---|---|
| path:spu00010 | Glycolysis / Gluconeogenesis | 53 | 5 | 4.61E-03 | 9.43E-02 |
| path:spu00040 | Pentose and glucuronate interconversions | 85 | 5 | 3.13E-02 | 5.88E-02 |
| path:spu00053 | Ascorbate and aldarate metabolism | 85 | 5 | 3.13E-02 | 5.88E-02 |
| path:spu00071 | Fatty acid degradation | 57 | 4 | 3.01E-02 | 7.02E-02 |
| path:spu00250 | Alanine, aspartate and glutamate metabolism | 36 | 3 | 3.81E-02 | 8.33E-02 |
| path:spu00270 | Cysteine and methionine metabolism | 70 | 5 | 1.47E-02 | 7.14E-02 |
| path:spu00280 | Valine, leucine and isoleucine degradation | 61 | 6 | 1.55E-03 | 9.84E-02 |
| path:spu00340 | Histidine metabolism | 37 | 4 | 6.91E-03 | 1.08E-01 |
| path:spu00350 | Tyrosine metabolism | 28 | 3 | 1.97E-02 | 1.07E-01 |
| path:spu00380 | Tryptophan metabolism | 74 | 6 | 4.15E-03 | 8.11E-02 |
| path:spu00410 | beta-Alanine metabolism | 30 | 3 | 2.37E-02 | 1.00E-01 |
| path:spu00620 | Pyruvate metabolism | 43 | 5 | 1.82E-03 | 1.16E-01 |
| path:spu00730 | Thiamine metabolism | 17 | 2 | 4.77E-02 | 1.18E-01 |
| path:spu00760 | Nicotinate and nicotinamide metabolism | 40 | 3 | 4.97E-02 | 7.50E-02 |
| path:spu00790 | Folate biosynthesis | 48 | 6 | 4.25E-04 | 1.25E-01 |
| path:spu00920 | Sulfur metabolism | 20 | 4 | 6.64E-04 | 2.00E-01 |
| path:spu01100 | Metabolic pathways | 1996 | 66 | 7.75E-07 | 3.31E-02 |
| path:spu01200 | Carbon metabolism | 138 | 7 | 2.42E-02 | 5.07E-02 |
| path:spu04146 | Peroxisome | 146 | 9 | 3.14E-03 | 6.16E-02 |
| path:spu04512 | ECM-receptor interaction | 50 | 4 | 1.96E-02 | 8.00E-02 |

Table S15. Pathway enrichment analysis from day 1 Upregulated genes using KEGG database

| KEGG ID | Pathway | N | Count | Pvalue | GeneRatio |
|---|---|---|---|---|---|
| path:spu00970 | Aminoacyl-tRNA biosynthesis | 74 | 10 | 1.96E-03 | 1.35E-01 |
| path:spu03008 | Ribosome biogenesis in eukaryotes | 91 | 26 | 1.41E-14 | 2.86E-01 |
| path:spu03010 | Ribosome | 140 | 30 | 5.25E-13 | 2.14E-01 |
| path:spu03013 | RNA transport | 170 | 16 | 5.02E-03 | 9.41E-02 |
| path:spu03015 | mRNA surveillance pathway | 72 | 7 | 4.76E-02 | 9.72E-02 |
| path:spu03020 | RNA polymerase | 31 | 7 | 4.21E-04 | 2.26E-01 |
| path:spu03040 | Spliceosome | 177 | 14 | 3.38E-02 | 7.91E-02 |
| path:spu03050 | Proteasome | 48 | 15 | 1.70E-09 | 3.13E-01 |
| path:spu04141 | Protein processing in endoplasmic reticulum | 166 | 13 | 4.28E-02 | 7.83E-02 |

Table S16. Pathway enrichment analysis from day 3 downgulated genes using KEGG database

| KEGG ID | Pathway | N | Count | Pvalue | GeneRatio |
|---|---|---|---|---|---|
| path:spu00340 | Histidine metabolism | 37 | 3 | 2.58E-02 | 8.11E-02 |
| path:spu00380 | Tryptophan metabolism | 74 | 4 | 3.90E-02 | 5.41E-02 |
| path:spu00500 | Starch and sucrose metabolism | 39 | 3 | 2.96E-02 | 7.69E-02 |
| path:spu00620 | Pyruvate metabolism | 43 | 3 | 3.80E-02 | 6.98E-02 |
| path:spu00670 | One carbon pool by folate | 17 | 2 | 3.43E-02 | 1.18E-01 |
| path:spu00730 | Thiamine metabolism | 17 | 2 | 3.43E-02 | 1.18E-01 |
| path:spu01100 | Metabolic pathways | 1996 | 43 | 4.28E-02 | 2.15E-02 |
| path:spu04145 | Phagosome | 165 | 11 | 1.14E-04 | 6.67E-02 |
| path:spu04146 | Peroxisome | 146 | 7 | 1.30E-02 | 4.79E-02 |

Table S17. Pathway enrichment analysis from day 3 upgulated genes using KEGG database

| KEGG ID | Pathway | N | Count | Pvalue | GeneRatio |
|---|---|---|---|---|---|
| path:spu03008 | Ribosome biogenesis in eukaryotes | 91 | 7 | 4.94E-03 | 7.69E-02 |
| path:spu03010 | Ribosome | 140 | 24 | 3.89E-15 | 1.71E-01 |
| path:spu03050 | Proteasome | 48 | 9 | 1.14E-06 | 1.88E-01 |
| path:spu04141 | Protein processing in endoplasmic reticulum | 166 | 13 | 1.06E-04 | 7.83E-02 |
| path:spu04145 | Phagosome | 165 | 8 | 3.73E-02 | 4.85E-02 |

Table S18. Clusters of coexpressed transcripts

| ID | Description | Cluster | Adjusted P-value | GeneID |
|---|---|---|---|---|
| Transcript_442613 | catalase | 1 | 5.41E-42 | 548621 |
| Transcript_268943 | cytochrome P450 2U1 | 2 | 4.53E-38 | 581885 |
| Transcript_319952 | SOX4 | 2 | 9.30E-34 | 593520 |
| Transcript_285077 | zinc transporter ZIP14 | 2 | 5.19E-31 | 763759 |
| Transcript_340144 | uncharacterized protein K02A2.6-like\|Reverse transcriptase domain-containing protein | 2 | 9.49E-30 | 105446129 |
| Transcript_328691 | MAM and LDL-receptor class A domain-containing protein 1 | 1 | 7.43E-27 | 574589 |
| Transcript_298293 | collagen alpha-1(XII) chain\|tenascin-X | 1 | 2.46E-26 | 763365 |
| Transcript_197637 | retinol dehydrogenase 12\|dehydrogenase/reductase SDR family member 13-like | 3 | 7.29E-26 | 589120 |
| Transcript_204404 | laminin subunit alpha\|laminin alpha chain | 1 | 2.41E-25 | 578626 |
| Transcript_298730 | laminin subunit beta-1\|laminin, beta 1 | 4 | 2.59E-24 | 582206 |
| Transcript_202375 | fibrillin-2\|uncharacterized protein LOC105436405 | 3 | 4.08E-24 | 105436405 |
| Transcript_287164 | nucleolar complex protein 2 homolog | 3 | 8.11E-24 | 585237 |
| Transcript_243955 | glycerol kinase | 3 | 2.87E-23 | 580504 |
| Transcript_445893 | ATPase family AAA domain-containing protein 3-B | 3 | 5.85E-23 | 588531 |
| Transcript_298217 | 4-aminobutyrate aminotransferase, mitochondrial | 4 | 4.79E-22 | 577656 |
| Transcript_264429 | sushi domain-containing protein 2\|uncharacterized protein K03H1.5 | 1 | 4.74E-21 | 583122 |
| Transcript_172789 | neuroblast differentiation-associated protein AHNAK-like | 1 | 1.09E-20 | 105441784 |
| Transcript_442463 | probable dimethyladenosine transferase | 2 | 1.72E-20 | 574553 |
| Transcript_280127 | RNA granule protein invertebrate\|TPA: RNA granule protein invertebrate | 2 | 3.94E-20 | 580088 |
| Transcript_406209 | sulfite oxidase | 4 | 7.13E-20 | 586173 |
| Transcript_238520 | lysyl oxidase-like 2 | 1 | 5.75E-19 | 585092 |
| Transcript_267297 | pumilio homolog 3 | 3 | 7.79E-19 | 583455 |
| Transcript_443315 | prestalk protein | 1 | 1.77E-18 | 590178 |
| Transcript_461742 | stromelysin-3-like\|matrix metalloproteinase-24-like | 1 | 2.08E-18 | 105436437 |
| Transcript_385039 | aminopeptidase N | 1 | 2.21E-18 | 578352 |
| Transcript_438891 | HLX | 5 | 1.38E-17 | 100892685 |
| Transcript_350383 | telomerase reverse transcriptase-long\|telomerase reverse transcriptase- | 3 | 2.59E-17 | 105447117 |
| Transcript_344692 | 26S proteasome non-ATPase regulatory subunit 12 | 3 | 2.62E-17 | 593806 |
| Transcript_429867 | HMX | 3 | 8.86E-17 | 579532 |
| Transcript_395263 | 26S proteasome non-ATPase regulatory subunit 5 | 2 | 1.55E-16 | 588345 |
| Transcript_198009 | uncharacterized protein LOC582064\|inter-alpha-trypsin inhibitor heavy chain H2 | 4 | 2.22E-16 | 582064 |
| Transcript_246166 | diphthine synthase | 3 | 2.57E-16 | 577892 |

| | | | | |
|---|---|---|---|---|
| Transcript_430669 | LOW QUALITY PROTEIN: uncharacterized protein LOC580672\|apolipoprotein B-100 | 4 | 2.78E-16 | 580672 |
| Transcript_328858 | uncharacterized protein LOC100893633\|Arylamine N-acetyltransferase | 4 | 2.78E-16 | 100893633 |
| Transcript_414587 | solute carrier family 35 member F2 | 2 | 6.89E-16 | 574685 |
| Transcript_364016 | prohibitin | 2 | 7.21E-16 | 584483 |
| Transcript_431156 | GTP-binding protein 8 | 1 | 8.05E-16 | 590744 |
| Transcript_366520 | GDP-mannose 4,6 dehydratase | 2 | 1.21E-15 | 588885 |
| Transcript_288662 | microfibril-associated glycoprotein 4\|tenascin-N | 1 | 1.84E-15 | 753235 |
| Transcript_351600 | peroxisomal multifunctional enzyme type 2\|hydroxysteroid (17-beta) dehydrogenase 4 | 1 | 2.44E-15 | 581580 |
| Transcript_267740 | ribosome biogenesis protein WDR12 | 2 | 3.19E-15 | 591474 |
| Transcript_404053 | mRNA turnover protein 4 homolog | 3 | 3.39E-15 | 578804 |
| Transcript_364345 | proteasome activator complex subunit 3 | 3 | 4.11E-15 | 585921 |
| Transcript_461114 | yrdC domain-containing protein, mitochondrial | 2 | 4.36E-15 | 588740 |
| Transcript_204651 | OTU domain-containing protein 4\|putative bifunctional UDP-N-acetylglucosamine transferase a | 3 | 6.67E-15 | 593098 |
| Transcript_425984 | translocator protein | 1 | 7.60E-15 | 583617 |
| Transcript_179579 | methenyltetrahydrofolate synthase domain-containing protein | 3 | 8.49E-15 | 588795 |
| Transcript_183015 | LOW QUALITY PROTEIN: histidine--tRNA ligase, cytoplasmic | 3 | 1.13E-14 | 105444109 |
| Transcript_206897 | actin, muscle | 1 | 1.22E-14 | 581500 |
| Transcript_421726 | ribosome production factor 2 homolog | 2 | 1.23E-14 | 755131 |
| Transcript_365697 | uncharacterized protein LOC589195 | 3 | 1.28E-14 | 589195 |
| Transcript_428650 | ammonium transporter Rh type B\|ammonium transporter Rh type B-A\|ammonium transporter | 1 | 1.34E-14 | 584796 |
| Transcript_431326 | 26S proteasome non-ATPase regulatory subunit 3 | 3 | 1.41E-14 | 578762 |
| Transcript_384474 | transmembrane protein KIAA1109 | 4 | 1.46E-14 | 586038 |
| Transcript_365782 | LOW QUALITY PROTEIN: eukaryotic translation initiation factor 3 subunit I | 2 | 2.21E-14 | 589571 |
| Transcript_279405 | ATP-dependent RNA helicase HAS1\|ATP-dependent RNA helicase DDX18\|DEAD (Asp-Glu-Ala-As | 3 | 2.62E-14 | 577986 |
| Transcript_197448 | negative elongation factor C/D\|negative elongation factor D | 2 | 3.61E-14 | 581078 |
| Transcript_375637 | protein Wnt-6 | 3 | 6.03E-14 | 585147 |
| Transcript_344765 | guanine nucleotide-binding protein-like NSN1\|guanine nucleotide-binding protein-like 3 homol | 2 | 7.57E-14 | 577855 |
| Transcript_354584 | enolase\|alpha-enolase-like | 1 | 7.95E-14 | 579256 |
| Transcript_423794 | activator of 90 kDa heat shock protein ATPase homolog 1 | 2 | 1.22E-13 | 575643 |
| Transcript_359489 | eukaryotic translation initiation factor 3 subunit K | 3 | 1.62E-13 | 587575 |
| Transcript_397594 | LOW QUALITY PROTEIN: RNA cytidine acetyltransferase\|N-acetyltransferase 10 (GCN5-related) | 2 | 1.63E-13 | 576671 |
| Transcript_373223 | RNA polymerase II subunit A C-terminal domain phosphatase SSU72 | 2 | 1.68E-13 | 574857 |
| Transcript_460570 | alpha-catulin | 1 | 1.81E-13 | 576107 |
| Transcript_417350 | ribosome biogenesis protein NOP53\|glioma tumor suppressor candidate region gene 2 protein | 2 | 1.98E-13 | 593605 |
| Transcript_333973 | tubulin polymerization-promoting protein family member 2 | 1 | 2.23E-13 | 577151 |
| Transcript_183358 | alpha-actinin\|alpha-actinin, sarcomeric | 1 | 2.26E-13 | 592971 |

| | | | | |
|---|---|---|---|---|
| Transcript_270919 | heterogeneous nuclear ribonucleoprotein C | 3 | 2.92E-13 | 100893477 |
| Transcript_263624 | protein Wnt-9a | 3 | 3.13E-13 | 575320 |
| Transcript_298954 | uncharacterized protein LOC577534 | 2 | 3.22E-13 | 577534 |
| Transcript_202461 | opsin Rh5 | 2 | 4.05E-13 | 579899 |
| Transcript_427877 | uncharacterized protein LOC593232\|Ankyrin repeat-containing domain-containing protein | 3 | 4.30E-13 | 593232 |
| Transcript_375646 | hydroxyacylglutathione hydrolase, mitochondrial | 1 | 5.05E-13 | 578596 |
| Transcript_373821 | mitochondrial intermembrane space import and assembly protein 40 | 2 | 5.98E-13 | 582540 |
| Transcript_256657 | carbohydrate sulfotransferase 1 | 2 | 7.10E-13 | 579726 |
| Transcript_412104 | probable cation-transporting ATPase 13A3\|ATPase type 13A3\|putative cation-transporting ATP | 1 | 7.52E-13 | 577790 |
| Transcript_314464 | CMP-N-acetylneuraminate-beta-1,4-galactoside alpha-2,3-sialyltransferase | 1 | 8.96E-13 | 754196 |
| Transcript_359719 | heat shock cognate 71 kDa protein\|heat shock 70kDa protein 8 | 2 | 9.55E-13 | 576276 |
| Transcript_395874 | U3 small nucleolar RNA-associated protein 18 homolog | 3 | 9.76E-13 | 579700 |
| Transcript_405927 | NADPH oxidase 4 | 2 | 1.11E-12 | 100889322 |
| Transcript_448868 | IgGFc-binding protein\|Fc fragment of IgG binding protein\|zonadhesin | 1 | 1.14E-12 | 762415 |
| Transcript_271439 | T-complex protein 1 subunit zeta | 2 | 1.41E-12 | 577393 |
| Transcript_330216 | aspartyl/asparaginyl beta-hydroxylase\|bromodomain-containing protein 4 | 1 | 1.92E-12 | 593797 |
| Transcript_200469 | aquaporin-8 | 1 | 2.12E-12 | 589852 |
| Transcript_197881 | eukaryotic translation initiation factor 3 subunit E | 2 | 2.28E-12 | 754815 |
| Transcript_182900 | sodium-coupled monocarboxylate transporter 1 | 1 | 2.33E-12 | 580845 |
| Transcript_396851 | soluble adenylyl cyclase | 3 | 2.73E-12 | 574069 |
| Transcript_383783 | bone morphogenetic protein 1 homolog | 1 | 2.81E-12 | 373360 |
| Transcript_431169 | T-complex protein 1 subunit theta | 3 | 3.11E-12 | 583086 |
| Transcript_443125 | nucleolar RNA helicase 2 | 2 | 3.20E-12 | 581409 |
| Transcript_277158 | ATP-binding cassette sub-family F member 1 | 2 | 3.40E-12 | 576400 |
| Transcript_249697 | lupus La protein homolog | 2 | 3.41E-12 | 105443896 |
| Transcript_442985 | asparagine synthetase [glutamine-hydrolyzing] | 2 | 3.42E-12 | 575946 |
| Transcript_262723 | probable serine/threonine-protein kinase DDB_G0278665 | 1 | 4.05E-12 | 577336 |
| Transcript_225361 | contactin-associated protein-like 2\|contactin-associated protein-like 5 | 1 | 4.57E-12 | 575215 |
| Transcript_433953 | PWP1 | 2 | 4.80E-12 | 583947 |
| Transcript_353989 | nidogen-1 | 1 | 5.18E-12 | 586685 |
| Transcript_350978 | ribosome biogenesis regulatory protein homolog | 3 | 5.94E-12 | 575632 |
| Transcript_294196 | low-density lipoprotein receptor-related protein 1 | 1 | 8.65E-12 | 582052 |
| Transcript_278301 | tubulin alpha-1A chain | 2 | 9.88E-12 | 754102 |
| Transcript_258721 | glutamyl aminopeptidase | 1 | 1.03E-11 | 587683 |
| Transcript_440792 | medium-chain acyl-CoA ligase ACSF2, mitochondrial\|acyl-CoA synthetase family member 2, mit | 1 | 1.35E-11 | 592091 |
| Transcript_394161 | pyrroline-5-carboxylate reductase 1, mitochondrial | 2 | 1.39E-11 | 583853 |

| | | | | |
|---|---|---|---|---|
| Transcript_267275 | prohibitin-2 | 3 | 1.70E-11 | 575755 |
| Transcript_263596 | BTF3L4 | 2 | 2.16E-11 | 586588 |
| Transcript_267873 | H/ACA ribonucleoprotein complex subunit 1 | 3 | 2.51E-11 | 592267 |
| Transcript_385986 | midasin | 3 | 2.92E-11 | 575173 |
| Transcript_336637 | eukaryotic peptide chain release factor GTP-binding subunit ERF3A\|G1 to S phase transition 2 | 3 | 2.92E-11 | 580307 |
| Transcript_175325 | TAP26 | 2 | 2.98E-11 | 579591 |
| Transcript_310211 | protein arginine N-methyltransferase 1 | 3 | 3.24E-11 | 590987 |
| Transcript_230511 | ribosomal RNA processing protein 1 homolog A\|ribosomal RNA processing protein 1 homolog B | 2 | 3.35E-11 | 594061 |
| Transcript_68378 | uncharacterized protein LOC584236 | 2 | 3.50E-11 | 584236 |
| Transcript_452859 | 25S rRNA (cytosine-C(5))-methyltransferase nop2 | 3 | 3.62E-11 | 582075 |
| Transcript_352347 | RNA-binding protein with serine-rich domain 1 | 3 | 3.88E-11 | 585553 |
| Transcript_354900 | mitochondrial import receptor subunit TOM40 homolog | 3 | 3.95E-11 | 585934 |
| Transcript_365226 | protein phosphatase 1G | 2 | 4.53E-11 | 594091 |
| Transcript_296057 | carbohydrate sulfotransferase 15-like | 1 | 4.53E-11 | 100892795 |
| Transcript_222572 | bone morphogenetic protein BMP2/4 | 3 | 4.69E-11 | 373196 |
| Transcript_216724 | uncharacterized protein LOC587705\|putative sterigmatocystin biosynthesis dehydrogenase stc | 1 | 4.87E-11 | 587705 |
| Transcript_311413 | amassin\|coelomocyte amassing protein | 3 | 4.95E-11 | 373503 |
| Transcript_263997 | mitochondrial import inner membrane translocase subunit Tim17-B | 3 | 5.59E-11 | 577928 |
| Transcript_242486 | polycomb protein eed-A | 3 | 5.77E-11 | 581239 |
| Transcript_454730 | LOW QUALITY PROTEIN: allene oxide synthase-lipoxygenase protein | 1 | 6.46E-11 | 584481 |
| Transcript_279511 | mucosa-associated lymphoid tissue lymphoma translocation protein 1\|mucosa-associated lymp | 2 | 6.81E-11 | 590194 |
| Transcript_322705 | THO complex subunit 6 homolog | 3 | 6.98E-11 | 580431 |
| Transcript_421292 | sphingosine-1-phosphate lyase 1 | 3 | 8.18E-11 | 585643 |
| Transcript_226274 | uncharacterized protein LOC583933 | 1 | 8.46E-11 | 583933 |
| Transcript_435175 | sodium- and chloride-dependent neutral and basic amino acid transporter B(0+) | 1 | 9.08E-11 | 594544 |
| Transcript_224605 | ribosomal RNA small subunit methyltransferase NEP1 | 2 | 9.38E-11 | 589275 |
| Transcript_342811 | epoxide hydrolase 1 | 1 | 9.89E-11 | 588166 |
| Transcript_328958 | H/ACA ribonucleoprotein complex subunit DKC1\|dyskeratosis congenita 1, dyskerin | 2 | 1.10E-10 | 584530 |
| Transcript_421302 | equilibrative nucleoside transporter 1\|microtubule-associated protein 10 | 4 | 1.13E-10 | 592207 |
| Transcript_339471 | fibrillin-1 | 1 | 1.23E-10 | 575027 |
| Transcript_431258 | methanethiol oxidase\|selenium-binding protein 1 | 1 | 1.23E-10 | 582523 |
| Transcript_7357 | cathepsin Z | 2 | 1.37E-10 | 580617 |
| Transcript_432532 | LMO2 | 2 | 1.48E-10 | 100892046 |
| Transcript_310947 | rRNA-processing protein FCF1 homolog | 3 | 1.49E-10 | 586103 |
| Transcript_407052 | C3 and PZP-like alpha-2-macroglobulin domain-containing protein 8\|C3 and PZP-like, alpha-2-m | 1 | 1.50E-10 | 579831 |
| Transcript_461041 | eEF1A lysine and N-terminal methyltransferase\|methyltransferase-like protein 13 | 2 | 1.67E-10 | 754600 |

| | | | | |
|---|---|---|---|---|
| Transcript_205415 | H/ACA ribonucleoprotein complex subunit 2-like protein | 3 | 1.73E-10 | 577044 |
| Transcript_254399 | vacuolar protein sorting-associated protein 13A\|vacuolar protein sorting-associated protein 13[illegible] | 1 | 1.75E-10 | 575234 |
| Transcript_439620 | uridine-cytidine kinase 2\|uridine-cytidine kinase 2-B | 2 | 1.86E-10 | 588945 |
| Transcript_208693 | neutral and basic amino acid transport protein rBAT | 1 | 1.88E-10 | 586832 |
| Transcript_395337 | formin-J | 3 | 2.03E-10 | 588660 |
| Transcript_184579 | RWD domain-containing protein 1 | 2 | 2.11E-10 | 590636 |
| Transcript_282088 | 39S ribosomal protein L17, mitochondrial | 2 | 2.40E-10 | 591498 |
| Transcript_271370 | rRNA 2'-O-methyltransferase fibrillarin | 3 | 2.46E-10 | 752335 |
| Transcript_417755 | transmembrane protein 69 | 2 | 2.59E-10 | 586174 |
| Transcript_409638 | heat shock 70 kDa protein 14 | 3 | 2.73E-10 | 588149 |
| Transcript_259246 | pre-mRNA-splicing factor 38B | 2 | 3.02E-10 | 100889783 |
| Transcript_308724 | uncharacterized protein LOC577145 | 1 | 3.19E-10 | 577145 |
| Transcript_330347 | stress-70 protein, mitochondrial\|heat shock 70kDa protein 9 (mortalin) | 3 | 3.20E-10 | 577721 |
| Transcript_215804 | leucine-rich repeat-containing protein 74B | 1 | 3.42E-10 | 584206 |
| Transcript_204158 | cystine/glutamate transporter | 4 | 3.52E-10 | 579243 |
| Transcript_243187 | DNA-directed RNA polymerase I subunit RPA49 | 2 | 3.56E-10 | 588897 |
| Transcript_290985 | glycine--tRNA ligase | 2 | 4.15E-10 | 578936 |
| Transcript_403536 | ribosome production factor 1 | 2 | 4.37E-10 | 592162 |
| Transcript_295742 | 10 kDa heat shock protein, mitochondrial | 3 | 4.72E-10 | 762428 |
| Transcript_197679 | uncharacterized protein LOC105446114 | 4 | 4.73E-10 | 105446114 |
| Transcript_456681 | CD82 antigen\|carbohydrate sulfotransferase 1-like | 1 | 4.92E-10 | 100889350 |
| Transcript_263817 | eukaryotic translation initiation factor 2 subunit 1 | 3 | 4.98E-10 | 586283 |
| Transcript_357723 | translation elongation factor 1B beta subunit | 3 | 6.19E-10 | 574071 |
| Transcript_234345 | bromodomain-containing protein 4-like\|dentin sialophosphoprotein | 3 | 6.36E-10 | 105441791 |
| Transcript_270485 | U3 small nucleolar RNA-interacting protein 2 | 2 | 6.67E-10 | 582919 |
| Transcript_202897 | dimethyladenosine transferase 2, mitochondrial | 2 | 6.97E-10 | 589505 |
| Transcript_427783 | uncharacterized protein LOC575162 | 1 | 7.09E-10 | 575162 |
| Transcript_319739 | carboxypeptidase B | 1 | 7.54E-10 | 764971 |
| Transcript_183115 | nucleolar GTP-binding protein 2 | 2 | 8.38E-10 | 585370 |
| Transcript_367031 | ribosome biogenesis protein bop1-B | 2 | 8.81E-10 | 575063 |
| Transcript_272225 | protein ABHD11\|alpha/beta hydrolase domain-containing protein 11 | 3 | 8.95E-10 | 575326 |
| Transcript_288791 | ATP-dependent RNA helicase DHX30 | 2 | 1.04E-09 | 577803 |
| Transcript_244563 | mitochondrial import inner membrane translocase subunit TIM44 | 3 | 1.08E-09 | 752190 |
| Transcript_438545 | all-trans-retinol 13,14-reductase\|putative all-trans-retinol 13,14-reductase | 4 | 1.09E-09 | 581285 |
| Transcript_172871 | protein RER1 | 2 | 1.09E-09 | 579195 |
| Transcript_417215 | alanine--glyoxylate aminotransferase 2, mitochondrial | 4 | 1.13E-09 | 594821 |

| | | | | |
|---|---|---|---|---|
| Transcript_256986 | mitochondrial import inner membrane translocase subunit Tim10-B | 3 | 1.16E-09 | 589835 |
| Transcript_266569 | stress-induced-phosphoprotein 1 | 3 | 1.17E-09 | 765087 |
| Transcript_447903 | uracil-DNA glycosylase | 4 | 1.29E-09 | 586702 |
| Transcript_342844 | peroxiredoxin | 2 | 1.31E-09 | 581408 |
| Transcript_411315 | propionyl-CoA carboxylase alpha chain, mitochondrial | 2 | 1.40E-09 | 582365 |
| Transcript_265130 | RRP15-like protein | 2 | 1.46E-09 | 577322 |
| Transcript_358391 | ribosome biogenesis protein NSA2 homolog | 3 | 1.55E-09 | 577954 |
| Transcript_264331 | fibrillin-2\|fibrillin-1 | 1 | 1.72E-09 | 582487 |
| Transcript_455249 | maleylacetoacetate isomerase | 5 | 1.89E-09 | 580863 |
| Transcript_443343 | lysine--tRNA ligase | 3 | 2.22E-09 | 588700 |
| Transcript_393409 | serine-enriched protein | 2 | 2.28E-09 | 100888499 |
| Transcript_264379 | 60 kDa heat shock protein, mitochondrial | 2 | 2.33E-09 | 590510 |
| Transcript_236885 | serine/threonine-protein kinase rio2 | 2 | 2.44E-09 | 579218 |
| Transcript_285991 | hypoxia up-regulated protein 1\|hypoxia up-regulated 1 | 3 | 2.63E-09 | 592339 |
| Transcript_242646 | 60S ribosomal protein L8 | 2 | 2.67E-09 | 591341 |
| Transcript_204616 | T-complex protein 1 subunit delta | 2 | 2.96E-09 | 579722 |
| Transcript_194648 | coiled-coil domain-containing protein 12 | 3 | 3.14E-09 | 585976 |
| Transcript_410118 | ATP-dependent RNA helicase DDX24 | 2 | 3.23E-09 | 578876 |
| Transcript_232798 | protein arginine N-methyltransferase 3 | 3 | 3.24E-09 | 588858 |
| Transcript_243757 | small nuclear ribonucleoprotein Sm D3 | 3 | 3.58E-09 | 105438529 |
| Transcript_431275 | eukaryotic translation initiation factor 2 subunit 2 | 3 | 3.61E-09 | 590481 |
| Transcript_237829 | gamma-butyrobetaine dioxygenase | 1 | 3.73E-09 | 577198 |
| Transcript_353055 | deoxyhypusine hydroxylase | 2 | 3.81E-09 | 582221 |
| Transcript_351300 | valine--tRNA ligase | 3 | 3.92E-09 | 590476 |
| Transcript_253045 | cytochrome P450 10 | 2 | 4.01E-09 | 576441 |
| Transcript_411364 | CCR4-NOT transcription complex subunit 11 | 2 | 4.27E-09 | 594372 |
| Transcript_215364 | 26S proteasome non-ATPase regulatory subunit 7 | 3 | 4.44E-09 | 575334 |
| Transcript_78944 | DNA-directed RNA polymerase I subunit RPA12 | 2 | 4.56E-09 | 100892475 |
| Transcript_331128 | deoxynucleotidyltransferase terminal-interacting protein 2 | 2 | 4.71E-09 | 586304 |
| Transcript_285870 | 26S proteasome regulatory subunit 6A-B\|26S protease regulatory subunit 6A-B\|26S protease re | 2 | 4.90E-09 | 576830 |
| Transcript_324100 | hydroxyacid oxidase 1 | 1 | 4.95E-09 | 584105 |
| Transcript_280947 | 26S proteasome non-ATPase regulatory subunit 4 | 2 | 5.06E-09 | 575737 |
| Transcript_279609 | 26S proteasome non-ATPase regulatory subunit 2 | 2 | 5.09E-09 | 588529 |
| Transcript_367924 | T-complex protein 1 subunit alpha | 3 | 5.28E-09 | 582138 |
| Transcript_359457 | alpha-2-macroglobulin | 3 | 5.54E-09 | 594798 |
| Transcript_272476 | leucine-rich repeat-containing protein 46 | 2 | 6.23E-09 | 100889057 |

| | | | | |
|---|---|---|---|---|
| Transcript_428533 | proteasomal ubiquitin receptor ADRM1 | 2 | 6.25E-09 | 753175 |
| Transcript_461744 | serine--tRNA ligase, cytoplasmic | 2 | 7.26E-09 | 588782 |
| Transcript_437045 | betaine--homocysteine S-methyltransferase 1 | 1 | 7.59E-09 | 752628 |
| Transcript_218326 | uncharacterized protein LOC100893287 | 1 | 7.60E-09 | 100893287 |
| Transcript_420760 | alkaline phosphatase | 1 | 8.51E-09 | 580300 |
| Transcript_454673 | protein SDA1 homolog\|SDA1 domain containing 1 | 3 | 8.75E-09 | 585590 |
| Transcript_441945 | queuosine salvage protein\|UPF0553 protein C9orf64 | 2 | 8.75E-09 | 584962 |
| Transcript_256633 | bifunctional purine biosynthesis protein PURH | 2 | 9.25E-09 | 588680 |
| Transcript_386779 | FRAS1-related extracellular matrix protein 1\|FRAS1 related extracellular matrix 1 | 1 | 9.46E-09 | 591943 |
| Transcript_297492 | laminin subunit alpha-2\|laminin subunit alpha-1 | 5 | 9.73E-09 | 587274 |
| Transcript_397023 | cadherin EGF LAG seven-pass G-type receptor 1 | 1 | 9.95E-09 | 105440441 |
| Transcript_190981 | superoxide dismutase [Cu-Zn] | 1 | 1.02E-08 | 579361 |
| Transcript_41462 | EF-hand calcium-binding domain-containing protein 6 | 4 | 1.06E-08 | 589273 |
| Transcript_271917 | dihydroflavonol 4-reductase\|putative NADPH-dependent methylglyoxal reductase GRP2\|putati | 3 | 1.09E-08 | 585152 |
| Transcript_268427 | steroid 17-alpha-hydroxylase/17,20 lyase | 4 | 1.19E-08 | 585029 |
| Transcript_384438 | thyrotropin-releasing hormone-degrading ectoenzyme | 1 | 1.20E-08 | 588093 |
| Transcript_298313 | spectrin alpha chain, non-erythrocytic 1 | 1 | 1.32E-08 | 580822 |
| Transcript_437221 | apoptosis-inducing factor 1, mitochondrial\|apoptosis-inducing factor, mitochondrion-associated | 3 | 1.37E-08 | 578253 |
| Transcript_456290 | claspin\|claspin homolog | 1 | 1.37E-08 | 575148 |
| Transcript_343920 | stAR-related lipid transfer protein 5 | 2 | 1.37E-08 | 584940 |
| Transcript_293280 | probable ribosome biogenesis protein RLP24 | 2 | 1.39E-08 | 581865 |
| Transcript_245310 | stAR-related lipid transfer protein 7, mitochondrial | 2 | 1.42E-08 | 578525 |
| Transcript_427274 | contactin-2 | 1 | 1.44E-08 | 100893855 |
| Transcript_431447 | BRCA2 and CDKN1A-interacting protein | 2 | 1.51E-08 | 586387 |
| Transcript_461384 | ATP-binding cassette sub-family C member 9 | 4 | 1.53E-08 | 594397 |
| Transcript_404538 | inactive C-alpha-formylglycine-generating enzyme 2 | 3 | 1.59E-08 | 576301 |
| Transcript_185289 | uncharacterized protein LOC100887836 | 1 | 1.75E-08 | 100887836 |
| Transcript_381525 | uronyl 2-sulfotransferase | 1 | 1.76E-08 | 593546 |
| Transcript_185057 | GDH/6PGL endoplasmic bifunctional protein\|hexose-6-phosphate dehydrogenase (glucose 1-de | 1 | 2.02E-08 | 590387 |
| Transcript_360124 | cytochrome c | 3 | 2.03E-08 | 575347 |
| Transcript_295374 | KH domain-containing, RNA-binding, signal transduction-associated protein 2 | 2 | 2.09E-08 | 588527 |
| Transcript_386824 | WD repeat-containing protein 74 | 2 | 2.16E-08 | 587044 |
| Transcript_302605 | proliferation-associated protein 2G4 | 2 | 2.23E-08 | 576641 |
| Transcript_442775 | Y+L amino acid transporter 2 | 2 | 2.24E-08 | 577249 |
| Transcript_217735 | protein RRP5 homolog\|programmed cell death 11 | 3 | 2.75E-08 | 584784 |
| Transcript_298943 | LOW QUALITY PROTEIN: small nuclear ribonucleoprotein F | 3 | 2.75E-08 | 592177 |

| | | | | |
|---|---|---|---|---|
| Transcript_276778 | electron transfer flavoprotein beta subunit lysine methyltransferase\|methyltransferase-like pro | 3 | 2.88E-08 | 589785 |
| Transcript_309454 | microtubule-actin cross-linking factor 1\|microtubule-actin cross-linking factor 1, isoforms 1/2/3 | 1 | 2.94E-08 | 756550 |
| Transcript_395351 | protein CDV3 homolog | 2 | 3.00E-08 | 593799 |
| Transcript_271655 | RRP12-like protein | 2 | 3.25E-08 | 587536 |
| Transcript_444753 | asparagine--tRNA ligase, cytoplasmic | 2 | 3.27E-08 | 574833 |
| Transcript_428848 | 26S proteasome non-ATPase regulatory subunit 6\|proteasome (prosome, macropain) 26S subu | 3 | 3.43E-08 | 590942 |
| Transcript_431583 | - | 3 | 3.70E-08 | 100891280 |
| Transcript_240340 | tRNA methyltransferase 10 homolog A | 3 | 3.72E-08 | 100889803 |
| Transcript_199752 | echinoidin-like | 1 | 3.72E-08 | 105446729 |
| Transcript_337159 | ergothioneine biosynthesis protein 1\|meiotically up-regulated gene 158 protein | 1 | 4.12E-08 | 584365 |
| Transcript_282269 | dipeptidase 1 | 1 | 4.12E-08 | 592585 |
| Transcript_393109 | ran-specific GTPase-activating protein | 2 | 4.20E-08 | 587798 |
| Transcript_342318 | aldehyde dehydrogenase, mitochondrial | 1 | 4.23E-08 | 581706 |
| Transcript_326720 | U3 small nucleolar ribonucleoprotein protein IMP3 | 2 | 4.47E-08 | 590066 |
| Transcript_214014 | glutamate-rich WD repeat-containing protein 1 | 3 | 4.70E-08 | 581222 |
| Transcript_240901 | suppressor of SWI4 1 homolog | 2 | 4.85E-08 | 582546 |
| Transcript_306776 | eukaryotic translation initiation factor 5B | 3 | 5.25E-08 | 763128 |
| Transcript_404542 | uncharacterized protein LOC576204 | 4 | 5.64E-08 | 576204 |
| Transcript_221851 | deleted in malignant brain tumors 1 protein-like | 1 | 5.78E-08 | 105440890 |
| Transcript_322501 | protein FAM199X | 3 | 5.93E-08 | 100893786 |
| Transcript_344436 | replication termination factor 2\|UPF0549 protein C20orf43 homolog\|protein RTF2 homolog | 2 | 5.99E-08 | 754268 |
| Transcript_404038 | 39S ribosomal protein L19, mitochondrial | 3 | 6.18E-08 | 589881 |
| Transcript_283496 | mitochondrial import inner membrane translocase subunit Tim23-like | 3 | 6.32E-08 | 105440270 |
| Transcript_376962 | lambda-crystallin homolog | 2 | 6.42E-08 | 589657 |
| Transcript_265994 | nascent polypeptide-associated complex subunit alpha | 2 | 7.11E-08 | 576369 |
| Transcript_311969 | uncharacterized protein RAB5IF homolog\|uncharacterized protein C20orf24 homolog | 3 | 7.24E-08 | 576217 |
| Transcript_340792 | WD repeat-containing protein 82 | 3 | 7.30E-08 | 100888929 |
| Transcript_219894 | serine/arginine repetitive matrix protein 2 | 1 | 7.35E-08 | 100888995 |
| Transcript_438464 | FAD synthase | 2 | 7.76E-08 | 593348 |
| Transcript_311263 | eukaryotic translation initiation factor 3 subunit M | 2 | 7.76E-08 | 589955 |
| Transcript_323033 | tubulin beta chain | 2 | 8.07E-08 | 373275 |
| Transcript_359669 | T-complex protein 1 subunit epsilon\|chaperonin containing TCP1, subunit 5 (epsilon) | 2 | 8.22E-08 | 575808 |
| Transcript_215999 | elongation factor Ts, mitochondrial | 3 | 9.32E-08 | 578811 |
| Transcript_430621 | alkylglycerol monooxygenase | 4 | 9.45E-08 | 579307 |
| Transcript_229332 | ADP-ribose pyrophosphatase, mitochondrial | 4 | 1.04E-07 | 756760 |
| Transcript_457631 | zinc finger protein 330 homolog | 2 | 1.05E-07 | 579935 |

| | | | | |
|---|---|---|---|---|
| Transcript_375389 | N-terminal Xaa-Pro-Lys N-methyltransferase 1 | 3 | 1.06E-07 | 579562 |
| Transcript_322851 | nucleolar and coiled-body phosphoprotein 1 | 3 | 1.07E-07 | 591574 |
| Transcript_225582 | ADP-ribosylation factor | 2 | 1.25E-07 | 763814 |
| Transcript_205826 | protein ABHD14B\|abhydrolase domain-containing protein 14A-like\|alpha/beta hydrolase doma | 1 | 1.26E-07 | 585657 |
| Transcript_448021 | proteasome subunit alpha type-6 | 3 | 1.28E-07 | 589393 |
| Transcript_434118 | voltage dependent calcium channel L-type | 1 | 1.32E-07 | 578128 |
| Transcript_331800 | uncharacterized protein LOC592324 | 2 | 1.33E-07 | 592324 |
| Transcript_365923 | insulin-like growth factor-binding protein complex acid labile subunit | 1 | 1.33E-07 | 105447591 |
| Transcript_403056 | UTP15, U3 small nucleolar ribonucleoprotein, homolog (S. cerevisiae)-like | 3 | 1.33E-07 | 754239 |
| Transcript_205321 | leucine-rich PPR motif-containing protein, mitochondrial | 2 | 1.39E-07 | 755486 |
| Transcript_291904 | uncharacterized protein LOC105438010\|Glycoside hydrolase family 31 domain containing prot | 4 | 1.47E-07 | 105438010 |
| Transcript_394693 | folylpolyglutamate synthase, mitochondrial | 2 | 1.49E-07 | 592205 |
| Transcript_50290 | uncharacterized protein LOC100890372 | 1 | 1.53E-07 | 100890372 |
| Transcript_216838 | LOW QUALITY PROTEIN: lipase maturation factor 2 | 4 | 1.57E-07 | 583944 |
| Transcript_434160 | ubiquitin-conjugating enzyme E2 N | 3 | 1.57E-07 | 590968 |
| Transcript_223216 | uncharacterized protein LOC100892961 | 4 | 1.58E-07 | 100892961 |
| Transcript_385327 | amidophosphoribosyltransferase | 2 | 1.71E-07 | 582049 |
| Transcript_316114 | eukaryotic translation initiation factor 2 subunit 3\|eukaryotic translation initiation factor 2 subu | 3 | 1.91E-07 | 582011 |
| Transcript_364746 | insulin-like growth factor 2 mRNA-binding protein 1 | 3 | 1.93E-07 | 576835 |
| Transcript_334779 | 3'(2'),5'-bisphosphate nucleotidase 1 | 4 | 1.94E-07 | 580070 |
| Transcript_185684 | nucleolar protein 6 | 2 | 1.95E-07 | 577631 |
| Transcript_196408 | DNA mismatch repair protein Mlh3 | 2 | 1.97E-07 | 575007 |
| Transcript_443664 | acid phosphatase type 7\|iron/zinc purple acid phosphatase-like protein | 1 | 2.13E-07 | 580652 |
| Transcript_429248 | histone-binding protein RBBP4\|retinoblastoma binding protein 4 | 3 | 2.13E-07 | 579451 |
| Transcript_351461 | 28S ribosomal protein S22, mitochondrial | 2 | 2.16E-07 | 589317 |
| Transcript_454778 | protein LLP homolog | 2 | 2.26E-07 | 764101 |
| Transcript_349228 | uncharacterized protein LOC591826 | 2 | 2.30E-07 | 591826 |
| Transcript_187299 | transcription initiation protein SPT3 homolog | 2 | 2.34E-07 | 586422 |
| Transcript_361302 | sideroflexin-5 | 1 | 2.64E-07 | 577642 |
| Transcript_416648 | nucleolar protein 14 | 3 | 2.78E-07 | 578764 |
| Transcript_225955 | endoribonuclease YbeY\|putative ribonuclease\|rRNA maturation factor homolog | 1 | 2.82E-07 | 588773 |
| Transcript_267958 | regulator of chromosome condensation | 3 | 2.90E-07 | 754700 |
| Transcript_352669 | zinc finger protein 593 | 2 | 2.96E-07 | 592435 |
| Transcript_384255 | zinc finger protein 362 | 2 | 2.99E-07 | 583226 |
| Transcript_422610 | hydroxyacid-oxoacid transhydrogenase, mitochondrial | 1 | 3.05E-07 | 575433 |
| Transcript_424059 | acetyl-coenzyme A transporter 1 | 2 | 3.05E-07 | 589856 |

| | | | | |
|---|---|---|---|---|
| Transcript_332395 | uncharacterized protein LOC591223 | 5 | 3.06E-07 | 591223 |
| Transcript_225467 | heat shock protein 75 kDa, mitochondrial | 3 | 3.10E-07 | 578227 |
| Transcript_444809 | eukaryotic peptide chain release factor subunit 1 | 3 | 3.11E-07 | 591483 |
| Transcript_195493 | CTP synthase 1 | 2 | 3.11E-07 | 575088 |
| Transcript_428926 | menin | 2 | 3.22E-07 | 589807 |
| Transcript_197652 | LOW QUALITY PROTEIN: tubulin alpha-1C chain | 2 | 3.23E-07 | 762594 |
| Transcript_203588 | DDB1- and CUL4-associated factor 13 | 3 | 3.39E-07 | 589486 |
| Transcript_280848 | nudC domain-containing protein 1 | 2 | 3.39E-07 | 590657 |
| Transcript_429226 | uncharacterized protein LOC590843 | 4 | 3.46E-07 | 590843 |
| Transcript_451457 | PHD finger protein 24 | 3 | 3.49E-07 | 100892446 |
| Transcript_343741 | proteasome maturation protein | 3 | 3.84E-07 | 579892 |
| Transcript_185040 | endothelin-converting enzyme 1 | 1 | 3.84E-07 | 755783 |
| Transcript_282876 | spermidine synthase | 2 | 4.12E-07 | 591935 |
| Transcript_427763 | major facilitator superfamily domain-containing protein 9 | 2 | 4.19E-07 | 577847 |
| Transcript_425248 | T-complex protein 1 subunit gamma | 3 | 4.26E-07 | 574601 |
| Transcript_286893 | proteasome subunit beta type-1 | 2 | 4.31E-07 | 754955 |
| Transcript_323172 | sialic acid synthase\|N-acetylneuraminic acid synthase | 2 | 4.43E-07 | 577884 |
| Transcript_373212 | fibrillin-1\|latent-transforming growth factor beta-binding protein 4 | 1 | 4.48E-07 | 100887851 |
| Transcript_177898 | sodium- and chloride-dependent GABA transporter 1 | 1 | 4.60E-07 | 581958 |
| Transcript_377252 | neural-cadherin | 1 | 4.62E-07 | 589984 |
| Transcript_364897 | FAST kinase domain-containing protein 4\|protein TBRG4 | 3 | 4.73E-07 | 580380 |
| Transcript_238012 | G patch domain-containing protein 4\|protein FAM133-like | 3 | 4.75E-07 | 581552 |
| Transcript_240840 | uncharacterized protein LOC105439633\|Autophagy-related protein 27 | 1 | 4.78E-07 | 105439633 |
| Transcript_359750 | glucosamine-6-phosphate isomerase 2 | 3 | 5.05E-07 | 585301 |
| Transcript_309937 | solute carrier organic anion transporter family member 2A1-like | 2 | 5.28E-07 | 752367 |
| Transcript_208822 | lysophosphatidylserine lipase ABHD12\|monoacylglycerol lipase ABHD12 | 2 | 5.35E-07 | 100891460 |
| Transcript_418613 | PIN2/TERF1-interacting telomerase inhibitor 1 | 2 | 5.37E-07 | 591382 |
| Transcript_294332 | 60S ribosome subunit biogenesis protein NIP7 homolog | 2 | 5.43E-07 | 579986 |
| Transcript_353843 | poly [ADP-ribose] polymerase tankyrase\|poly(ADP-ribose) polymerase pme-5 | 4 | 5.86E-07 | 588883 |
| Transcript_205105 | uncharacterized protein LOC578177\|low-density lipoprotein receptor-related protein 1B | 1 | 5.99E-07 | 578177 |
| Transcript_364601 | acetolactate synthase-like protein | 2 | 6.53E-07 | 593426 |
| Transcript_321936 | RNA-binding protein 34 | 3 | 6.77E-07 | 576551 |
| Transcript_367437 | secreted frizzled-related protein 3-like | 3 | 6.84E-07 | 581781 |
| Transcript_367877 | vesicular integral-membrane protein VIP36\|VIP36-like protein | 3 | 7.04E-07 | 578461 |
| Transcript_399833 | serine/threonine-protein phosphatase CPPED1\|calcineurin-like phosphoesterase domain-conta | 2 | 7.07E-07 | 583499 |
| Transcript_323748 | cryptochrome-1 | 1 | 7.23E-07 | 580742 |

| | | | | |
|---|---|---|---|---|
| Transcript_394678 | di-N-acetylchitobiase | 2 | 7.28E-07 | 578451 |
| Transcript_232547 | protein LMBR1L | 3 | 7.31E-07 | 578121 |
| Transcript_340709 | DNA polymerase beta | 2 | 7.47E-07 | 582628 |
| Transcript_393067 | signal peptide, CUB and EGF-like domain-containing protein 1 | 1 | 8.42E-07 | 577317 |
| Transcript_262552 | cytochrome P450 3A9 | 1 | 8.45E-07 | 588156 |
| Transcript_226152 | myosin-16\|myosin-6 | 1 | 8.72E-07 | 580674 |
| Transcript_367772 | tubulin alpha-1 chain | 2 | 8.81E-07 | 105439767 |
| Transcript_204519 | mitochondrial-processing peptidase subunit alpha\|peptidase (mitochondrial processing) alpha | 3 | 9.15E-07 | 100889464 |
| Transcript_6160 | 60S ribosomal protein L27a | 2 | 9.97E-07 | 580148 |
| Transcript_228286 | elongation factor Tu, mitochondrial | 3 | 1.02E-06 | 583958 |
| Transcript_320536 | 4-hydroxyphenylpyruvate dioxygenase-like protein | 3 | 1.03E-06 | 576414 |
| Transcript_290528 | TEF | 3 | 1.06E-06 | 580363 |
| Transcript_359106 | GPI ethanolamine phosphate transferase 3 | 3 | 1.10E-06 | 584071 |
| Transcript_302349 | bystin | 2 | 1.11E-06 | 578981 |
| Transcript_260705 | sorbin and SH3 domain-containing protein 1\|vinexin | 1 | 1.13E-06 | 574636 |
| Transcript_205888 | CDGSH iron-sulfur domain-containing protein 2 homolog A | 1 | 1.13E-06 | 586672 |
| Transcript_279040 | KLF13 | 2 | 1.17E-06 | 576795 |
| Transcript_230031 | importin-7 | 2 | 1.18E-06 | 586884 |
| Transcript_365373 | ankyrin repeat and protein kinase domain-containing protein 1 | 3 | 1.24E-06 | 100889589 |
| Transcript_278797 | protein transport protein Sec24A\|basic proline-rich protein-like | 2 | 1.25E-06 | 579127 |
| Transcript_305867 | actin-5C | 2 | 1.26E-06 | 581650 |
| Transcript_303652 | derlin-1 | 3 | 1.30E-06 | 592785 |
| Transcript_344547 | T-complex protein 1 subunit beta | 2 | 1.33E-06 | 580864 |
| Transcript_366717 | uncharacterized protein LOC582810\|Tudor domain-containing protein | 2 | 1.33E-06 | 582810 |
| Transcript_358088 | fucose mutarotase | 4 | 1.37E-06 | 100888310 |
| Transcript_329212 | eukaryotic translation initiation factor 3 subunit J | 2 | 1.40E-06 | 578606 |
| Transcript_344098 | nucleolar GTP-binding protein 1\|GTP binding protein 4 | 3 | 1.40E-06 | 575126 |
| Transcript_374375 | X-ray radiation resistance-associated protein 1 | 3 | 1.45E-06 | 584905 |
| Transcript_198110 | DNA-directed RNA polymerases I and III subunit RPAC1 | 3 | 1.45E-06 | 590827 |
| Transcript_409226 | CD151 antigen | 4 | 1.46E-06 | 592232 |
| Transcript_185863 | interaptin\|nuclear anchorage protein 1\|nucleoprotein TPR\|uncharacterized protein LOC58842 | 1 | 1.63E-06 | 588428 |
| Transcript_411421 | wnt inhibitory factor 1 | 1 | 1.70E-06 | 100892419 |
| Transcript_393713 | multifunctional protein ADE2 | 2 | 1.76E-06 | 575874 |
| Transcript_212609 | NHL repeat-containing protein 2 | 2 | 1.83E-06 | 591178 |
| Transcript_441990 | uncharacterized protein LOC100888048 | 1 | 1.85E-06 | 100888048 |
| Transcript_243213 | protein-serine O-palmitoleoyltransferase porcupine\|protein-cysteine N-palmitoyltransferase po | 2 | 1.98E-06 | 577959 |

| | | | | |
|---|---|---|---|---|
| Transcript_249746 | DNA-directed RNA polymerase, mitochondrial | 2 | 1.98E-06 | 762772 |
| Transcript_286756 | GDP-fucose protein O-fucosyltransferase 1 | 3 | 2.03E-06 | 585387 |
| Transcript_235731 | MAM and LDL-receptor class A domain-containing protein 2 | 1 | 2.05E-06 | 755161 |
| Transcript_440018 | nucleolar complex protein 4 homolog | 2 | 2.11E-06 | 591488 |
| Transcript_341101 | A disintegrin and metalloproteinase with thrombospondin motifs 18\|ADAM metallopeptidase w | 3 | 2.36E-06 | 574964 |
| Transcript_180918 | phosphopantothenoylcysteine decarboxylase subunit SIS2-like | 1 | 2.38E-06 | 105438224 |
| Transcript_439703 | protein CBFA2T1\|protein CBFA2T3-like | 1 | 2.40E-06 | 576535 |
| Transcript_405538 | uncharacterized protein LOC100891656 | 1 | 2.45E-06 | 100891656 |
| Transcript_292194 | FERM domain-containing protein 1 | 1 | 2.46E-06 | 592633 |
| Transcript_110850 | 60S ribosomal protein L3 | 2 | 2.89E-06 | 586477 |
| Transcript_434866 | uncharacterized protein LOC589939\|Harbinger transposase-derived nuclease domain-containir | 3 | 2.92E-06 | 589939 |
| Transcript_438152 | DNA excision repair protein ERCC-6-like\|transcriptional regulator ATRX | 1 | 2.95E-06 | 575953 |
| Transcript_446005 | 2-oxoglutarate-dependent dioxygenase htyE\|UPF0676 protein C1494.01 | 1 | 3.10E-06 | 752472 |
| Transcript_286899 | uncharacterized protein LOC587999 | 2 | 3.21E-06 | 587999 |
| Transcript_325335 | uncharacterized protein LOC593876 | 1 | 3.21E-06 | 593876 |
| Transcript_249330 | eukaryotic translation initiation factor 3 subunit H-B | 2 | 3.27E-06 | 575080 |
| Transcript_443947 | uncharacterized protein LOC579607\|putative oxidoreductase YteT-like | 1 | 3.33E-06 | 579607 |
| Transcript_190658 | pyrroline-5-carboxylate reductase 3 | 2 | 3.41E-06 | 586502 |
| Transcript_443207 | Bardet-Biedl syndrome 1 protein | 2 | 3.41E-06 | 578507 |
| Transcript_350598 | RNA 3'-terminal phosphate cyclase-like protein | 2 | 3.67E-06 | 592597 |
| Transcript_239281 | cholinesterase 1\|thyroglobulin | 1 | 3.85E-06 | 583460 |
| Transcript_367596 | transitional endoplasmic reticulum ATPase | 3 | 3.99E-06 | 575609 |
| Transcript_381250 | prostaglandin E synthase 2 | 3 | 4.04E-06 | 584887 |
| Transcript_409717 | isochorismatase domain-containing protein 2\|isochorismatase domain-containing protein 2, mi | 4 | 4.04E-06 | 590365 |
| Transcript_185238 | multifunctional methyltransferase subunit TRM112-like protein\|tRNA methyltransferase 112 ho | 2 | 4.21E-06 | 579707 |
| Transcript_444350 | melanotransferrin | 3 | 4.37E-06 | 581063 |
| Transcript_306787 | methyltransferase-like protein 24 | 1 | 4.40E-06 | 100891683 |
| Transcript_271897 | HEAT repeat-containing protein 1 | 3 | 4.60E-06 | 589888 |
| Transcript_302913 | L-lactate dehydrogenase | 1 | 4.65E-06 | 586683 |
| Transcript_386663 | pancreatic lipase-related protein 2 | 1 | 4.67E-06 | 592851 |
| Transcript_432931 | NADPH oxidase 5 | 1 | 4.74E-06 | 752031 |
| Transcript_306891 | graves disease carrier protein homolog | 2 | 4.80E-06 | 584753 |
| Transcript_382713 | 60S ribosomal protein L19 | 2 | 4.80E-06 | 593861 |
| Transcript_416522 | uncharacterized protein LOC100888035 | 2 | 4.85E-06 | 100888035 |
| Transcript_431677 | coatomer subunit beta' | 2 | 4.95E-06 | 589321 |
| Transcript_185030 | DNA-dependent protein kinase catalytic subunit | 1 | 5.16E-06 | 586799 |

| | | | | |
|---|---|---|---|---|
| Transcript_381229 | Golgi SNAP receptor complex member 1 | 2 | 5.23E-06 | 593134 |
| Transcript_301702 | unconventional myosin-XVI\|formin-like protein 20\|myosin XVI\|putative F-box protein At1g473 | 1 | 5.29E-06 | 587143 |
| Transcript_197661 | LOW QUALITY PROTEIN: tubulin alpha-1 chain | 2 | 5.35E-06 | 588319 |
| Transcript_206262 | cleavage stimulation factor subunit 2 | 2 | 5.44E-06 | 753333 |
| Transcript_316435 | neprilysin-1\|endothelin-converting enzyme 1 | 1 | 5.66E-06 | 580325 |
| Transcript_266841 | tuftelin | 4 | 5.71E-06 | 576303 |
| Transcript_319974 | uncharacterized protein LOC100890537 | 1 | 5.79E-06 | 100890537 |
| Transcript_223102 | transducin beta-like protein 3 | 3 | 6.10E-06 | 100891641 |
| Transcript_234991 | ADAMTS-like protein 1 | 2 | 6.11E-06 | 580296 |
| Transcript_277138 | uncharacterized protein LOC100891149 | 3 | 6.28E-06 | 100891149 |
| Transcript_43695 | b(0,+)-type amino acid transporter 1\|B(0,+)-type amino acid transporter 1-like | 1 | 6.33E-06 | 579553 |
| Transcript_447128 | maltase 1\|alpha-glucosidase | 1 | 6.40E-06 | 583103 |
| Transcript_190954 | uncharacterized protein LOC593642\|CAP domain-containing protein | 4 | 6.55E-06 | 593642 |
| Transcript_84237 | actin CyI, cytoplasmic | 1 | 6.60E-06 | 592912 |
| Transcript_412252 | protein SEC13 homolog | 3 | 6.76E-06 | 583776 |
| Transcript_284067 | programmed cell death protein 2-like | 2 | 6.81E-06 | 576484 |
| Transcript_262309 | N-alpha-acetyltransferase 11 | 2 | 6.82E-06 | 580091 |
| Transcript_396324 | phosphopantothenate--cysteine ligase | 3 | 6.92E-06 | 763180 |
| Transcript_302120 | cadherin-23\|fat-like cadherin-related tumor suppressor homolog\|protocadherin Fat 1 | 1 | 6.98E-06 | 594286 |
| Transcript_433507 | MKX | 2 | 7.15E-06 | 105436421 |
| Transcript_205651 | lysophosphatidylcholine acyltransferase 2 | 1 | 7.37E-06 | 579107 |
| Transcript_202496 | ribosome biogenesis protein BRX1 homolog | 3 | 7.55E-06 | 583114 |
| Transcript_425468 | dnaJ homolog subfamily B member 1 | 2 | 7.82E-06 | 752244 |
| Transcript_315415 | putative uncharacterized protein CXorf58 | 2 | 7.85E-06 | 593302 |
| Transcript_244512 | aspartate--tRNA ligase, cytoplasmic | 3 | 8.02E-06 | 588558 |
| Transcript_272010 | proto-oncogene Wnt-3 | 3 | 8.05E-06 | 585683 |
| Transcript_285582 | splicing factor 3A subunit 2 | 2 | 8.28E-06 | 576257 |
| Transcript_438398 | U2 small nuclear ribonucleoprotein B''\|U1 small nuclear ribonucleoprotein A-like | 3 | 8.35E-06 | 584276 |
| Transcript_289715 | uncharacterized protein LOC763307\|ZU5 domain-containing protein | 3 | 8.56E-06 | 763307 |
| Transcript_328267 | 40S ribosomal protein S26 | 2 | 8.78E-06 | 576654 |
| Transcript_356949 | ubiquitin-conjugating enzyme E2 K | 2 | 9.01E-06 | 585095 |
| Transcript_440889 | serine/threonine-protein phosphatase 4 catalytic subunit | 2 | 9.06E-06 | 594644 |
| Transcript_405218 | 40S ribosomal protein S11 | 2 | 9.09E-06 | 579026 |
| Transcript_375963 | uncharacterized protein LOC581850 | 1 | 9.12E-06 | 581850 |
| Transcript_299201 | mesencephalic astrocyte-derived neurotrophic factor homolog | 3 | 9.30E-06 | 755636 |
| Transcript_215281 | decaprenyl-diphosphate synthase subunit 2 | 2 | 9.56E-06 | 580439 |

| | | | | |
|---|---|---|---|---|
| Transcript_368597 | ubiquitin carboxyl-terminal hydrolase 14 | 2 | 1.00E-05 | 581896 |
| Transcript_346637 | nuclear migration protein nudC | 2 | 1.01E-05 | 589717 |
| Transcript_351605 | 26S proteasome non-ATPase regulatory subunit 10 | 3 | 1.02E-05 | 576864 |
| Transcript_244261 | acidic leucine-rich nuclear phosphoprotein 32 family member E | 2 | 1.02E-05 | 100892466 |
| Transcript_440816 | tyrosine aminotransferase | 2 | 1.02E-05 | 592114 |
| Transcript_381701 | rho-related BTB domain-containing protein 1 | 1 | 1.02E-05 | 575159 |
| Transcript_445237 | 39S ribosomal protein S30, mitochondrial\|28S ribosomal protein S30, mitochondrial | 3 | 1.06E-05 | 591808 |
| Transcript_298879 | DNA-directed RNA polymerase III subunit RPC6 | 2 | 1.08E-05 | 580860 |
| Transcript_275685 | uncharacterized protein LOC587541 | 4 | 1.09E-05 | 587541 |
| Transcript_381489 | papilin | 1 | 1.12E-05 | 584177 |
| Transcript_362661 | ribosome-recycling factor, mitochondrial | 3 | 1.13E-05 | 587941 |
| Transcript_179642 | tetraspanin-5 | 1 | 1.14E-05 | 582219 |
| Transcript_404610 | uncharacterized protein F13E9.13, mitochondrial\|Membrane complex biogenesis protein, BtpA | 3 | 1.14E-05 | 579795 |
| Transcript_310943 | tryptophan--tRNA ligase, cytoplasmic | 2 | 1.16E-05 | 574927 |
| Transcript_180218 | proton-coupled amino acid transporter 1 | 4 | 1.20E-05 | 587558 |
| Transcript_277419 | protein SET | 2 | 1.23E-05 | 583082 |
| Transcript_316855 | deleted in malignant brain tumors 1 protein\|galectin-3-binding protein pseudogene\|scavenger | 1 | 1.25E-05 | 754160 |
| Transcript_217175 | mitochondrial import inner membrane translocase subunit Tim22 | 3 | 1.27E-05 | 592058 |
| Transcript_271076 | carbohydrate sulfotransferase 15 | 4 | 1.34E-05 | 582046 |
| Transcript_434835 | alpha-aminoadipic semialdehyde dehydrogenase | 1 | 1.34E-05 | 588376 |
| Transcript_412907 | tubulin alpha-2/alpha-4 chain | 2 | 1.35E-05 | 582621 |
| Transcript_429645 | N-acetylated-alpha-linked acidic dipeptidase 2-like | 1 | 1.36E-05 | 583418 |
| Transcript_235859 | sucrase-isomaltase, intestinal | 1 | 1.40E-05 | 588081 |
| Transcript_267184 | 28S ribosomal protein S5, mitochondrial | 3 | 1.43E-05 | 583862 |
| Transcript_380459 | cryptochrome-2 | 4 | 1.46E-05 | 583959 |
| Transcript_453750 | uncharacterized protein LOC581715\|probable enoyl-CoA hydratase echA8-like\|probable enoyl- | 4 | 1.49E-05 | 581715 |
| Transcript_213782 | nicalin-1 | 3 | 1.50E-05 | 587749 |
| Transcript_323832 | aminoacylase-1 | 1 | 1.70E-05 | 577570 |
| Transcript_402336 | mitochondrial ribonuclease P catalytic subunit\|mitochondrial ribonuclease P protein 3 | 3 | 1.71E-05 | 578934 |
| Transcript_359571 | LOW QUALITY PROTEIN: uncharacterized protein LOC577952 | 1 | 1.71E-05 | 577952 |
| Transcript_189718 | MYBBP1A | 2 | 1.71E-05 | 583345 |
| Transcript_388247 | mitochondrial glutamate carrier 2 | 2 | 1.74E-05 | 753367 |
| Transcript_369099 | protein Abitram\|protein FAM206A-like\|protein Simiate | 3 | 1.75E-05 | 757335 |
| Transcript_305359 | metabotropic glutamate receptor 1\|extracellular calcium-sensing receptor-like | 3 | 1.75E-05 | 577623 |
| Transcript_456418 | ficolin-2-like | 1 | 1.80E-05 | 583514 |
| Transcript_433359 | probable serine/threonine-protein kinase drkD\|putative serine/threonine-protein kinase drkD | 2 | 1.85E-05 | 577165 |

| | | | | |
|---|---|---|---|---|
| Transcript_360852 | ribosomal RNA-processing protein 7 homolog A | 3 | 1.88E-05 | 580493 |
| Transcript_236563 | protein FAM166B-like | 5 | 1.92E-05 | 105439658 |
| Transcript_322912 | 39S ribosomal protein L13, mitochondrial | 3 | 2.04E-05 | 585375 |
| Transcript_373532 | putative helicase MOV-10 | 3 | 2.05E-05 | 583129 |
| Transcript_374213 | uncharacterized protein LOC763830 | 3 | 2.06E-05 | 763830 |
| Transcript_442862 | zinc finger CCCH domain-containing protein 15 | 2 | 2.07E-05 | 580460 |
| Transcript_401581 | protein canopy homolog 2 | 3 | 2.08E-05 | 588644 |
| Transcript_453440 | glutamate receptor 2 | 1 | 2.10E-05 | 591929 |
| Transcript_343792 | elongation factor 1-delta-like | 3 | 2.16E-05 | 575168 |
| Transcript_371946 | uncharacterized protein LOC100889820 | 4 | 2.23E-05 | 100889820 |
| Transcript_337768 | coiled-coil domain-containing protein 58 | 3 | 2.23E-05 | 576818 |
| Transcript_230454 | exportin-5 | 3 | 2.31E-05 | 592021 |
| Transcript_314891 | periodic tryptophan protein 2 homolog | 3 | 2.43E-05 | 752047 |
| Transcript_321961 | organic cation transporter protein | 1 | 2.46E-05 | 582688 |
| Transcript_299268 | 39S ribosomal protein L52, mitochondrial | 3 | 2.49E-05 | 752505 |
| Transcript_227249 | cytochrome P450 2J6 | 1 | 2.49E-05 | 575697 |
| Transcript_329073 | uncharacterized protein LOC100889633 | 2 | 2.50E-05 | 100889633 |
| Transcript_449172 | transcription initiation factor IIA subunit 1 | 2 | 2.52E-05 | 100888038 |
| Transcript_66007 | cytospin-A | 1 | 2.56E-05 | 583396 |
| Transcript_423340 | inverted formin-2 | 1 | 2.58E-05 | 579910 |
| Transcript_423272 | phe13-bombesin receptor | 1 | 2.75E-05 | 100888815 |
| Transcript_276375 | uncharacterized protein LOC584281 | 1 | 2.75E-05 | 584281 |
| Transcript_243879 | carbonyl reductase [NADPH] 1 | 4 | 2.76E-05 | 574551 |
| Transcript_310622 | actin related protein 1 | 1 | 2.83E-05 | 373192 |
| Transcript_380140 | cardiolipin synthase (CMP-forming)\|probable cardiolipin synthase (CMP-forming) | 2 | 2.86E-05 | 587370 |
| Transcript_378334 | DNA-directed RNA polymerase II subunit RPB7 | 3 | 2.88E-05 | 752151 |
| Transcript_298921 | development-specific protein LVN1.2 | 1 | 2.90E-05 | 100888941 |
| Transcript_393283 | LOW QUALITY PROTEIN: xanthine dehydrogenase/oxidase | 1 | 3.15E-05 | 576712 |
| Transcript_388184 | serine/arginine repetitive matrix protein 1 | 1 | 3.16E-05 | 575766 |
| Transcript_401245 | uncharacterized protein LOC593760 | 4 | 3.16E-05 | 593760 |
| Transcript_434004 | hydroxyacyl-coenzyme A dehydrogenase, mitochondrial | 3 | 3.18E-05 | 582126 |
| Transcript_384548 | salivary glue protein Sgs-3 | 3 | 3.18E-05 | 100892638 |
| Transcript_380262 | cytochrome P450 4V2 | 4 | 3.20E-05 | 581875 |
| Transcript_42693 | 40S ribosomal protein S13 | 2 | 3.25E-05 | 581196 |
| Transcript_206502 | nucleolar protein 56\|NOP56 ribonucleoprotein homolog | 2 | 3.29E-05 | 590067 |
| Transcript_195549 | sodium-dependent phosphate transporter 2 | 2 | 3.35E-05 | 582615 |

| | | | | |
|---|---|---|---|---|
| Transcript_331211 | probable Na(+)/H(+) antiporter nhx-9 | 1 | 3.38E-05 | 591586 |
| Transcript_372883 | protein dispatched homolog 1-like | 1 | 3.39E-05 | 580689 |
| Transcript_308094 | sodium-dependent dopamine transporter\|solute carrier family 6 (neurotransmitter transporter | 1 | 3.48E-05 | 584179 |
| Transcript_438519 | degenerin deg-1 | 1 | 3.53E-05 | 100891272 |
| Transcript_307696 | ATP-binding cassette sub-family B member 8, mitochondrial | 3 | 3.69E-05 | 754756 |
| Transcript_370380 | serine/threonine-protein kinase NLK\|serine/threonine-protein kinase NLK2 | 2 | 3.73E-05 | 574783 |
| Transcript_402303 | GTP 3',8-cyclase, mitochondrial\|cyclic pyranopterin monophosphate synthase, mitochondrial\|n | 4 | 3.77E-05 | 578700 |
| Transcript_340758 | proteasome subunit alpha type-2 | 3 | 3.77E-05 | 579111 |
| Transcript_423204 | FOXA | 1 | 3.80E-05 | 578584 |
| Transcript_443748 | pyridine nucleotide-disulfide oxidoreductase domain-containing protein 2\|pyridine nucleotide- | 4 | 3.86E-05 | 587519 |
| Transcript_426155 | mitotic apparatus protein p62 | 3 | 3.98E-05 | 105443132 |
| Transcript_373050 | dihydropteridine reductase | 1 | 4.06E-05 | 575893 |
| Transcript_268252 | 39S ribosomal protein L47, mitochondrial | 3 | 4.15E-05 | 588415 |
| Transcript_261599 | elongation factor 1 alpha | 2 | 4.40E-05 | 548620 |
| Transcript_315154 | peroxisomal membrane protein 11A | 1 | 4.41E-05 | 576910 |
| Transcript_254082 | probable ATP-dependent RNA helicase DDX56 | 2 | 4.51E-05 | 585968 |
| Transcript_394798 | band 7 protein AGAP004871-like | 1 | 4.55E-05 | 578433 |
| Transcript_253869 | uncharacterized protein LOC100889419\|IgGFc-binding protein, N-terminal domain containing p | 1 | 4.65E-05 | 100889419 |
| Transcript_258821 | transcription initiation factor TFIID subunit 10 | 2 | 4.67E-05 | 752563 |
| Transcript_433800 | proteasome subunit alpha type-7 | 2 | 4.80E-05 | 579458 |
| Transcript_459696 | MYNN | 2 | 4.88E-05 | 100893972 |
| Transcript_347359 | ectonucleotide pyrophosphatase/phosphodiesterase family member 7-like | 5 | 4.90E-05 | 577518 |
| Transcript_355848 | NACHT domain- and WD repeat-containing protein 1\|NACHT and WD repeat domain-containing | 1 | 4.97E-05 | 591208 |
| Transcript_196171 | peptidyl-prolyl cis-trans isomerase | 2 | 5.02E-05 | 585216 |
| Transcript_139365 | deleted in malignant brain tumors 1 protein | 1 | 5.10E-05 | 753862 |
| Transcript_267269 | 39S ribosomal protein L23, mitochondrial | 2 | 5.29E-05 | 587414 |
| Transcript_219950 | citrate synthase, mitochondrial | 2 | 5.60E-05 | 590444 |
| Transcript_223195 | 15-hydroxyprostaglandin dehydrogenase [NAD(+)] | 4 | 5.63E-05 | 591898 |
| Transcript_230867 | RNA N6-adenosine-methyltransferase mettl16\|methyltransferase-like protein 16 | 3 | 5.67E-05 | 588854 |
| Transcript_279744 | eukaryotic translation initiation factor 4 gamma 3\|eukaryotic translation initiation factor 4 gam | 2 | 5.68E-05 | 588274 |
| Transcript_330100 | NRL/MAF | 1 | 5.85E-05 | 581716 |
| Transcript_254029 | uncharacterized protein LOC105441448 | 1 | 6.12E-05 | 105441448 |
| Transcript_431131 | proteasome subunit beta type-7 | 3 | 6.38E-05 | 588252 |
| Transcript_263253 | E3 ubiquitin-protein ligase TRIM23 | 1 | 6.38E-05 | 591350 |
| Transcript_364264 | DNA polymerase epsilon subunit 4 | 2 | 6.53E-05 | 763808 |
| Transcript_181686 | patched domain-containing protein 3 | 2 | 6.55E-05 | 100888465 |

| | | | | |
|---|---|---|---|---|
| Transcript_203172 | uncharacterized protein LOC577523\|malate dehydrogenase-like | 1 | 6.68E-05 | 577523 |
| Transcript_427147 | carbohydrate sulfotransferase 11 | 1 | 6.68E-05 | 589022 |
| Transcript_414827 | guanine nucleotide exchange factor MSS4 | 2 | 6.83E-05 | 591831 |
| Transcript_197695 | quinone oxidoreductase-like\|uncharacterized protein LOC764164\|Alcohol dehydrogenase, C-te | 2 | 7.08E-05 | 764164 |
| Transcript_269277 | ribosomal protein S14 | 2 | 7.19E-05 | 574895 |
| Transcript_246683 | PMS1 protein homolog 1 | 1 | 7.21E-05 | 592483 |
| Transcript_447058 | coiled-coil domain-containing protein 86\|uncharacterized protein LOC100889012 | 3 | 7.35E-05 | 100889012 |
| Transcript_428449 | COX assembly mitochondrial protein homolog | 3 | 7.46E-05 | 754298 |
| Transcript_373487 | leucine-rich repeat-containing protein 58 | 2 | 7.64E-05 | 592464 |
| Transcript_297301 | isthmin | 2 | 8.01E-05 | 762678 |
| Transcript_274172 | 60S ribosomal protein L31 | 2 | 8.02E-05 | 589085 |
| Transcript_214012 | RNA transcription, translation and transport factor protein\|UPF0568 protein C14orf166 homolo | 3 | 8.11E-05 | 586176 |
| Transcript_180063 | endothelin-converting enzyme homolog | 1 | 8.20E-05 | 594281 |
| Transcript_335203 | cyclin-dependent kinase 2 | 1 | 8.26E-05 | 585950 |
| Transcript_436916 | scavenger receptor cysteine-rich domain-containing group B protein | 1 | 8.28E-05 | 577634 |
| Transcript_168671 | EEF1A1 | 2 | 8.38E-05 | 548620 |
| Transcript_196792 | hydroxyproline dehydrogenase | 1 | 8.47E-05 | 576696 |
| Transcript_437063 | serine/threonine-protein kinase TBK1 | 3 | 8.56E-05 | 579073 |
| Transcript_269134 | octopamine receptor | 2 | 8.60E-05 | 100893874 |
| Transcript_280627 | methylenetetrahydrofolate reductase | 1 | 8.96E-05 | 589576 |
| Transcript_317810 | kin of IRRE-like protein 1 | 1 | 9.04E-05 | 762504 |
| Transcript_387048 | peptidyl-prolyl cis-trans isomerase-like 1 | 2 | 9.09E-05 | 575873 |
| Transcript_316602 | DNA mismatch repair protein Msh2\|mutS homolog 2, colon cancer, nonpolyposis type 1 | 1 | 9.13E-05 | 589363 |
| Transcript_263803 | phosphate carrier protein, mitochondrial | 2 | 9.29E-05 | 577743 |
| Transcript_249042 | thioredoxin-like protein 1 | 3 | 9.29E-05 | 581305 |
| Transcript_323338 | probable assembly chaperone of rpl4\|UPF0661 TPR repeat-containing protein C16D10.01c | 2 | 9.32E-05 | 100891594 |
| Transcript_218894 | 40S ribosomal protein S15a\|ribosomal protein S24 | 2 | 9.49E-05 | 373469 |
| Transcript_77032 | cubilin | 2 | 9.56E-05 | 580195 |
| Transcript_214740 | glycerophosphocholine phosphodiesterase GPCPD1 | 1 | 9.94E-05 | 594641 |
| Transcript_381580 | short-chain collagen C4 | 1 | 9.95E-05 | 100893425 |
| Transcript_373705 | tctex1 domain-containing protein 1 | 1 | 0.000100228 | 588988 |
| Transcript_255002 | protein giant | 2 | 0.000100627 | 105444027 |
| Transcript_328974 | GRAM domain-containing protein 4 | 1 | 0.00010098 | 581215 |
| Transcript_294422 | persulfide dioxygenase ETHE1, mitochondrial | 5 | 0.000102325 | 585574 |
| Transcript_203149 | DNA-directed RNA polymerases I, II, and III subunit RPABC1 | 3 | 0.00010449 | 578317 |
| Transcript_360539 | methionine adenosyltransferase | 3 | 0.00010822 | 548618 |

| | | | | |
|---|---|---|---|---|
| Transcript_73952 | aminoacyl tRNA synthase complex-interacting multifunctional protein 2 | 2 | 0.00010822 | 575132 |
| Transcript_303046 | U3 small nucleolar RNA-associated protein 14 homolog A | 3 | 0.000111727 | 585850 |
| Transcript_199821 | probable ATP-dependent RNA helicase DDX47 | 3 | 0.000112486 | 581057 |
| Transcript_175282 | uncharacterized protein LOC105442054 | 4 | 0.000113657 | 105442054 |
| Transcript_267731 | 39S ribosomal protein L12, mitochondrial | 3 | 0.000120962 | 585458 |
| Transcript_461577 | deoxyribodipyrimidine photo-lyase | 1 | 0.000123123 | 593941 |
| Transcript_448396 | receptor for egg jelly protein | 1 | 0.000127512 | 576647 |
| Transcript_88669 | 40S ribosomal protein S5 | 2 | 0.000129515 | 592668 |
| Transcript_383623 | histidine ammonia-lyase | 1 | 0.000131478 | 583726 |
| Transcript_247937 | EGR1 | 1 | 0.000141967 | 100892559 |
| Transcript_368914 | protein CNPPD1 | 2 | 0.000154939 | 752566 |
| Transcript_428975 | RING finger protein 215 | 4 | 0.000155181 | 753231 |
| Transcript_235182 | programmed cell death protein 2 | 3 | 0.000155446 | 585086 |
| Transcript_280475 | major facilitator superfamily domain-containing protein 6 | 3 | 0.000157081 | 763022 |
| Transcript_459174 | notchless protein homolog 1 | 2 | 0.000161391 | 585438 |
| Transcript_411220 | RING-box protein 1 | 3 | 0.000167486 | 591601 |
| Transcript_88201 | 40S ribosomal protein S4 | 2 | 0.000169885 | 583495 |
| Transcript_339559 | lysozyme 3 | 1 | 0.000170656 | 583337 |
| Transcript_195407 | xylulose kinase\|xylulokinase homolog (H. influenzae) | 1 | 0.000170904 | 589304 |
| Transcript_265718 | sodium-dependent phosphate transport protein 2C | 2 | 0.000171783 | 576137 |
| Transcript_253116 | dynein regulatory complex subunit 6\|F-box/LRR-repeat protein 13 | 1 | 0.000175152 | 577951 |
| Transcript_448836 | tubulointerstitial nephritis antigen-like | 1 | 0.000176097 | 576823 |
| Transcript_215764 | U6 snRNA-associated Sm-like protein LSm2 | 3 | 0.000186826 | 587779 |
| Transcript_443991 | oxidoreductase HTATIP2-like | 3 | 0.000191678 | 592505 |
| Transcript_276623 | histone H4 | 2 | 0.000196637 | 752213 |
| Transcript_360643 | nucleoside diphosphate kinase B | 3 | 0.000199818 | 594617 |
| Transcript_345101 | serine/threonine-protein kinase Nek6 | 1 | 0.00020157 | 100893147 |
| Transcript_445434 | 28S ribosomal protein S17, mitochondrial | 3 | 0.000201605 | 753889 |
| Transcript_352645 | arginine--tRNA ligase, cytoplasmic | 3 | 0.000202136 | 575790 |
| Transcript_199943 | lens fiber major intrinsic protein | 1 | 0.000202136 | 594743 |
| Transcript_253163 | venom phosphodiesterase 2 | 1 | 0.000202136 | 579858 |
| Transcript_278701 | NHP2-like protein 1 | 3 | 0.000206072 | 587294 |
| Transcript_220262 | uncharacterized protein LOC581634 | 2 | 0.000206087 | 581634 |
| Transcript_314700 | uncharacterized protein LOC587761 | 1 | 0.000207742 | 587761 |
| Transcript_402975 | eukaryotic initiation factor 4A-III | 2 | 0.000209348 | 580266 |
| Transcript_284772 | TAF1A | 2 | 0.000213915 | 105440467 |

| | | | | |
|---|---|---|---|---|
| Transcript_361334 | chordin | 1 | 0.000215196 | 580173 |
| Transcript_457700 | lengsin-like | 2 | 0.000216409 | 591380 |
| Transcript_300296 | trimeric intracellular cation channel type 1B.1\|trimeric intracellular cation channel type A | 4 | 0.000227596 | 100890448 |
| Transcript_338151 | histone H3, embryonic | 2 | 0.000227596 | 581030 |
| Transcript_383403 | mediator of RNA polymerase II transcription subunit 18 | 3 | 0.000232301 | 576519 |
| Transcript_456392 | RIIa domain-containing protein 1 | 3 | 0.00023765 | 752632 |
| Transcript_299206 | 39S ribosomal protein L49, mitochondrial | 2 | 0.000241306 | 593029 |
| Transcript_383531 | natural resistance-associated macrophage protein 2 | 2 | 0.000251469 | 576440 |
| Transcript_327352 | macrophage mannose receptor 1 | 1 | 0.000256987 | 100893009 |
| Transcript_172390 | protein SSUH2 homolog | 1 | 0.000257132 | 763611 |
| Transcript_353965 | uncharacterized protein LOC100892584 | 2 | 0.0002581 | 100892584 |
| Transcript_454078 | RNA-binding protein NOB1 | 2 | 0.000262544 | 575793 |
| Transcript_443183 | MYC | 2 | 0.000262997 | 373385 |
| Transcript_329842 | SRA stem-loop-interacting RNA-binding protein, mitochondrial-like | 3 | 0.000265293 | 754241 |
| Transcript_464404 | peroxiredoxin-1 | 2 | 0.000271849 | 590165 |
| Transcript_380053 | cholinesterase 1 | 1 | 0.000275228 | 591159 |
| Transcript_462380 | metalloproteinase inhibitor 3 | 3 | 0.000279283 | 100889149 |
| Transcript_372078 | potassium voltage-gated channel subfamily KQT member 4\|potassium voltage-gated channel su | 3 | 0.000280671 | 579971 |
| Transcript_206791 | uncharacterized protein LOC105442064\|Death effector domain-containing protein | 1 | 0.000288649 | 105442064 |
| Transcript_86447 | glyceraldehyde-3-phosphate dehydrogenase | 2 | 0.000304165 | 578260 |
| Transcript_392219 | neuropeptide Y receptor type 2-like | 3 | 0.000306609 | 100891742 |
| Transcript_327467 | chromosome transmission fidelity protein 8 homolog | 3 | 0.000309435 | 763690 |
| Transcript_308925 | dnaJ homolog subfamily C member 2 | 2 | 0.000310386 | 583525 |
| Transcript_202368 | eukaryotic initiation factor 4a | 2 | 0.000311754 | 575736 |
| Transcript_166464 | translation elongation factor 2 | 2 | 0.000315605 | 592801 |
| Transcript_179487 | uncharacterized protein LOC580673 | 4 | 0.000318961 | 580673 |
| Transcript_134695 | angiotensin-converting enzyme 2 | 1 | 0.000330507 | 580220 |
| Transcript_280569 | TNF receptor-associated factor 6-A | 3 | 0.000332355 | 585060 |
| Transcript_255311 | WD repeat-containing protein 37 | 1 | 0.000344097 | 581583 |
| Transcript_461472 | growth hormone secretagogue receptor type 1\|neuropeptides capa receptor\|growth hormone | 2 | 0.000351928 | 592924 |
| Transcript_345624 | 40S ribosomal protein S9 | 2 | 0.000351928 | 586110 |
| Transcript_230874 | ubiquitin | 2 | 0.000355053 | 584839 |
| Transcript_25798 | ATP synthase F(0) complex subunit C2, mitochondrial\|ATP synthase lipid-binding protein, mitoc | 3 | 0.000360871 | 583818 |
| Transcript_397056 | coatomer subunit epsilon | 3 | 0.000367007 | 582090 |
| Transcript_333839 | G2/mitotic-specific cyclin-B3 | 5 | 0.000367007 | 591240 |
| Transcript_416611 | zinc transporter ZIP12 | 1 | 0.000368191 | 587958 |

| | | | | |
|---|---|---|---|---|
| Transcript_196818 | UDP-glucuronosyltransferase 2C1\|UDP-glucuronosyltransferase 1-2\|UDP-glucuronosyltransfera | 1 | 0.000369716 | 589187 |
| Transcript_280665 | uncharacterized protein LOC100890259 | 1 | 0.000376789 | 100890259 |
| Transcript_249751 | putative Dol-P-Glc:Glc(2)Man(9)GlcNAc(2)-PP-Dol alpha-1,2-glucosyltransferase | 3 | 0.000383409 | 590617 |
| Transcript_483697 | ras-related protein ORAB-1 | 2 | 0.000384483 | 373418 |
| Transcript_323044 | pleiotropic regulator 1 | 2 | 0.000388456 | 756406 |
| Transcript_189980 | uncharacterized protein LOC100890349\|L-idonate 5-dehydrogenase | 1 | 0.000406171 | 100890349 |
| Transcript_301658 | nucleolar protein 10 | 2 | 0.000408603 | 590723 |
| Transcript_459372 | fibrillin-3\|bypass of stop codon protein 1 | 4 | 0.000410913 | 591161 |
| Transcript_199327 | delta-aminolevulinic acid dehydratase | 2 | 0.00041335 | 577020 |
| Transcript_183069 | glutamate receptor 1 | 2 | 0.000415045 | 587837 |
| Transcript_192833 | 5-oxoprolinase\|5-oxoprolinase (ATP-hydrolysing) | 1 | 0.000427737 | 589767 |
| Transcript_373239 | ras-related protein Rap-2c | 2 | 0.000440545 | 753090 |
| Transcript_386406 | WD repeat-containing protein 43 | 2 | 0.000451267 | 592550 |
| Transcript_238124 | protein MAK16 homolog A | 2 | 0.000460165 | 586606 |
| Transcript_233725 | uncharacterized protein C2orf50 | 1 | 0.000494308 | 578348 |
| Transcript_300378 | CDP-diacylglycerol--glycerol-3-phosphate 3-phosphatidyltransferase, mitochondrial | 3 | 0.000520562 | 581340 |
| Transcript_421581 | pre-rRNA-processing protein TSR2 homolog | 2 | 0.000525637 | 592405 |
| Transcript_236174 | 40S ribosomal protein S3 | 2 | 0.000532436 | 593744 |
| Transcript_249550 | sorbitol dehydrogenase | 1 | 0.000534208 | 585570 |
| Transcript_234548 | angiopoietin-4-like | 1 | 0.0005769 | 100891514 |
| Transcript_128686 | pre-rRNA 2'-O-ribose RNA methyltransferase FTSJ3\|pre-rRNA processing protein FTSJ3 | 3 | 0.000590448 | 582089 |
| Transcript_333875 | uncharacterized protein LOC578017 | 5 | 0.000597208 | 578017 |
| Transcript_208042 | ST8 alpha-N-acetyl-neuraminide alpha-2,8-sialyltransferase 4 | 1 | 0.000631501 | 586329 |
| Transcript_257273 | methylsterol monooxygenase 1 | 1 | 0.000641983 | 100891893 |
| Transcript_308851 | alcohol dehydrogenase class-3 | 1 | 0.000649203 | 579220 |
| Transcript_385086 | pre-piRNA 3'-exonuclease trimmer\|poly(A)-specific ribonuclease PARN-like domain-containing | 2 | 0.000665071 | 100891016 |
| Transcript_408947 | phosphopantothenoylcysteine decarboxylase | 3 | 0.000674726 | 584017 |
| Transcript_273618 | SHC SH2 domain-binding protein 1 | 4 | 0.000680453 | 575406 |
| Transcript_418431 | ketimine reductase mu-crystallin | 1 | 0.000684468 | 579068 |
| Transcript_214978 | basic leucine zipper and W2 domain-containing protein 1 | 2 | 0.000692153 | 587479 |
| Transcript_432419 | beta-lactamase domain-containing protein 2 | 1 | 0.000718546 | 575771 |
| Transcript_196327 | creatine kinase, flagellar | 5 | 0.00071996 | 580751 |
| Transcript_221788 | NADPH oxidase | 1 | 0.000721382 | 757329 |
| Transcript_329183 | E3 ubiquitin-protein ligase UHRF1\|histone-lysine N-methyltransferase, H3 lysine-9 specific SUV | 1 | 0.000725484 | 756912 |
| Transcript_346536 | cAMP-dependent protein kinase catalytic subunit 1\|catalytic subunit of cAMP-dependent histor | 2 | 0.000751818 | 589947 |
| Transcript_366071 | N-acetylgalactosamine-6-sulfatase-like | 1 | 0.000766197 | 579130 |

| | | | | |
|---|---|---|---|---|
| Transcript_235911 | phospholipase B1, membrane-associated | 1 | 0.000775871 | 100890912 |
| Transcript_403482 | aldo-keto reductase family 1 member A1\|alcohol dehydrogenase | 4 | 0.000784233 | 576680 |
| Transcript_334842 | uncharacterized protein LOC580196 | 2 | 0.000786461 | 580196 |
| Transcript_235072 | uncharacterized protein LOC579112 | 4 | 0.000788949 | 579112 |
| Transcript_82280 | calmodulin | 2 | 0.000790274 | 575365 |
| Transcript_424005 | soma ferritin | 3 | 0.000806 | 591499 |
| Transcript_419016 | dnaJ homolog subfamily C member 5\|cysteine string protein | 2 | 0.000807136 | 577697 |
| Transcript_12298 | 40S ribosomal protein S23 | 2 | 0.000841834 | 590530 |
| Transcript_368000 | syntenin-1 | 2 | 0.000847174 | 578747 |
| Transcript_448298 | protein Wnt-7b | 1 | 0.000850909 | 581981 |
| Transcript_374203 | protein Asterix | 3 | 0.000866756 | 592477 |
| Transcript_121057 | V-type proton ATPase 16 kDa proteolipid subunit | 2 | 0.000903696 | 593221 |
| Transcript_276903 | ERI1 exoribonuclease 2 | 1 | 0.00096432 | 585927 |
| Transcript_388827 | ADP/ATP translocase 3 | 2 | 0.000969427 | 575225 |
| Transcript_240937 | uracil phosphoribosyltransferase homolog | 2 | 0.000983598 | 575128 |
| Transcript_444850 | probable RNA-binding protein EIF1AD\|putative RNA-binding protein EIF1AD | 3 | 0.000991161 | 578771 |
| Transcript_427937 | DNA repair protein XRCC3 | 3 | 0.00101429 | 763443 |
| Transcript_286704 | extended synaptotagmin-2 | 5 | 0.001044619 | 579674 |
| Transcript_263515 | polycystic kidney disease protein 1-like 2 | 1 | 0.001052967 | 579144 |
| Transcript_357605 | glutathione S-transferase P | 3 | 0.001055612 | 580662 |
| Transcript_438353 | EMX1 | 4 | 0.001088813 | 577702 |
| Transcript_264220 | ADP-ribosylation factor 1\|ADP-ribosylation factor-like | 3 | 0.001110619 | 586151 |
| Transcript_208170 | uncharacterized protein LOC105441928 | 1 | 0.001110863 | 105441928 |
| Transcript_383458 | probable 60S ribosomal protein L37-A | 2 | 0.001114113 | 591582 |
| Transcript_51640 | RCC1 and BTB domain-containing protein 1\|ultraviolet-B receptor UVR8\|probable E3 ubiquitin- | 4 | 0.001129851 | 577290 |
| Transcript_305797 | WD repeat-containing protein 31 | 1 | 0.001163089 | 585021 |
| Transcript_240632 | calpain small subunit 1 | 4 | 0.001196235 | 100889224 |
| Transcript_171991 | protein cornichon homolog 1 | 3 | 0.00119957 | 582532 |
| Transcript_398758 | serine/threonine-protein phosphatase PGAM5, mitochondrial | 2 | 0.001214001 | 588383 |
| Transcript_353555 | caspase-6 | 1 | 0.001318326 | 584221 |
| Transcript_188732 | tetratricopeptide repeat protein 32 | 2 | 0.001350098 | 588488 |
| Transcript_264574 | conserved oligomeric Golgi complex subunit 2\|component of oligomeric golgi complex 2 | 3 | 0.001377664 | 589392 |
| Transcript_392565 | methionyl-tRNA formyltransferase, mitochondrial | 2 | 0.001379209 | 586344 |
| Transcript_235778 | ST8 alpha-N-acetyl-neuraminide alpha-2,8-sialyltransferase 7\|TM2 domain-containing protein | 1 | 0.001456409 | 583563 |
| Transcript_463187 | mitochondrial import inner membrane translocase subunit Tim10 B | 2 | 0.001462773 | 574660 |
| Transcript_359454 | dynein light chain LC6, flagellar outer arm | 4 | 0.001472937 | 591043 |

| | | | | |
|---|---|---|---|---|
| Transcript_402965 | PRELI domain containing protein 3B\|protein slowmo homolog 2 | 2 | 0.001481224 | 100889391 |
| Transcript_434653 | pre-rRNA-processing protein TSR1 homolog\|TSR1, 20S rRNA accumulation, homolog | 2 | 0.001486958 | 594000 |
| Transcript_323407 | uncharacterized protein LOC763845 | 1 | 0.001498958 | 763845 |
| Transcript_456790 | acyloxyacyl hydrolase | 1 | 0.001508263 | 592185 |
| Transcript_309275 | uncharacterized protein LOC100889927 | 1 | 0.001508866 | 100889927 |
| Transcript_350913 | lysozyme | 4 | 0.001579517 | 587263 |
| Transcript_291331 | matrix metalloproteinase-24 | 2 | 0.001579939 | 577128 |
| Transcript_228378 | ADP-ribosylation factor-like protein 6 | 2 | 0.001582577 | 591507 |
| Transcript_383671 | anoctamin-8 | 1 | 0.001595851 | 588981 |
| Transcript_456874 | coiled-coil domain-containing protein 51 | 2 | 0.00167281 | 585678 |
| Transcript_328102 | protein Aster-B\|GRAM domain-containing protein 1B | 4 | 0.001674503 | 754516 |
| Transcript_173387 | HNF4G | 1 | 0.001696184 | 574894 |
| Transcript_453741 | uncharacterized protein LOC100892807 | 3 | 0.001833911 | 100892807 |
| Transcript_396224 | sodium/glucose cotransporter 4 | 1 | 0.001869933 | 593500 |
| Transcript_184289 | F-box/LRR-repeat protein 7 | 1 | 0.001893942 | 580713 |
| Transcript_435724 | coiled-coil domain-containing protein 151 | 5 | 0.001915415 | 100893724 |
| Transcript_278188 | leucine-rich repeat-containing protein 15 | 1 | 0.001944018 | 100893380 |
| Transcript_228502 | S-crystallin SL11 | 1 | 0.001945235 | 586571 |
| Transcript_430758 | uncharacterized protein LOC100888279 | 1 | 0.001948734 | 100888279 |
| Transcript_415734 | peptidase M20 domain-containing protein 2 | 4 | 0.001958887 | 584225 |
| Transcript_292151 | uncharacterized protein LOC105437474 | 1 | 0.002019107 | 105437474 |
| Transcript_359398 | cytochrome P450 26A1\|cytochrome P450 26B1 | 1 | 0.002045033 | 763716 |
| Transcript_417667 | ER membrane protein complex subunit 8\|ER membrane protein complex subunit 8 pseudogene | 3 | 0.002058001 | 582477 |
| Transcript_363862 | RNA-binding protein cabeza | 1 | 0.00208612 | 105447104 |
| Transcript_257591 | fibrillin-2 | 1 | 0.002112952 | 581033 |
| Transcript_224730 | cytochrome c oxidase subunit 5B, mitochondrial | 3 | 0.002126058 | 581274 |
| Transcript_456276 | heparan sulfate glucosamine 3-O-sulfotransferase 1 | 1 | 0.002149764 | 764585 |
| Transcript_290553 | isoamyl acetate-hydrolyzing esterase 1 homolog | 4 | 0.002263181 | 591412 |
| Transcript_324078 | protein PBDC1\|UPF0368 protein Cxorf26-like | 2 | 0.00230034 | 577502 |
| Transcript_236625 | CDX1 | 1 | 0.002327014 | 584191 |
| Transcript_309101 | uncharacterized protein LOC105443655 | 3 | 0.002403175 | 105443655 |
| Transcript_350196 | ATP-binding cassette sub-family F member 2\|ATP-binding cassette, sub-family F (GCN20), mem | 2 | 0.002443557 | 581859 |
| Transcript_204615 | U6 snRNA-associated Sm-like protein LSm7 | 3 | 0.002487358 | 594039 |
| Transcript_284394 | EF-hand calcium-binding domain-containing protein 12 | 1 | 0.002516391 | 763537 |
| Transcript_229121 | LOW QUALITY PROTEIN: MMP37-like protein, mitochondrial\|MMP37-like protein, mitochondria | 2 | 0.002561667 | 575960 |
| Transcript_310649 | receptor-type guanylate cyclase gcy-8\|insulin-like growth factor 1 receptor | 1 | 0.002569395 | 591965 |

| | | | | |
|---|---|---|---|---|
| Transcript_239164 | 3-hydroxyanthranilate 3,4-dioxygenase | 1 | 0.002593445 | 582154 |
| Transcript_234886 | multidrug resistance-associated protein 4\|ATP-binding cassette, sub-family C (CFTR/MRP), mem | 5 | 0.002593978 | 591982 |
| Transcript_326960 | uncharacterized protein LOC100891910 | 1 | 0.002627634 | 100891910 |
| Transcript_186728 | methylglutaconyl-CoA hydratase, mitochondrial | 1 | 0.002651092 | 577505 |
| Transcript_189831 | L-gulonolactone oxidase | 5 | 0.002653064 | 764246 |
| Transcript_214947 | calcium load-activated calcium channel\|transmembrane and coiled-coil domain-containing prot | 3 | 0.002669245 | 100892898 |
| Transcript_444322 | tubulin beta-4B chain | 5 | 0.002730604 | 593362 |
| Transcript_309473 | autocrine proliferation repressor protein A | 1 | 0.002840025 | 100889791 |
| Transcript_254514 | 14-3-3 family protein artA\|14-3-3 protein 3 | 2 | 0.002856679 | 581376 |
| Transcript_187684 | bcl-2 homologous antagonist/killer | 2 | 0.002938261 | 588418 |
| Transcript_54760 | testis-specific serine/threonine-protein kinase 4-like | 5 | 0.00297144 | 575157 |
| Transcript_304049 | uncharacterized aarF domain-containing protein kinase 1\|putative aarF domain-containing prot | 2 | 0.003065219 | 574821 |
| Transcript_363577 | uncharacterized protein LOC105438509 | 3 | 0.003084148 | 105438509 |
| Transcript_340557 | small nuclear ribonucleoprotein polypeptide E-like | 3 | 0.003114499 | 580958 |
| Transcript_182361 | rho GTPase-activating protein 25 | 2 | 0.00313536 | 582939 |
| Transcript_204566 | ruvB-like 1 | 2 | 0.003144896 | 577255 |
| Transcript_354492 | presenilins-associated rhomboid-like protein, mitochondrial | 2 | 0.003374583 | 576709 |
| Transcript_374584 | 28S ribosomal protein S34, mitochondrial | 2 | 0.003395751 | 753857 |
| Transcript_208329 | univin | 3 | 0.003400545 | 373488 |
| Transcript_243976 | S-methylmethionine--homocysteine S-methyltransferase BHMT2-like | 1 | 0.00340357 | 582542 |
| Transcript_343639 | glycine-rich cell wall structural protein 1\|glycine-rich protein DOT1 | 2 | 0.003455777 | 764696 |
| Transcript_236992 | cyclic nucleotide-gated cation channel alpha-3 | 3 | 0.003662216 | 576815 |
| Transcript_326935 | deleted in malignant brain tumors 1 protein-like\|neurotrypsin-like | 1 | 0.003695293 | 762897 |
| Transcript_445886 | neurotrypsin | 1 | 0.003720023 | 578211 |
| Transcript_270797 | uncharacterized protein LOC575472 | 1 | 0.003755055 | 575472 |
| Transcript_456555 | scavenger receptor cysteine-rich protein type 12 | 1 | 0.003772405 | 373431 |
| Transcript_448686 | fibrinogen C domain-containing protein 1-like | 1 | 0.003887687 | 105440826 |
| Transcript_450213 | thioredoxin domain-containing protein 5 | 2 | 0.004022943 | 764720 |
| Transcript_243778 | UDP-glucuronosyltransferase 2C1 | 1 | 0.004132616 | 592958 |
| Transcript_179967 | condensin complex subunit 3-like | 1 | 0.004145511 | 105447056 |
| Transcript_187613 | solute carrier family 28 member 3 | 1 | 0.004382769 | 574602 |
| Transcript_280373 | adenosylhomocysteinase | 1 | 0.004452691 | 574714 |
| Transcript_317804 | uncharacterized protein LOC100891690 | 3 | 0.004527764 | 100891690 |
| Transcript_438092 | ATP-binding cassette, sub-family G (WHITE), member 2 | 1 | 0.004755746 | 578540 |
| Transcript_356568 | cystathionine gamma-lyase\|putative cystathionine gamma-lyase 2 | 2 | 0.0051019 | 762952 |
| Transcript_182233 | nose resistant to fluoxetine protein 6 | 1 | 0.005139831 | 587932 |

| | | | | |
|---|---|---|---|---|
| Transcript_438569 | acid-sensing ion channel 1A | 1 | 0.005184244 | 575477 |
| Transcript_314048 | transmembrane 6 superfamily member 1 | 4 | 0.005222331 | 588706 |
| Transcript_376909 | 5'-nucleotidase | 1 | 0.005354264 | 753743 |
| Transcript_368980 | fumarylacetoacetate hydrolase domain-containing protein 2 | 5 | 0.005436549 | 763463 |
| Transcript_271521 | fos-related antigen 1 | 1 | 0.0054842 | 579838 |
| Transcript_28806 | queuine tRNA-ribosyltransferase catalytic subunit 1-like | 2 | 0.00553486 | 579235 |
| Transcript_319918 | uncharacterized protein LOC100888641 | 1 | 0.005688136 | 100888641 |
| Transcript_395198 | synaptotagmin-14\|synaptotagmin-16 | 5 | 0.005940282 | 577619 |
| Transcript_330815 | fibronectin-like | 3 | 0.005999686 | 105447587 |
| Transcript_271208 | glycine-rich cell wall structural protein-like | 4 | 0.006002727 | 105447344 |
| Transcript_200340 | uncharacterized protein YER152C | 4 | 0.006272248 | 753534 |
| Transcript_269278 | HES4 | 1 | 0.006281806 | 592057 |
| Transcript_443316 | balbiani ring protein 3\|prestalk protein\|whey acidic protein-like | 1 | 0.006407029 | 755393 |
| Transcript_74981 | DNA-directed RNA polymerases I, II, and III subunit RPABC3 | 2 | 0.006545429 | 591688 |
| Transcript_308440 | dimethylaniline monooxygenase [N-oxide-forming] 5 | 1 | 0.006648833 | 580981 |
| Transcript_190979 | leucine-rich repeat-containing protein 69 | 1 | 0.006656512 | 579382 |
| Transcript_326137 | LOW QUALITY PROTEIN: tRNA (adenine(37)-N6)-methyltransferase\|nef-associated protein 1 | 3 | 0.006798408 | 577363 |
| Transcript_462766 | sodium-dependent multivitamin transporter | 2 | 0.006889503 | 578787 |
| Transcript_234973 | nitric oxide-associated protein 1 | 2 | 0.006984558 | 576629 |
| Transcript_437119 | amassin-3 | 3 | 0.007112091 | 580528 |
| Transcript_201140 | ficolin-1-like | 1 | 0.007272042 | 587736 |
| Transcript_299780 | rRNA methyltransferase 2, mitochondrial | 3 | 0.00755268 | 582250 |
| Transcript_315513 | sushi domain-containing protein 2 | 1 | 0.007619095 | 580458 |
| Transcript_443412 | E3 ubiquitin-protein ligase MARCH8 | 3 | 0.007636877 | 579071 |
| Transcript_282963 | acetoacetyl-CoA synthetase | 3 | 0.007722755 | 590598 |
| Transcript_426847 | retinol dehydrogenase 8 | 2 | 0.007781435 | 574716 |
| Transcript_341560 | cytochrome b-c1 complex subunit 6, mitochondrial | 3 | 0.007857858 | 590393 |
| Transcript_231242 | ATP-citrate synthase | 1 | 0.008053833 | 594785 |
| Transcript_392370 | methyltransferase-like protein 22 | 1 | 0.008061862 | 100894107 |
| Transcript_50022 | 60S ribosomal protein L30 | 2 | 0.00815734 | 577852 |
| Transcript_219767 | lactase-phlorizin hydrolase | 1 | 0.008398588 | 581938 |
| Transcript_268430 | ZNF260 | 4 | 0.008456303 | 100892543 |
| Transcript_198022 | P-selectin | 2 | 0.008474676 | 105437801 |
| Transcript_433401 | proteasome assembly chaperone 3 | 2 | 0.008654302 | 100892637 |
| Transcript_230273 | dynein heavy chain 8, axonemal\|dynein, axonemal, heavy chain 8 | 1 | 0.008769573 | 763294 |
| Transcript_402513 | acidic phospholipase A2\|basic phospholipase A2 nigroxin A | 1 | 0.00888625 | 757121 |

| | | | | |
|---|---|---|---|---|
| Transcript_326493 | sodium-independent sulfate anion transporter | 1 | 0.008989487 | 574650 |
| Transcript_388333 | ribonuclease H2 subunit C | 3 | 0.009211094 | 752797 |
| Transcript_462732 | nesprin-1 | 5 | 0.009332795 | 105437706 |
| Transcript_193119 | titin\|muscle M-line assembly protein unc-89\|putative titin-like | 3 | 0.00939858 | 590007 |
| Transcript_210888 | diamine acetyltransferase 2 | 3 | 0.009551632 | 100893060 |
| Transcript_440549 | uncharacterized protein LOC577038 | 1 | 0.010145783 | 577038 |
| Transcript_48454 | 40S ribosomal protein S16 | 2 | 0.010706301 | 577040 |
| Transcript_459460 | THAP6 | 3 | 0.010762146 | 577120 |
| Transcript_336905 | uncharacterized protein LOC100888517 | 1 | 0.010835345 | 100888517 |
| Transcript_437026 | uncharacterized protein K02A2.6-like | 5 | 0.010979249 | 105444440 |
| Transcript_449725 | calcineurin B homologous protein 1 | 2 | 0.010979249 | 576455 |
| Transcript_354357 | APOBEC1 complementation factor | 1 | 0.011175702 | 575684 |
| Transcript_258396 | uncharacterized protein LOC588928 | 1 | 0.011267375 | 588928 |
| Transcript_453256 | WD repeat-containing protein 86 | 3 | 0.01200482 | 100888294 |
| Transcript_226806 | cytochrome P450 3A24 | 4 | 0.012022093 | 576854 |
| Transcript_413396 | 3-ketoacyl-CoA thiolase A, peroxisomal | 1 | 0.012424432 | 587557 |
| Transcript_280723 | scavenger receptor cysteine-rich protein | 1 | 0.012532041 | 373211 |
| Transcript_61404 | dynein heavy chain 5, axonemal | 1 | 0.012843572 | 577371 |
| Transcript_13256 | protein disulfide-isomerase A5 | 5 | 0.013269666 | 576717 |
| Transcript_292279 | dnaJ homolog subfamily C member 28 | 1 | 0.013626876 | 587875 |
| Transcript_256585 | microfibril-associated glycoprotein 4 | 1 | 0.013830912 | 582445 |
| Transcript_453252 | N-acetyltransferase 9-like protein | 2 | 0.014015317 | 586164 |
| Transcript_374389 | alpha-(1,3)-fucosyltransferase 6 | 4 | 0.014272136 | 579774 |
| Transcript_265355 | protein boule-like\|bromodomain-containing protein 4-like | 1 | 0.014681263 | 100892519 |
| Transcript_297498 | EH domain-containing protein 1\|EH domain-containing protein 3-like | 2 | 0.014744428 | 587455 |
| Transcript_204631 | tyrosine-protein kinase receptor Tie-2-like | 1 | 0.014745656 | 105439696 |
| Transcript_352756 | protein TFG | 3 | 0.014753533 | 594769 |
| Transcript_400118 | uncharacterized protein LOC100893690 | 1 | 0.015012477 | 100893690 |
| Transcript_451319 | 40S ribosomal protein S12 | 2 | 0.015117118 | 590740 |
| Transcript_440408 | N-fatty-acyl-amino acid synthase/hydrolase PM20D1\|probable carboxypeptidase PM20D1\|puta | 4 | 0.015510861 | 579513 |
| Transcript_285315 | N-alpha-acetyltransferase 40 | 2 | 0.015578148 | 577667 |
| Transcript_361820 | L-threonine 3-dehydrogenase, mitochondrial\|inactive L-threonine 3-dehydrogenase, mitochond | 2 | 0.015611387 | 584440 |
| Transcript_189682 | peroxisomal carnitine O-octanoyltransferase | 1 | 0.015664898 | 585404 |
| Transcript_214210 | late histone H2A.L3 | 3 | 0.015709704 | 373348 |
| Transcript_367907 | uncharacterized protein LOC586472\|Small GTPase superfamily domain containing protein | 1 | 0.015758585 | 586472 |
| Transcript_393644 | SAFB-like transcription modulator | 2 | 0.01587077 | 585734 |

| | | | | |
|---|---|---|---|---|
| Transcript_257884 | uncharacterized protein LOC583434\|urease | 1 | 0.016315729 | 583434 |
| Transcript_195199 | arylsulfatase | 1 | 0.016536437 | 582391 |
| Transcript_257979 | echinoderm microtubule-associated protein-like 4\|uncharacterized protein LOC581284 | 1 | 0.016740806 | 581284 |
| Transcript_321307 | mitochondrial fission regulator 2\|mitochondrial fission regulator 1 | 3 | 0.016869408 | 764790 |
| Transcript_240999 | cyclin-dependent kinases regulatory subunit | 1 | 0.017248255 | 100891492 |
| Transcript_2061 | mitochondrial ATP synthase alpha subunit precursor | 2 | 0.018413908 | 373382 |
| Transcript_355725 | protein disulfide-isomerase A3 | 2 | 0.018926929 | 577673 |
| Transcript_258178 | zinc finger protein 474-like | 5 | 0.01959868 | 100888783 |
| Transcript_225983 | histamine N-methyltransferase | 4 | 0.019730461 | 582038 |
| Transcript_388758 | uncharacterized protein LOC591506\|UPF0394 inner membrane protein yeeE-like | 1 | 0.020558406 | 591506 |
| Transcript_252605 | cation-dependent mannose-6-phosphate receptor-like | 1 | 0.020779714 | 105438545 |
| Transcript_307974 | malignant fibrous histiocytoma-amplified sequence 1 homolog | 3 | 0.020863101 | 100888805 |
| Transcript_325168 | zonadhesin-like | 1 | 0.021381996 | 105438765 |
| Transcript_269460 | probable D-lactate dehydrogenase, mitochondrial\|lactate dehydrogenase D | 4 | 0.021486102 | 591812 |
| Transcript_256749 | peptidyl-prolyl cis-trans isomerase G\|serine/arginine repetitive matrix protein 2-like | 1 | 0.021654179 | 100890356 |
| Transcript_258900 | KRP170 | 4 | 0.021974774 | 373241 |
| Transcript_386762 | uncharacterized protein LOC100892326 | 4 | 0.022071669 | 100892326 |
| Transcript_392136 | somatostatin receptor type 2 | 1 | 0.022255228 | 100889243 |
| Transcript_405707 | ADP-ribosyl cyclase\|ADP-ribosyl cyclase beta | 1 | 0.022643114 | 580868 |
| Transcript_280991 | phytanoyl-CoA dioxygenase domain-containing protein 1 | 1 | 0.022977014 | 753147 |
| Transcript_314947 | microfibril-associated glycoprotein 4-like | 3 | 0.022996409 | 581168 |
| Transcript_396061 | putative ISG12 protein | 1 | 0.023872748 | 403119 |
| Transcript_412518 | ATP-dependent (S)-NAD(P)H-hydrate dehydratase | 1 | 0.025975248 | 586141 |
| Transcript_246157 | LOW QUALITY PROTEIN: tubulin beta chain | 5 | 0.026130789 | 594231 |
| Transcript_351435 | medium-chain acyl-CoA ligase ACSF2, mitochondrial-like\|acyl-CoA synthetase family member 2, | 1 | 0.02642635 | 593582 |
| Transcript_200945 | cathepsin L1 | 3 | 0.027167837 | 575203 |
| Transcript_308628 | protein PRY1\|golgi-associated plant pathogenesis-related protein 1 | 1 | 0.027167837 | 100893059 |
| Transcript_398823 | uncharacterized protein LOC589927\|CUB domain-containing protein | 1 | 0.027751362 | 589927 |
| Transcript_343503 | LOW QUALITY PROTEIN: very early blastula protein 4\|very early blastula protein 4 | 5 | 0.02946305 | 373489 |
| Transcript_194733 | uncharacterized protein LOC579958 | 1 | 0.029757117 | 579958 |
| Transcript_241860 | denticleless protein homolog | 4 | 0.030087247 | 584715 |
| Transcript_265286 | tumor suppressor candidate 2 | 2 | 0.030240058 | 594794 |
| Transcript_254071 | actin, cytoskeletal 3B\|actin, cytoskeletal 3 | 3 | 0.03034686 | 100890099 |
| Transcript_459535 | transmembrane protein 187 | 4 | 0.031438231 | 100887935 |
| Transcript_193614 | somatostatin receptor type 5 | 2 | 0.031787754 | 105444813 |
| Transcript_386066 | zinc finger protein 236 | 2 | 0.031981685 | 100890022 |

| | | | | |
|---|---|---|---|---|
| Transcript_407578 | solute carrier family 23 member 1 | 1 | 0.032288234 | 581718 |
| Transcript_373383 | uncharacterized protein LOC581986\|proteinase T | 1 | 0.032789184 | 581986 |
| Transcript_284740 | uncharacterized protein C16orf96 homolog\|uncharacterized protein C16orf96\|uncharacterized | 5 | 0.033005953 | 755388 |
| Transcript_57336 | profilin\|spCoel1 | 3 | 0.034328683 | 373409 |
| Transcript_299519 | uncharacterized protein LOC105447398 | 2 | 0.034769137 | 105447398 |
| Transcript_289174 | annexin A7 | 3 | 0.035301747 | 764615 |
| Transcript_453447 | NAD kinase | 3 | 0.035404269 | 584340 |
| Transcript_346874 | alpha-tocopherol transfer protein-like | 3 | 0.035599419 | 575290 |
| Transcript_324882 | uncharacterized protein LOC105440703 | 3 | 0.035609898 | 105440703 |
| Transcript_375307 | uncharacterized protein LOC100888794 | 1 | 0.037345332 | 100888794 |
| Transcript_285525 | uncharacterized protein LOC100891603 | 3 | 0.037526216 | 100891603 |
| Transcript_315280 | meiosis 1 arrest protein | 1 | 0.037581022 | 100893539 |
| Transcript_54635 | uncharacterized protein LOC100891675 | 3 | 0.03765683 | 100891675 |
| Transcript_359165 | calponin homology domain-containing protein DDB_G0272472\|intracellular protein transport p | 1 | 0.037852399 | 100890558 |
| Transcript_281352 | centrosomal protein of 290 kDa-like | 2 | 0.038176421 | 105442844 |
| Transcript_117530 | D(2) dopamine receptor A-like | 1 | 0.038798225 | 105439410 |
| Transcript_82652 | mitochondrial ribosome-associated GTPase 1 | 4 | 0.039178574 | 588789 |
| Transcript_427415 | paraneoplastic antigen Ma3-like | 2 | 0.039625395 | 105447116 |
| Transcript_345735 | DLX5 | 3 | 0.04058574 | 593496 |
| Transcript_218552 | cytosolic beta-glucosidase | 1 | 0.04098007 | 589416 |
| Transcript_322346 | triple QxxK/R motif-containing protein-like\|acyl-CoA synthetase family member 3, mitochondri | 4 | 0.041155658 | 105440874 |
| Transcript_300712 | protein DDI1 homolog 2 | 2 | 0.04169834 | 590177 |
| Transcript_67216 | cGMP-dependent protein kinase 1 | 2 | 0.042140028 | 589462 |
| Transcript_390895 | vesicular inhibitory amino acid transporter-like | 1 | 0.043654839 | 755450 |
| Transcript_366842 | bindin | 5 | 0.044607927 | 373276 |
| Transcript_207308 | uncharacterized protein LOC587864 | 1 | 0.045911275 | 587864 |
| Transcript_177363 | LOW QUALITY PROTEIN: aromatic-L-amino-acid decarboxylase | 1 | 0.04611209 | 577767 |
| Transcript_371808 | dexamethasone-induced Ras-related protein 1 | 3 | 0.04725451 | 578089 |
| Transcript_206242 | uncharacterized protein LOC756858\|Solute-binding protein family 3/N-terminal domain of Mlt | 1 | 0.047665727 | 756858 |
| Transcript_64977 | 40S ribosomal protein S15 | 3 | 0.048248049 | 574998 |
| Transcript_364074 | LOW QUALITY PROTEIN: zinc finger protein 708-like | 2 | 0.048728883 | 593726 |

Table S19. Functional analysis of coexpressed clusters

| Cluster | Category | Term | Count | % | PValue | Genes | List Total | Pop Hits | Pop Total | Fold Enrichment | Bonferroni | Benjamini | FDR |
|---|---|---|---|---|---|---|---|---|---|---|---|---|---|
| 1 | Biological Process | GO:0044699~single-organism process | 23 | 7.565789474 | 2.70E-04 | 591943, 591159, 592958, 755450, 100891492, 577570, 757121, 581938, 105439410, 577952, 580868, | 28 | 81 | 162 | 1.642857143 | 0.086471814 | 0.090428842 | 0.090428842 |
| | Biological Process | GO:0044763~single-organism cellular process | 20 | 6.578947368 | 0.006625553 | 591943, 591159, 592958, 755450, 100891492, 577570, 757121, 105439410, 577952, 580868, 1008892 | 28 | 76 | 162 | 1.522556391 | 0.892142638 | 1 | 1 |
| | Biological Process | GO:0005975~carbohydrate metabolic process | 4 | 1.315789474 | 0.033087446 | 586329, 586832, 581938, 583563 | 28 | 5 | 162 | 4.628571429 | 0.999987273 | 1 | 1 |
| | Biological Process | GO:0044710~single-organism metabolic process | 9 | 2.960526316 | 0.050405274 | 100891893, 592958, 586329, 574714, 577570, 757121, 581938, 583563, 100891910 | 28 | 27 | 162 | 1.928571429 | 0.99999997 | 1 | 1 |
| | Biological Process | GO:0044707~single-multicellular organism process | 4 | 1.315789474 | 0.058741299 | 591159, 373360, 580173, 105439410 | 28 | 6 | 162 | 3.857142857 | 0.999999998 | 1 | 1 |
| | Biological Process | GO:0098609~cell-cell adhesion | 3 | 0.986842105 | 0.072334369 | 591943, 591159, 105440441 | 28 | 3 | 162 | 5.785714286 | 1 | 1 | 1 |
| | Biological Process | GO:0006082~organic acid metabolic process | 5 | 1.644736842 | 0.088248913 | 100891893, 592958, 574714, 577570, 100891910 | 28 | 11 | 162 | 2.62987013 | 1 | 1 | 1 |
| | Biological Process | GO:0032501~multicellular organismal process | 4 | 1.315789474 | 0.091301958 | 591159, 373360, 580173, 105439410 | 28 | 7 | 162 | 3.306122449 | 1 | 1 | 1 |
| | Biological Process | GO:0022610~biological adhesion | 4 | 1.315789474 | 0.091301958 | 591943, 591159, 576823, 105440441 | 28 | 7 | 162 | 3.306122449 | 1 | 1 | 1 |
| | Biological Process | GO:0007155~cell adhesion | 4 | 1.315789474 | 0.091301958 | 591943, 591159, 576823, 105440441 | 28 | 7 | 162 | 3.306122449 | 1 | 1 | 1 |
| | Cellular Component | GO:0016020~membrane | 28 | 9.210526316 | 3.25E-08 | 591159, 373211, 373431, 755450, 585092, 578540, 100893380, 105439410, 592585, 105437474, 5769 | 38 | 61 | 183 | 2.210526316 | 2.18E-06 | 1.25E-06 | 1.21E-06 |
| | Cellular Component | GO:0031224~intrinsic component of membrane | 25 | 8.223684211 | 3.73E-08 | 591159, 755450, 578540, 100893380, 105439410, 592585, 105437474, 576910, 582219, 586832, 1054 | 38 | 49 | 183 | 2.457035446 | 2.50E-06 | 1.25E-06 | 1.21E-06 |
| | Cellular Component | GO:0016021~integral component of membrane | 24 | 7.894736842 | 1.79E-07 | 591943, 586672, 591159, 592958, 755450, 578540, 100893380, 100888815, 105439410, 105437474, 5 | 38 | 48 | 183 | 2.407894737 | 1.20E-05 | 3.99E-06 | 3.87E-06 |
| | Cellular Component | GO:0044425~membrane part | 25 | 8.223684211 | 5.07E-07 | 591159, 755450, 578540, 100893380, 105439410, 592585, 105437474, 576910, 582219, 586832, 1054 | 38 | 54 | 183 | 2.229532164 | 3.40E-05 | 8.49E-06 | 8.24E-06 |
| | Cellular Component | GO:0071944~cell periphery | 6 | 1.973684211 | 0.017738484 | 580868, 100889243, 591159, 582219, 105440441, 589852 | 38 | 9 | 183 | 3.210526316 | 0.698548349 | 0.198079733 | 0.192166905 |
| | Cellular Component | GO:0005886~plasma membrane | 6 | 1.973684211 | 0.017738484 | 580868, 100889243, 591159, 582219, 105440441, 589852 | 38 | 9 | 183 | 3.210526316 | 0.698548349 | 0.198079733 | 0.192166905 |
| | Molecular Function | GO:0060089~molecular transducer activity | 8 | 2.631578947 | 0.002391091 | 100889243, 591159, 576823, 373431, 373211, 585092, 105439410, 100888815 | 46 | 10 | 183 | 3.182608696 | 0.318209368 | 0.191287272 | 0.191287272 |
| | Molecular Function | GO:0004872~receptor activity | 8 | 2.631578947 | 0.002391091 | 100889243, 591159, 576823, 373431, 373211, 585092, 105439410, 100888815 | 46 | 10 | 183 | 3.182608696 | 0.318209368 | 0.191287272 | 0.191287272 |
| | Molecular Function | GO:0038024~cargo receptor activity | 4 | 1.315789474 | 0.046606505 | 576823, 373431, 373211, 585092 | 46 | 4 | 183 | 3.97826087 | 0.999517442 | 1 | 1 |
| | Molecular Function | GO:0005044~scavenger receptor activity | 4 | 1.315789474 | 0.046606505 | 576823, 373431, 373211, 585092 | 46 | 4 | 183 | 3.97826087 | 0.999517442 | 1 | 1 |
| | Molecular Function | GO:0043167~ion binding | 12 | 3.947368421 | 0.060239568 | 591943, 575697, 100891893, 373360, 577570, 757121, 585092, 580652, 105440441, 592971, 592585, | 46 | 29 | 183 | 1.646176912 | 0.999951834 | 1 | 1 |
| | Molecular Function | GO:0005509~calcium ion binding | 5 | 1.644736842 | 0.063060918 | 591943, 373360, 757121, 105440441, 592971 | 46 | 7 | 183 | 2.841614907 | 0.999970227 | 1 | 1 |
| | Molecular Function | GO:0003824~catalytic activity | 27 | 8.881578947 | 0.067218842 | 591159, 577570, 585092, 578540, 577198, 592585, 574714, 585404, 373360, 580652, 586832, 588773 | 46 | 86 | 183 | 1.248988878 | 0.999985386 | 1 | 1 |
| | Molecular Function | GO:0016787~hydrolase activity | 14 | 4.605263158 | 0.07561082 | 591159, 577570, 757121, 578540, 581938, 592585, 580868, 576823, 574714, 373360, 583337, 580652 | 46 | 37 | 183 | 1.505287897 | 0.999996558 | 1 | 1 |
| | Molecular Function | GO:0043169~cation binding | 11 | 3.618421053 | 0.086149331 | 591943, 575697, 100891893, 373360, 577570, 757121, 585092, 580652, 105440441, 592971, 592585 | 46 | 27 | 183 | 1.620772947 | 0.99999945 | 1 | 1 |
| | Molecular Function | GO:0046872~metal ion binding | 11 | 3.618421053 | 0.086149331 | 591943, 575697, 100891893, 373360, 577570, 757121, 585092, 580652, 105440441, 592971, 592585 | 46 | 27 | 183 | 1.620772947 | 0.99999945 | 1 | 1 |
| | Molecular Function | GO:0016788~hydrolase activity, acting on ester bonds | 4 | 1.315789474 | 0.096329578 | 591159, 757121, 580652, 580300 | 46 | 5 | 183 | 3.182608696 | 0.999999908 | 1 | 1 |
| | Molecular Function | GO:0008237~metallopeptidase activity | 4 | 1.315789474 | 0.096329578 | 373360, 577570, 588773, 592585 | 46 | 5 | 183 | 3.182608696 | 0.999999908 | 1 | 1 |
| 2 | Biological Process | GO:1901566~organonitrogen compound biosynthetic proce | 23 | 7.324840764 | 9.44E-04 | 593029, 593744, 575080, 591935, 588945, 577040, 762952, 591341, 593861, 577020, 591498, 590740 | 58 | 38 | 162 | 1.690562613 | 0.378755347 | 0.475805574 | 0.475805574 |
| | Biological Process | GO:0044271~cellular nitrogen compound biosynthetic proc | 26 | 8.280254777 | 0.003317447 | 752563, 588945, 591935, 577255, 591498, 590740, 574895, 575225, 584839, 581865, 100892475, 373 | 58 | 48 | 162 | 1.512931034 | 0.812649242 | 0.835996592 | 0.835996592 |
| | Biological Process | GO:1901564~organonitrogen compound metabolic process | 23 | 7.324840764 | 0.00631878 | 593029, 593744, 575080, 591935, 588945, 577040, 762952, 591341, 593861, 577020, 591498, 590740 | 58 | 42 | 162 | 1.52955665 | 0.959024032 | 1 | 1 |
| | Biological Process | GO:0044249~cellular biosynthetic process | 28 | 8.917197452 | 0.013615712 | 752563, 754268, 588945, 591935, 577255, 591498, 590740, 574895, 575225, 584839, 581865, 100892 | 58 | 57 | 162 | 1.372050817 | 0.999001708 | 1 | 1 |
| | Biological Process | GO:0043043~peptide biosynthetic process | 16 | 5.095541401 | 0.014924636 | 593029, 593744, 575080, 577040, 591341, 593861, 591498, 590740, 373469, 577852, 574895, 575225 | 58 | 27 | 162 | 1.655172414 | 0.999488781 | 1 | 1 |
| | Biological Process | GO:0006412~translation | 16 | 5.095541401 | 0.014924636 | 593029, 593744, 575080, 577040, 591341, 593861, 591498, 590740, 373469, 577852, 574895, 575225 | 58 | 27 | 162 | 1.655172414 | 0.999488781 | 1 | 1 |
| | Biological Process | GO:0043604~amide biosynthetic process | 16 | 5.095541401 | 0.014924636 | 593029, 593744, 575080, 577040, 591341, 593861, 591498, 590740, 373469, 577852, 574895, 575225 | 58 | 27 | 162 | 1.655172414 | 0.999488781 | 1 | 1 |
| | Biological Process | GO:0009058~biosynthetic process | 28 | 8.917197452 | 0.01872805 | 752563, 754268, 588945, 591935, 577255, 591498, 590740, 574895, 575225, 584839, 581865, 100892 | 58 | 58 | 162 | 1.348394768 | 0.999927247 | 1 | 1 |
| | Biological Process | GO:1901576~organic substance biosynthetic process | 28 | 8.917197452 | 0.01872805 | 752563, 754268, 588945, 591935, 577255, 591498, 590740, 574895, 575225, 584839, 581865, 100892 | 58 | 58 | 162 | 1.348394768 | 0.999927247 | 1 | 1 |
| | Biological Process | GO:0006518~peptide metabolic process | 16 | 5.095541401 | 0.034271009 | 593029, 593744, 575080, 577040, 591341, 593861, 591498, 590740, 373469, 577852, 574895, 575225 | 58 | 29 | 162 | 1.541022592 | 0.999999977 | 1 | 1 |
| | Biological Process | GO:0043603~cellular amide metabolic process | 16 | 5.095541401 | 0.049078681 | 593029, 593744, 575080, 577040, 591341, 593861, 591498, 590740, 373469, 577852, 574895, 575225 | 58 | 30 | 162 | 1.489655172 | 1 | 1 | 1 |
| | Biological Process | GO:0034645~cellular macromolecule biosynthetic process | 21 | 6.687898089 | 0.0528505 | 591688, 752563, 593029, 754268, 593744, 575080, 577040, 591341, 593861, 577255, 591498, 590740 | 58 | 43 | 162 | 1.364073777 | 1 | 1 | 1 |
| | Biological Process | GO:0009059~macromolecule biosynthetic process | 21 | 6.687898089 | 0.0528505 | 591688, 752563, 593029, 754268, 593744, 575080, 577040, 591341, 593861, 577255, 591498, 590740 | 58 | 43 | 162 | 1.364073777 | 1 | 1 | 1 |
| | Biological Process | GO:0034641~cellular nitrogen compound metabolic proces | 32 | 10.1910828 | 0.070295795 | 592405, 752563, 754268, 579591, 588945, 591935, 577255, 591498, 590740, 582919, 574895, 575225 | 58 | 74 | 162 | 1.207828518 | 1 | 1 | 1 |
| | Biological Process | GO:0006807~nitrogen compound metabolic process | 33 | 10.50955414 | 0.073219103 | 592405, 752563, 754268, 579591, 588945, 591935, 577255, 591498, 590740, 582919, 574895, 575225 | 58 | 77 | 162 | 1.197044335 | 1 | 1 | 1 |
| | Cellular Component | GO:0005840~ribosome | 14 | 4.458598726 | 2.43E-04 | 593029, 593744, 577040, 580091, 591341, 593861, 591498, 590740, 373469, 577852, 574895, 584839 | 59 | 18 | 183 | 2.412429379 | 0.033490967 | 0.03406049 | 0.032357465 |
| | Cellular Component | GO:0044391~ribosomal subunit | 10 | 3.184713376 | 0.001873756 | 577852, 593029, 581865, 593744, 577040, 591341, 593861, 587414, 591498, 590740 | 59 | 12 | 183 | 2.584745763 | 0.230928925 | 0.131162907 | 0.124604762 |
| | Cellular Component | GO:0005829~cytosol | 10 | 3.184713376 | 0.004539866 | 577852, 581865, 593744, 588945, 577040, 580091, 591341, 577020, 593861, 590740 | 59 | 13 | 183 | 2.385919166 | 0.471138308 | 0.139107249 | 0.132151887 |
| | Cellular Component | GO:0022626~cytosolic ribosome | 8 | 2.547770701 | 0.004968116 | 577852, 581865, 593744, 577040, 580091, 591341, 593861, 590740 | 59 | 9 | 183 | 2.757062147 | 0.502057038 | 0.139107249 | 0.132151887 |
| | Cellular Component | GO:0044445~cytosolic part | 8 | 2.547770701 | 0.004968116 | 577852, 581865, 593744, 577040, 580091, 591341, 593861, 590740 | 59 | 9 | 183 | 2.757062147 | 0.502057038 | 0.139107249 | 0.132151887 |
| | Cellular Component | GO:0030529~intracellular ribonucleoprotein complex | 18 | 5.732484076 | 0.010999297 | 593029, 587044, 593744, 575080, 577040, 580091, 591341, 593861, 591498, 590740, 373469, 582919 | 59 | 34 | 183 | 1.642073779 | 0.787419209 | 0.185105101 | 0.175849846 |
| | Cellular Component | GO:1990904~ribonucleoprotein complex | 18 | 5.732484076 | 0.010999297 | 593029, 587044, 593744, 575080, 577040, 580091, 591341, 593861, 591498, 590740, 373469, 582919 | 59 | 34 | 183 | 1.642073779 | 0.787419209 | 0.185105101 | 0.175849846 |
| | Cellular Component | GO:0043228~non-membrane-bounded organelle | 23 | 7.324840764 | 0.012551192 | 591688, 593029, 587044, 581030, 593744, 577040, 580091, 373275, 591341, 593861, 577255, 591498 | 59 | 48 | 183 | 1.486228814 | 0.829375031 | 0.185105101 | 0.175849846 |
| | Cellular Component | GO:0043232~intracellular non-membrane-bounded organel | 23 | 7.324840764 | 0.012551192 | 591688, 593029, 587044, 581030, 593744, 577040, 580091, 373275, 591341, 593861, 577255, 591498 | 59 | 48 | 183 | 1.486228814 | 0.829375031 | 0.185105101 | 0.175849846 |
| | Cellular Component | GO:0015934~large ribosomal subunit | 7 | 2.229299363 | 0.013221793 | 577852, 593029, 581865, 591341, 593861, 587414, 591498 | 59 | 8 | 183 | 2.713983051 | 0.844855302 | 0.185105101 | 0.175849846 |
| | Cellular Component | GO:0044444~cytoplasmic part | 24 | 7.643312102 | 0.018000281 | 593029, 593744, 575080, 588945, 588418, 577040, 580091, 591341, 593861, 577020, 373418, 591498 | 59 | 52 | 183 | 1.431551499 | 0.921369521 | 0.229094479 | 0.217639756 |
| | Cellular Component | GO:0005737~cytoplasm | 30 | 9.554140127 | 0.025748533 | 588945, 373275, 373418, 591498, 590740, 574660, 574895, 575225, 584839, 581865, 575808, 593029 | 59 | 71 | 183 | 1.310575316 | 0.974061219 | 0.300399556 | 0.285379578 |
| | Cellular Component | GO:0005622~intracellular | 47 | 14.96815287 | 0.064294176 | 592405, 752563, 754268, 581030, 579591, 588945, 373275, 373418, 763814, 577255, 591498, 590740 | 59 | 130 | 183 | 1.121382008 | 0.999908902 | 0.64317898 | 0.611020031 |
| | Cellular Component | GO:0005623~cell | 51 | 16.24203822 | 0.064317898 | 373418, 591498, 591178, 590165, 584839, 100892475, 373382, 752213, 575808, 591688, 593029, 587 | 59 | 144 | 183 | 1.098516949 | 0.999909225 | 0.64317898 | 0.611020031 |
| | Cellular Component | GO:0044424~intracellular part | 45 | 14.33121019 | 0.075363329 | 592405, 752563, 754268, 581030, 579591, 588945, 373275, 373418, 577255, 591498, 590740, 574660 | 59 | 124 | 183 | 1.12561509 | 0.999982783 | 0.703391071 | 0.668221517 |
| | Cellular Component | GO:0032991~macromolecular complex | 26 | 8.280254777 | 0.080766921 | 581030, 577255, 591498, 590740, 574660, 582919, 574895, 584839, 581865, 100892475, 373382, 752 | 59 | 64 | 183 | 1.260063559 | 0.999992421 | 0.706710556 | 0.671375028 |
| | Cellular Component | GO:0022625~cytosolic large ribosomal subunit | 4 | 1.27388535 | 0.094691349 | 577852, 581865, 591341, 593861 | 59 | 4 | 183 | 3.101694915 | 0.999999106 | 0.779811106 | 0.740820551 |
| | Molecular Function | GO:0005198~structural molecule activity | 17 | 5.414012739 | 3.17E-05 | 593029, 593744, 577040, 373275, 591341, 593861, 591498, 590740, 373469, 577852, 574895, 575225 | 63 | 21 | 183 | 2.351473923 | 0.005981249 | 0.005999114 | 0.005999114 |
| | Molecular Function | GO:0003735~structural constituent of ribosome | 15 | 4.777070064 | 7.91E-05 | 593029, 593744, 577040, 591341, 593861, 591498, 590740, 373469, 577852, 574895, 575225, 584839 | 63 | 18 | 183 | 2.420634921 | 0.014841786 | 0.007476218 | 0.007476218 |
| | Biological Process | GO:0071840~cellular component organization or biogenesis | 26 | 10.6122449 | 0.008717447 | 587575, 589835, 585590, 592267, 575126, 574998, 105438529, 577928, 589486, 582011, 580493, 588 | 61 | 48 | 162 | 1.43852459 | 0.994677905 | 1 | 1 |
| | Biological Process | GO:0044260~cellular macromolecule metabolic process | 37 | 15.10204082 | 0.01136628 | 587575, 590942, 586283, 590968, 575790, 592267, 593496, 578762, 578317, 575203, 580958, 575347 | 61 | 77 | 162 | 1.276133702 | 0.998925564 | 1 | 1 |
| | Biological Process | GO:0043170~macromolecule metabolic process | 37 | 15.10204082 | 0.01136628 | 587575, 590942, 586283, 590968, 575790, 592267, 593496, 578762, 578317, 575203, 580958, 575347 | 61 | 77 | 162 | 1.276133702 | 0.998925564 | 1 | 1 |
| | Biological Process | GO:0016070~RNA metabolic process | 18 | 7.346938776 | 0.020049679 | 588854, 752151, 577044, 594039, 575790, 373348, 592267, 593496, 578317, 577954, 575632, 580958 | 61 | 31 | 162 | 1.542041248 | 0.999994504 | 1 | 1 |
| | Biological Process | GO:0006508~proteolysis | 8 | 3.265306122 | 0.031373568 | 590942, 591601, 575334, 575203, 575347, 100889149, 585921, 578762 | 61 | 10 | 162 | 2.124590164 | 0.999999995 | 1 | 1 |
| | Biological Process | GO:0051641~cellular localization | 8 | 3.265306122 | 0.031373568 | 575632, 589835, 585590, 585934, 589195, 577928, 591499, 581274 | 61 | 10 | 162 | 2.124590164 | 0.999999995 | 1 | 1 |
| | Biological Process | GO:0009057~macromolecule catabolic process | 8 | 3.265306122 | 0.031373568 | 590942, 591601, 575334, 575203, 752151, 100889149, 585921, 578762 | 61 | 10 | 162 | 2.124590164 | 0.999999995 | 1 | 1 |
| | Biological Process | GO:0090304~nucleic acid metabolic process | 20 | 8.163265306 | 0.032613223 | 588854, 590968, 752151, 577044, 594039, 575790, 373348, 592267, 593496, 578317, 577954, 575632 | 61 | 37 | 162 | 1.435533895 | 0.999999998 | 1 | 1 |

| | | | | | | | | | | | | | |
|---|---|---|---|---|---|---|---|---|---|---|---|---|---|
| 3 | Biological Process | GO:0006139~nucleobase-containing compound metabolic p | 24 | 9.795918367 | 0.034737512 | 584340, 588854, 590968, 752151, 577044, 594039, 575790, 373348, 592267, 593496, 578317, 574069 | 61 | 47 | 162 | 1.356121381 | 0.999999999 | 1 | 1 |
| | Biological Process | GO:0046483~heterocycle metabolic process | 24 | 9.795918367 | 0.047021722 | 584340, 588854, 590968, 752151, 577044, 594039, 575790, 373348, 592267, 593496, 578317, 574069 | 61 | 48 | 162 | 1.327868852 | 1 | 1 | 1 |
| | Biological Process | GO:0022613~ribonucleoprotein complex biogenesis | 14 | 5.714285714 | 0.050722687 | 587575, 588854, 585590, 577044, 592267, 577954, 575632, 575126, 574998, 105438529, 589486, 582 | 61 | 24 | 162 | 1.549180328 | 1 | 1 | 1 |
| | Biological Process | GO:0016043~cellular component organization | 18 | 7.346938776 | 0.060468956 | 587575, 588415, 589835, 585590, 373348, 373503, 580307, 763690, 580528, 583958, 585934, 589392 | 61 | 34 | 162 | 1.405978785 | 1 | 1 | 1 |
| | Biological Process | GO:0044248~cellular catabolic process | 8 | 3.265306122 | 0.060621815 | 590942, 591601, 575334, 575203, 752151, 100889149, 585921, 578762 | 61 | 11 | 162 | 1.931445604 | 1 | 1 | 1 |
| | Biological Process | GO:0010467~gene expression | 26 | 10.6122449 | 0.060796025 | 587575, 586283, 575790, 593496, 578317, 580958, 574998, 576519, 105438529, 589486, 582011, 580 | 61 | 54 | 162 | 1.278688525 | 1 | 1 | 1 |
| | Biological Process | GO:0006725~cellular aromatic compound metabolic proces | 24 | 9.795918367 | 0.062339684 | 584340, 588854, 590968, 752151, 577044, 594039, 575790, 373348, 592267, 593496, 578317, 574069 | 61 | 49 | 162 | 1.300769488 | 1 | 1 | 1 |
| | Biological Process | GO:1990542~mitochondrial transmembrane transport | 5 | 2.040816327 | 0.06318405 | 589835, 585934, 589195, 577928, 581274 | 61 | 5 | 162 | 2.655737705 | 1 | 1 | 1 |
| | Biological Process | GO:0006839~mitochondrial transport | 5 | 2.040816327 | 0.06318405 | 589835, 585934, 589195, 577928, 581274 | 61 | 5 | 162 | 2.655737705 | 1 | 1 | 1 |
| | Biological Process | GO:0030163~protein catabolic process | 7 | 2.857142857 | 0.064304928 | 590942, 591601, 575334, 575203, 100889149, 585921, 578762 | 61 | 9 | 162 | 2.06557377 | 1 | 1 | 1 |
| | Biological Process | GO:0051649~establishment of localization in cell | 7 | 2.857142857 | 0.064304928 | 575632, 589835, 585590, 585934, 589195, 577928, 581274 | 61 | 9 | 162 | 2.06557377 | 1 | 1 | 1 |
| | Biological Process | GO:0046907~intracellular transport | 7 | 2.857142857 | 0.064304928 | 575632, 589835, 585590, 585934, 589195, 577928, 581274 | 61 | 9 | 162 | 2.06557377 | 1 | 1 | 1 |
| | Biological Process | GO:0044265~cellular macromolecule catabolic process | 7 | 2.857142857 | 0.064304928 | 590942, 591601, 575334, 575203, 752151, 585921, 578762 | 61 | 9 | 162 | 2.06557377 | 1 | 1 | 1 |
| | Biological Process | GO:0042254~ribosome biogenesis | 11 | 4.489795918 | 0.072996416 | 575632, 577954, 588854, 575126, 585590, 574998, 577044, 589486, 580493, 592267, 586103 | 61 | 18 | 162 | 1.62295082 | 1 | 1 | 1 |
| | Biological Process | GO:0006996~organelle organization | 14 | 5.714285714 | 0.073674626 | 588415, 589835, 585590, 373348, 373503, 763690, 580528, 583958, 585934, 589392, 574998, 589195 | 61 | 25 | 162 | 1.487213115 | 1 | 1 | 1 |
| | Biological Process | GO:0044085~cellular component biogenesis | 16 | 6.530612245 | 0.079112304 | 587575, 588854, 585590, 577044, 592267, 373503, 577954, 575632, 580528, 575126, 574998, 105438 | 61 | 30 | 162 | 1.416393443 | 1 | 1 | 1 |
| | Biological Process | GO:1901360~organic cyclic compound metabolic process | 24 | 9.795918367 | 0.081036534 | 584340, 588854, 590968, 752151, 577044, 594039, 575790, 373348, 592267, 593496, 578317, 574069 | 61 | 50 | 162 | 1.274754098 | 1 | 1 | 1 |
| | Biological Process | GO:0006396~RNA processing | 10 | 4.081632653 | 0.081966607 | 575632, 577954, 588854, 580958, 105438529, 577044, 594039, 589486, 580493, 586103 | 61 | 16 | 162 | 1.659836066 | 1 | 1 | 1 |
| | Biological Process | GO:0034641~cellular nitrogen compound metabolic proces | 33 | 13.46938776 | 0.090703903 | 584340, 587575, 586283, 590968, 575790, 592267, 593496, 578317, 580958, 575347, 105443655, 574 | 61 | 74 | 162 | 1.184315463 | 1 | 1 | 1 |
| | Biological Process | GO:0044238~primary metabolic process | 41 | 16.73469388 | 0.095328282 | 584340, 587575, 590942, 586283, 590968, 575790, 592267, 593496, 578762, 578317, 575203, 580958 | 61 | 96 | 162 | 1.134221311 | 1 | 1 | 1 |
| | Cellular Component | GO:0044428~nuclear part | 18 | 7.346938776 | 3.36E-05 | 591601, 752151, 577044, 594039, 373348, 592267, 591574, 578317, 577954, 575632, 575126, 580958 | 67 | 22 | 183 | 2.234735414 | 0.0060654 | 0.006083767 | 0.005848483 |
| | Cellular Component | GO:0031981~nuclear lumen | 14 | 5.714285714 | 0.001168745 | 752151, 577044, 373348, 592267, 591574, 578317, 577954, 575632, 575126, 576519, 583455, 589486 | 67 | 18 | 183 | 2.124378109 | 0.190765557 | 0.091662373 | 0.08811742 |
| | Cellular Component | GO:0043234~protein complex | 24 | 9.795918367 | 0.001519266 | 587575, 590942, 591601, 589835, 586283, 752151, 574071, 594039, 373348, 578762, 578317, 575334 | 67 | 40 | 183 | 1.63880597 | 0.240576906 | 0.091662373 | 0.08811742 |
| | Cellular Component | GO:0032991~macromolecular complex | 33 | 13.46938776 | 0.003446902 | 587575, 590942, 589835, 586283, 592267, 578762, 578317, 580958, 574998, 585921, 576519, 105438 | 67 | 64 | 183 | 1.408348881 | 0.464720854 | 0.124304455 | 0.1194971 |
| | Cellular Component | GO:0031974~membrane-enclosed lumen | 17 | 6.93877551 | 0.00401343 | 588415, 589835, 752151, 577044, 373348, 592267, 591574, 578317, 577954, 575632, 575126, 575347 | 67 | 26 | 183 | 1.785878301 | 0.51707435 | 0.124304455 | 0.1194971 |
| | Cellular Component | GO:0005622~intracellular | 56 | 22.85714286 | 0.00412059 | 590968, 575290, 579451, 591499, 592785, 592267, 591574, 578762, 578317, 575126, 575203, 580958 | 67 | 130 | 183 | 1.176578645 | 0.526388345 | 0.124304455 | 0.1194971 |
| | Cellular Component | GO:0005730~nucleolus | 11 | 4.489795918 | 0.005775559 | 578317, 575632, 577954, 575126, 583455, 577044, 589486, 580493, 592267, 591574, 586103 | 67 | 14 | 183 | 2.146055437 | 0.649504234 | 0.149339452 | 0.143563893 |
| | Cellular Component | GO:0070013~intracellular organelle lumen | 15 | 6.12244898 | 0.008795471 | 588415, 752151, 577044, 373348, 592267, 591574, 578317, 577954, 575632, 575126, 576519, 583455 | 67 | 23 | 183 | 1.781310837 | 0.797906061 | 0.17688669 | 0.170045769 |
| | Cellular Component | GO:0043233~organelle lumen | 15 | 6.12244898 | 0.008795471 | 588415, 752151, 577044, 373348, 592267, 591574, 578317, 577954, 575632, 575126, 576519, 583455 | 67 | 23 | 183 | 1.781310837 | 0.797906061 | 0.17688669 | 0.170045769 |
| | Cellular Component | GO:0005634~nucleus | 24 | 9.795918367 | 0.013239748 | 591601, 588854, 590968, 752151, 578771, 577044, 594039, 579451, 373348, 592267, 593496, 591574 | 67 | 45 | 183 | 1.456716418 | 0.910399978 | 0.203917025 | 0.196030731 |
| | Cellular Component | GO:0043231~intracellular membrane-bounded organelle | 36 | 14.69387755 | 0.013519361 | 589835, 590968, 579451, 592785, 592267, 593496, 591574, 578317, 575126, 575203, 580958, 575347 | 67 | 76 | 183 | 1.293794187 | 0.914880224 | 0.203917025 | 0.196030731 |
| | Cellular Component | GO:0043227~membrane-bounded organelle | 36 | 14.69387755 | 0.013519361 | 589835, 590968, 579451, 592785, 592267, 593496, 591574, 578317, 575126, 575203, 580958, 575347 | 67 | 76 | 183 | 1.293794187 | 0.914880224 | 0.203917025 | 0.196030731 |
| | Cellular Component | GO:0005615~extracellular space | 7 | 2.857142857 | 0.02670786 | 580528, 575203, 373488, 100889149, 373196, 585147, 373503 | 67 | 8 | 183 | 2.389925373 | 0.9925523 | 0.328818067 | 0.316101346 |
| | Cellular Component | GO:0044421~extracellular region part | 7 | 2.857142857 | 0.02670786 | 580528, 575203, 373488, 100889149, 373196, 585147, 373503 | 67 | 8 | 183 | 2.389925373 | 0.9925523 | 0.328818067 | 0.316101346 |
| | Cellular Component | GO:0044424~intracellular part | 52 | 21.2244898 | 0.027250116 | 590968, 579451, 591499, 592785, 592267, 591574, 578762, 578317, 575126, 575203, 580958, 574998 | 67 | 124 | 183 | 1.145402022 | 0.993266902 | 0.328818067 | 0.316101346 |
| | Cellular Component | GO:0005623~cell | 58 | 23.67346939 | 0.040687985 | 590968, 575290, 579451, 591499, 592785, 592267, 591574, 578762, 578317, 575126, 575203, 580958 | 67 | 144 | 183 | 1.100124378 | 0.999457076 | 0.460282831 | 0.442481838 |
| | Cellular Component | GO:0044422~organelle part | 29 | 11.83673469 | 0.047921827 | 589835, 592785, 592267, 591574, 578317, 575126, 580958, 575347, 574998, 576519, 105438529, 577 | 67 | 62 | 183 | 1.277563794 | 0.999862043 | 0.481880589 | 0.463244323 |
| | Cellular Component | GO:0044446~intracellular organelle part | 29 | 11.83673469 | 0.047921827 | 589835, 592785, 592267, 591574, 578317, 575126, 580958, 575347, 574998, 576519, 105438529, 577 | 67 | 62 | 183 | 1.277563794 | 0.999862043 | 0.481880589 | 0.463244323 |
| | Cellular Component | GO:0005576~extracellular region | 7 | 2.857142857 | 0.057043871 | 580528, 575203, 373488, 100889149, 373196, 585147, 373503 | 67 | 9 | 183 | 2.124378109 | 0.999975848 | 0.543417932 | 0.522401769 |
| | Cellular Component | GO:0044464~cell part | 56 | 22.85714286 | 0.071033853 | 590968, 575290, 579451, 591499, 592785, 592267, 591574, 578762, 578317, 575126, 575203, 580958 | 67 | 140 | 183 | 1.092537313 | 0.999998386 | 0.642856373 | 0.617994525 |
| | Molecular Function | GO:0003676~nucleic acid binding | 21 | 8.571428571 | 3.97E-04 | 587575, 586283, 752151, 574071, 589939, 578771, 577044, 594039, 577120, 373348, 593496, 578317 | 55 | 37 | 183 | 1.888452088 | 0.064153823 | 0.066290996 | 0.066290996 |
| | Molecular Function | GO:0003723~RNA binding | 14 | 5.714285714 | 0.002018721 | 587575, 586283, 752151, 574071, 578771, 577044, 594039, 580307, 583958, 100893477, 574998, 105 | 55 | 22 | 183 | 2.117355372 | 0.286424619 | 0.168563229 | 0.168563229 |
| | Molecular Function | GO:1901363~heterocyclic compound binding | 30 | 12.24489796 | 0.004679523 | 587575, 586283, 590968, 589939, 575790, 593496, 577721, 578317, 575126, 575347, 105443655, 574 | 55 | 70 | 183 | 1.425974026 | 0.543110853 | 0.260493427 | 0.260493427 |
| | Molecular Function | GO:0097159~organic cyclic compound binding | 30 | 12.24489796 | 0.006280826 | 587575, 586283, 590968, 589939, 575790, 593496, 577721, 578317, 575126, 575347, 105443655, 574 | 55 | 71 | 183 | 1.405889885 | 0.650833305 | 0.262224501 | 0.262224501 |
| | Molecular Function | GO:0008135~translation factor activity, RNA binding | 7 | 2.857142857 | 0.066618535 | 587575, 580307, 583958, 586283, 574071, 578771, 582011 | 55 | 11 | 183 | 2.117355372 | 0.999990003 | 1 | 1 |
| | Molecular Function | GO:0008565~protein transporter activity | 4 | 1.632653061 | 0.077764695 | 589835, 585934, 589392, 577928 | 55 | 4 | 183 | 3.327272727 | 0.999998655 | 1 | 1 |
| 4 | Biological Process | GO:0007017~microtubule-based process | 3 | 4.109589041 | 0.06469339 | 373241, 575406, 591043 | 10 | 8 | 162 | 6.075 | 0.999919737 | 1 | 1 |
| | Cellular Component | GO:0005875~microtubule associated complex | 3 | 4.109589041 | 0.011451821 | 373241, 575406, 591043 | 13 | 3 | 183 | 14.07692308 | 0.339424825 | 0.412265572 | 0.412265572 |
| | Cellular Component | GO:0016021~integral component of membrane | 7 | 9.589041096 | 0.060588203 | 592207, 579307, 579243, 588706, 585029, 105438010, 594397 | 13 | 48 | 183 | 2.052884615 | 0.89460589 | 0.496871214 | 0.496871214 |
| | Cellular Component | GO:0031224~intrinsic component of membrane | 7 | 9.589041096 | 0.066621097 | 592207, 579307, 579243, 588706, 585029, 105438010, 594397 | 13 | 49 | 183 | 2.010989011 | 0.916422902 | 0.496871214 | 0.496871214 |
| | Cellular Component | GO:0044430~cytoskeletal part | 3 | 4.109589041 | 0.069009891 | 373241, 575406, 591043 | 13 | 7 | 183 | 6.032967033 | 0.923788166 | 0.496871214 | 0.496871214 |
| | Cellular Component | GO:0015630~microtubule cytoskeleton | 3 | 4.109589041 | 0.069009891 | 373241, 575406, 591043 | 13 | 7 | 183 | 6.032967033 | 0.923788166 | 0.496871214 | 0.496871214 |
| | Molecular Function | GO:0003824~catalytic activity | 11 | 15.06849315 | 0.051247748 | 579307, 578700, 587263, 586173, 373241, 586702, 585029, 105438010, 590365, 594397, 574551 | 15 | 86 | 183 | 1.560465116 | 0.987954167 | 1 | 1 |
| | Molecular Function | GO:0016798~hydrolase activity, acting on glycosyl bonds | 3 | 4.109589041 | 0.091617021 | 587263, 586702, 105438010 | 15 | 7 | 183 | 5.228571429 | 0.999687683 | 1 | 1 |
| 5 | Biological Process | GO:0044763~single-organism cellular process | 5 | 19.23076923 | 0.046405712 | 575157, 580863, 373276, 591240, 593362 | 5 | 76 | 162 | 2.131578947 | 0.990039682 | 1 | 1 |
| | Biological Process | GO:0044699~single-organism process | 5 | 19.23076923 | 0.060178132 | 575157, 580863, 373276, 591240, 593362 | 5 | 81 | 162 | 2 | 0.997571053 | 1 | 1 |
